# Cell type-specific epigenomic variation and its association with genotype in the human breast

**DOI:** 10.1101/2025.04.16.648998

**Authors:** Axel Hauduc, Jonathan Steif, Misha Bilenky, Michelle Moksa, Qi Cao, Shengsen Ding, Connie Eaves, Martin Hirst

## Abstract

**Background:** Understanding the interplay between genomic variation and the epigenome is fundamental to the study of development and mechanisms of disease. Previous studies have leveraged population-scale genotype surveys to associate alleles with epigenomic states in heterogenous tissue types. However, epigenomes are inherently cell type-specific, giving rise to unique genome–epigenome interactions that can influence distinct functional states and susceptibility to disease. Moreover, the extent of individual variation in cell type-specific epigenotypes remains poorly understood, posing additional challenges to accurately link genotypes with epigenomic features.

**Results:** We generated comprehensive genomic and epigenomic profiles of four functionally defined human breast epithelial cell types from eight healthy individuals. To quantify inter-individual epigenomic variation, we developed a statistical framework that measures variability in histone modification landscapes across individuals. This analysis revealed substantially greater variation in repressive chromatin marked by H3K27me3 than in active chromatin marked by H3K27ac and H3K4me3. Integrative chromatin state analysis further identified enhancer elements as the principal source of epigenomic divergence between individuals. Stable enhancer states corresponded to high-confidence *cis*-regulatory elements that underpin cell type-specific transcriptional programs, whereas variable enhancer states were enriched for environmentally responsive regulatory circuits. Mapping genetic variants associated with chromatin state variation uncovered extensive cell type-specificity, with nearly 90% of regulatory variants detected in only a single cell type. These associations were strongly enriched within active regulatory chromatin and, when integrated with gene expression, enabled the prioritization of functional regulatory variants. We experimentally validated one such variant, rs75071948, demonstrating allele-specific regulation of *ANXA1* expression using CRISPR/Cas9 genome editing.

**Conclusions:** Our study defines the landscape of normal epigenomic variation across the major human breast epithelial cell types and demonstrates that genome–epigenome interactions are highly cell type-specific. These findings establish cell type as a critical determinant of the functional interpretation of regulatory genetic variation and provide a framework for understanding how inherited genetic variation shapes normal breast biology and disease susceptibility.

## Background

Following the widespread availability of reference human genomes, genome-wide association studies (GWAS) have become an invaluable tool for studying the impact of genetic variation by associating specific alleles with traits and diseases [1–3]. A key finding of GWAS is the significant enrichment of variant–trait associations in non-coding regions of the genome, where more than 90% of these loci were found [4,5]. These variants are enriched within regulatory elements and are thought to act by modulating the affinity of DNA-binding proteins such as transcription factors (TFs). TFs play a critical role in various regulatory processes, including directing the addition of covalent post-translational modifications to DNA-associated histone proteins (histone marks) and, ultimately, gene transcription [6–8]. Thus, through the action of TFs, genetic variants influence transcriptional and functional states. However, the functional consequences of genetic variants emerge from their interactions with the surrounding epigenomic landscape, which varies systematically across both cell types and individuals.

Defining how genetic variation is translated into regulatory function therefore requires parallel characterization of this epigenomic variability.

Epigenomic states are highly cell type-specific, and comprehensive profiling of purified cell populations has established consensus reference maps across diverse tissues and individuals [9–11]. More recently, single-cell epigenomic approaches have refined this view, revealing that functionally defined cell types occupy continuous distributions of epigenomic states within a set of bounds rather than discrete regulatory entities [12–15]. These reference maps have provided the foundation for large-scale studies linking genetic variation to molecular phenotypes through molecular quantitative trait loci (molQTLs) [16–19]. Integrating histone modifications and chromatin accessibility with genetic variation substantially improves the functional interpretation of non-coding variants compared with transcriptome-based analyses alone. Whereas only approximately 43% of genome-wide association study (GWAS) loci colocalize with expression quantitative trait loci (eQTLs), the incorporation of chromatin accessibility and histone acetylation QTLs explains a much larger fraction of disease-associated regulatory variation [20].

A defining feature of molQTLs is their context specificity. Both GWAS loci and molQTLs are strongly enriched within tissue-specific regulatory landscapes [21], while an increasing number of studies demonstrate that many regulatory associations are further restricted to individual cell types [22,23]. In blood, approximately 30% of *cis*-eQTLs are cell type-specific [21,24–30], whereas in the brain this proportion reaches nearly 46% [31,32]. Population-scale single-cell QTL mapping has reinforced this principle: the OneK1K study, profiling 1.27 million peripheral blood mononuclear cells from 982 individuals, found that most *cis*-eQTLs exhibit cell type-specific allelic effects, including dynamic regulatory effects that emerge along differentiation trajectories [33]. Similarly, approximately 40% of histone quantitative trait loci (hQTLs) identified in immune cells are restricted to individual cell types [26]. Despite these advances, nearly all high-resolution molQTL studies have been performed in blood-derived cell types or immortalized cell lines. Consequently, the extent to which genetic variation shapes epigenomic regulation across purified cell types within solid human tissues remains largely unknown, representing a major gap in our understanding of genome–epigenome interactions in normal tissue biology and disease.

Furthermore, beyond variant–epigenome interactions themselves, the landscape of epigenomic variation within which these associations arise remains poorly understood. Histone marks are among the most dynamic epigenomic modifications and are organized into cell type-specific patterns that define cell type function and identity [9,34,35], yet our understanding of histone mark heterogeneity within functionally defined cell types remains limited [36]. The foundational characterization of inter-individual variability in chromatin states was performed in lymphoblastoid cell lines, where enhancer activity was identified as particularly diverse across individuals while gene expression remained comparatively stable, establishing the regulatory enhancer layer as the principal substrate of inter-individual epigenomic variation [37–39]. This pattern was extended to primary immune cell populations through the BLUEPRINT consortium, which profiled approximately 200 donors across three immune cell types and demonstrated that hQTLs are more cell type-specific than eQTLs or methylation QTLs measured at matched loci in the same dataset [40]. Among primary solid tissues, only brain has been characterized at comparable population scale through bulk H3K27ac profiling of several hundred postmortem samples [41] and a recent multi-tissue extension that included heart, lung, and skeletal muscle [42]. Cell type-resolved population epigenomes within solid tissues remain rarer still, restricted to a small number of brain-specific efforts including FANS-sorted neuronal and neuron-depleted H3K4me3 and H3K27ac profiling in 157 prefrontal cortex samples [43] and a microglia-specific chromatin accessibility study in 150 donors [44]. No equivalent cell type-resolved characterization of histone modification stability and variability has been reported in other primary solid tissues.

The human breast provides an ideal system for investigating cell type-resolved epigenomic variation and genome–epigenome interactions, as its major cell populations can be isolated from normal tissue by fluorescence-activated cell sorting (FACS). Here, we generated International Human Epigenome Consortium (IHEC)-compliant reference epigenomes for four purified breast cell types from eight healthy individuals and paired these with matched whole-genome sequences. We quantified inter-individual variation in histone modifications and chromatin states, defined stable and variable regulatory compartments using ChromHMM concordance analysis, and integrated these with transcription factor motif enrichment to identify the regulatory programs underlying epigenomic variation. We further mapped genetic variants associated with chromatin-state variation, which we term *cis* histone binary trait loci (*cis*-hBTLs), and then prioritized through orthogonal evidence and validated in vitro.

Our analyses reveal a two-compartment organization of the breast enhancer epigenome. Stable enhancer states define the core regulatory architecture of each epithelial cell type and are enriched for lineage-specifying transcription factor networks, whereas variable enhancer states are associated with environmentally responsive regulatory programs shared across cell types.

Genome–epigenome associations are highly cell type-specific, with the vast majority of *cis*-hBTLs restricted to a single cell type and preferentially occurring within active regulatory chromatin. Finally, integration with gene expression enabled the prioritization of functional regulatory variants, including experimental validation of an allele-specific regulatory variant at the *ANXA1* locus. Together, these findings define the stable and variable components of the normal human breast epigenome and demonstrate that both cell identity and inter-individual epigenomic variation are fundamental determinants of the functional interpretation of regulatory genetic variation.

## Results

### Epigenomic characterization of breast cell types

We performed whole-genome sequencing (WGS), chromatin immunoprecipitation followed by sequencing (ChIP-seq) targeting histone modifications, and RNA sequencing (RNA-seq) on four functionally characterized breast cell types from eight individuals (**Fig. 1a**). Fresh breast tissue was collected from reduction mammoplasties and, following single cell dissociation, sorted by FACS using EpCAM and CD49f surface markers to define four constituent cell types [45]: luminal cells (LCs), luminal progenitors (LPs), basal cells (BCs), and stromal cells (SCs), with the latter primarily being composed of fibroblasts, as well as adipocytes and tissue-resident immune cells [46] (**Fig. 1a**). Across all individuals, WGS was performed from genomic DNA extracted from one of four cell types per individual, and following sequence alignment [47], single-nucleotide variants (SNVs) were called [48]. On average, 3.8 million SNVs were identified per individual, of which 99.5% were previously annotated in dbSNP [49], and 88% were shared between two or more individuals (Additional file 1: **Fig. S1**). ChIP-seq for six histone modifications (H3K27ac, H3K4me1, H3K4me3, H3K36me3, H3K27me3, and H3K9me3) marking active promoters/enhancers, poised enhancers, active promoters, active gene bodies, facultative heterochromatin, and constitutive heterochromatin, respectively, was performed on each of the 32 unique samples as previously described, following IHEC standards [11,50]. ChIP-seq alignments were adjusted [51,52] to remove reference-bias-sensitive reads identified from variants called in each individual, and peaks were called [53]. We found that reference bias-robust alignment of ChIP-seq reads shifted a small proportion of overall peak coverage across histone marks, with median Jaccard correlation of variant-aware and hg38-aligned peaks within samples being 0.92 (Additional file 1: **Fig. S2**), corresponding to approximately 8% of peak territory being reassigned. Per-mark Jaccard medians ranged from 0.88 for the broader H3K9me3 mark to 0.95 for the sharper H3K27ac and H3K4me3 marks, indicating that broad chromatin domains are more susceptible to alignment-bias artifacts than narrow peaks. RNA-seq reads were processed through JAGuaR [54], and gene transcription across protein-coding genes was quantified [55].

**Fig. 1.**
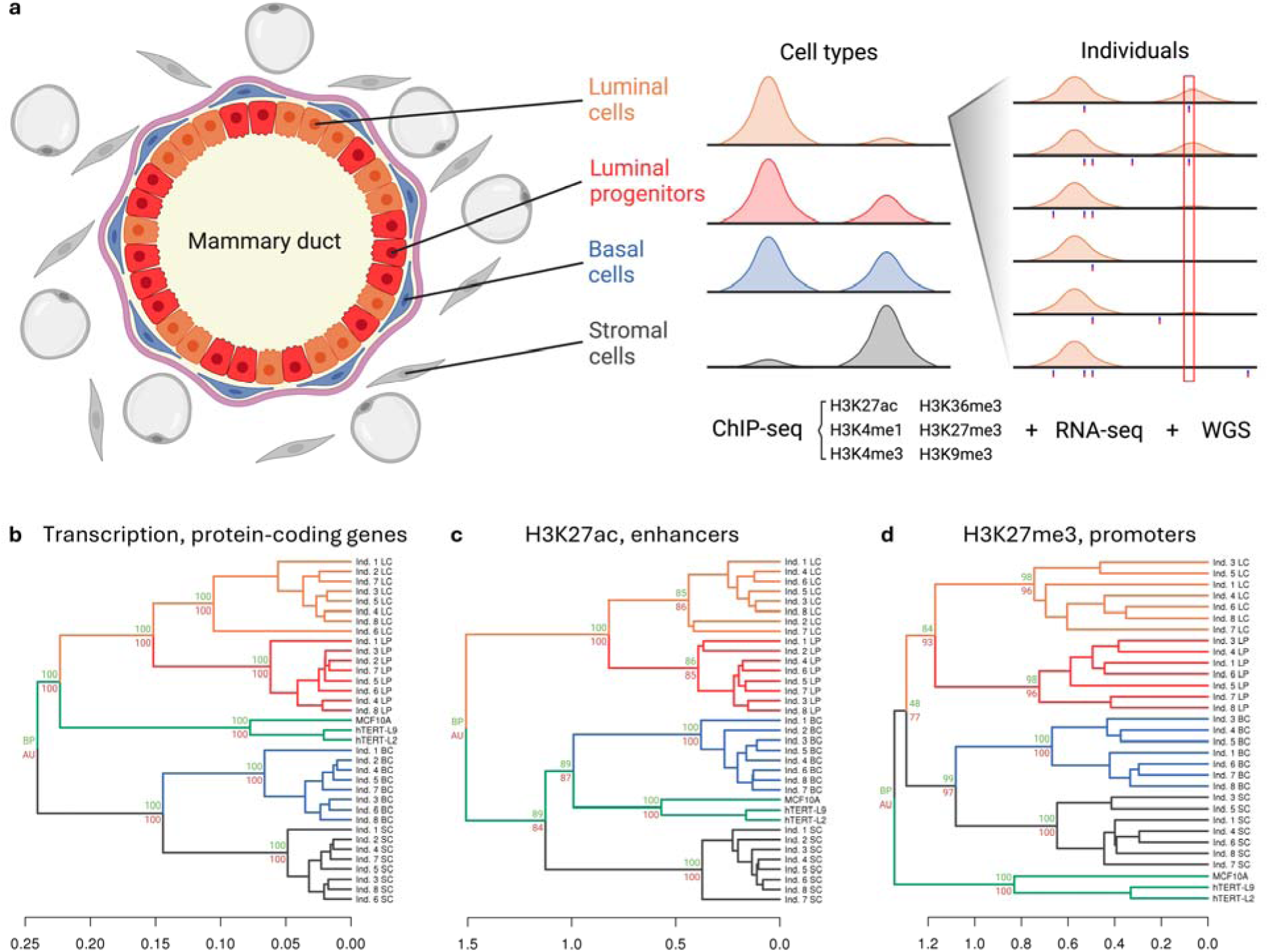
**Study overview and visualization of breast cell type epigenomic relationships**. **a**, Data collection overview. Left, cross-sectional diagram of ductal structure of the breast illustrating the four cell types included in the study; middle, representation of histone modification peaks across cell types; right, representation of genotype and variation in histone modification peaks across different individuals. Dendrograms representing unsupervised clustering of Spearman distances using complete linkage agglomerative clustering on; **b**, protein-coding gene expression; **c**, H3K27ac coverage of the top 1000 variable enhancers [50]; **d**, H3K27me3 coverage of the top 1000 variable promoters. Branches are shaded by cell type (luminal cell (LC), orange; luminal progenitor (LP), red; basal cell (BC), blue; stromal cell (SC), gray; mammary cell lines, green). Bootstrap Probability (BP, green) and Approximately Unbiased (AU, red) values are provided for all major branches.

To define cell type-specific epigenomic states, we performed unsupervised hierarchical clustering on histone mark occupancy in genomic regions relevant to each mark profiled (promoters, enhancers, or gene bodies), and independently on protein-coding gene transcription (**Fig. 1b–1d**, Additional file 1: **Fig. S3**). Cell type clustering based on histone peak landscapes recapitulated known relationships established from cellular characterizations of breast cell types, with LPs and LCs being the most closely related cell types [45,50]. As a comparator, we included matched ChIP-seq datasets of three previously published breast epithelial cell lines that consistently clustered together, away from the primary cell types, reinforcing the importance of studying epigenomic states in primary cell types. Depending on histone mark, the cell lines clustered either closest to basal or luminal cells or formed a distinct branch, potentially reflecting their previously reported mixed basal and luminal characteristics [56], which are distinct from those of primary cells. Active histone marks associated with promoters or enhancers (H3K27ac, H3K4me1, and H3K4me3) proved to be the most effective in distinguishing between cell types, followed by the repressive and transcription-associated marks H3K27me3, H3K9me3, and H3K36me3. As predicted from their functional relationships, and as previously reported [50], LPs and LCs are more similar to each other than BCs as viewed through their epigenomes and transcriptomes, and this relationship is observed consistently across the eight individuals in this study (**Fig. 1b–1d**).

### Epigenomic variation in breast cell types

We next assessed cell type epigenomic variation across the cohort, focusing on H3K27ac, H3K4me1, H3K4me3, and H3K27me3, which are associated with promoter and enhancer regions linked to gene regulation and best classified samples into their respective cell types. In order to do this, we developed an approach to adjust for the contribution of measurement variability within total observed variability. For this, genomes were tiled into 50 base pair (bp) bins and variation in bin occupancy across individuals was calculated for each cell type. Bin values were computed using a Bayesian hierarchical model that accounted for sample-specific effects when estimating the group epigenomic state within each bin (see Methods). The resulting 0 estimates represent likelihoods of histone mark peak presence from zero to one across individuals for each cell type, with zero indicating the absence of an observation in any individual, one indicating stable presence across all individuals, and fractional values indicating different levels of variability (Additional file 1: **Fig. S4**). The degree of variation observed by this measure differed by histone mark, with active marks displaying lower variability compared to H3K27me3, a feature that also varied by genomic context (**Fig. 2a**, Additional file 1: **Fig. S5**). Cell type did not significantly impact mean histone 0 estimates within histone mark and feature categories. We found the most stable histone marks were H3K27ac and H3K4me3 in the context of promoter regions, where median histone 0 estimates were 0.94 and 0.97 across cell types, respectively. H3K27ac and H3K4me3 were also distinct from other histone marks in that their variability significantly differed between promoters and enhancers (*t*-test, *p* < 2.2 × 10^-16^, mean Δ0 = 0.087, 95% CI = 0.085–0.088), with their enhancer regions having median histone 0 estimates of 0.49 and 0.37, respectively. Interestingly, H3K27me3 was significantly more variable than other marks across both promoter and enhancer regions (*t*-test, *p* < 2.2 × 10^-16^, mean Δ0 = 0.132, 95% CI = 0.131– 0.133), with the exception of H3K4me3 at enhancers.

**Fig. 2.**
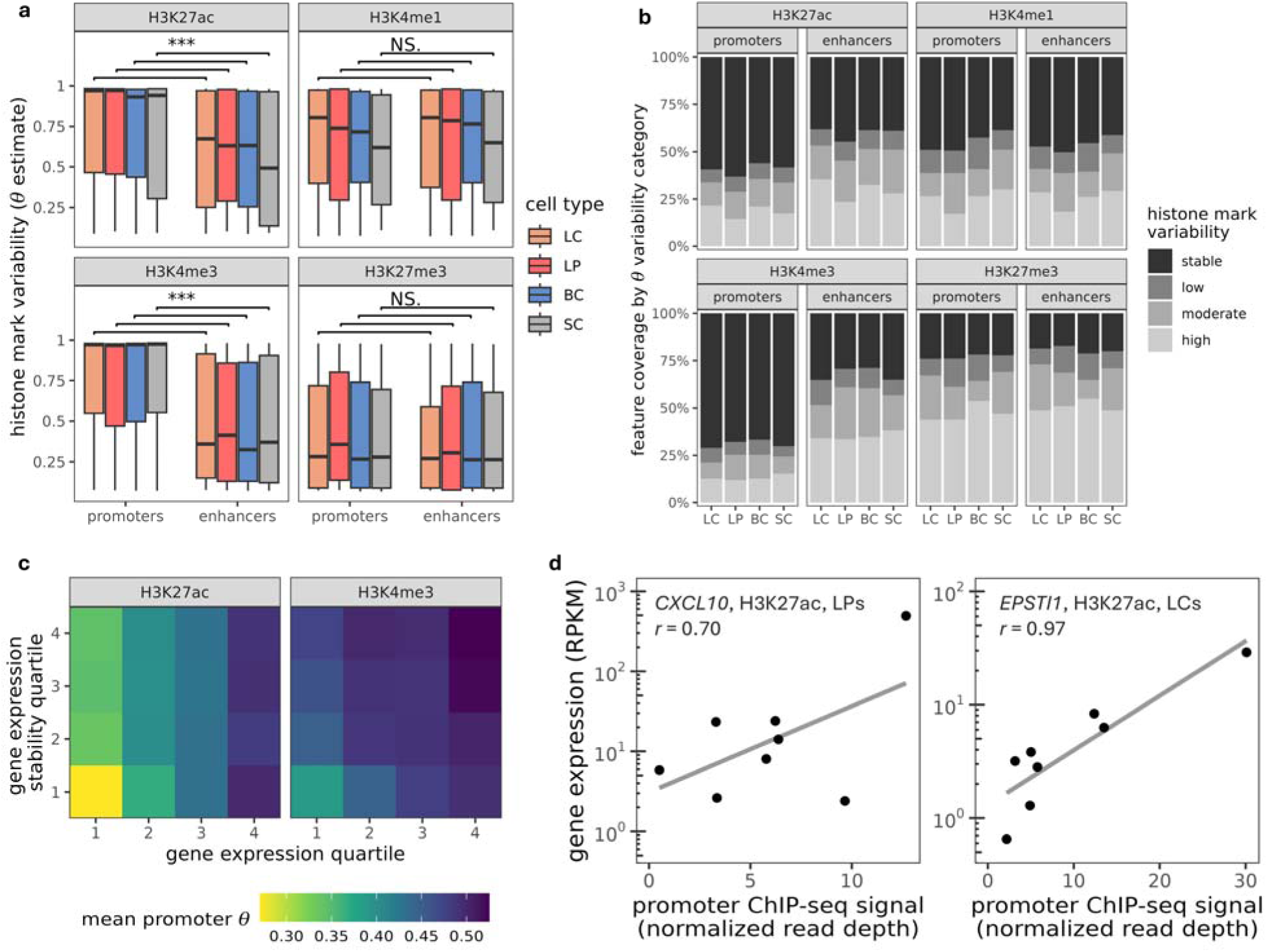
**Histone modification variability in breast cell types**. **a**, Box and whisker plot of Bayesian theta values (histone modification variability estimates) in promoters and enhancers across breast cell types (* for *p* < 0.05, ** for *p* < 0.01, and *** for *p* < 0.001). **b**, Bayesian theta values reported as quartiles excluding unobserved (0) bins in promoters and enhancers across breast cell types as stable (0.75 – 1), low (0.50 – 0.75), moderate (0.25 – 0.50) and high (≤ 0.25). **c**, Heatmap showing the association of mean promoter Bayesian theta with gene expression and variability quartiles. **d**, Correlation of H3K27ac promoter ChIP-seq read depth and gene expression of *CXCL10* (left) in luminal progenitors (LP) and *EPSTI1* in luminal cells (LC) (right).

Previous work has noted that H3K27me3 is highly variable across different cell types and linked with both cell type-specific gene transcription and methylation [57], and we find that in breast cell types, this variability extends to inter-individual variation within a cell type. To better understand the structure of this variability across genomic features, we classified histone 0 bin values into quartiles with variability described in increasing order as stable, low, moderate, and high (**Fig. 2b**). This revealed generally bimodal variability signatures across histone modification and cell type contexts, where most bins were either stable or highly variable, and fewer bins captured intermediate levels of variability across individuals. Although this bimodal structure was common within modifications, the proportions of stable versus variable bins differed significantly between them. For example, H3K27ac and H3K4me3 at promoters were the most stable overall, with an average of 59% and 69% of promoter regions classified as stable, and only 19% and 13% classified as highly variable, respectively. In contrast, H3K27me3 was predominantly classified as variable, with average of 47% promoter and 51% enhancer regions classified as highly variable.

We next sought to characterize the functional implications of histone mark variability by focusing on active, promoter associated marks H3K27ac and H3K4me3. Mean histone 0 at gene promoters was computed, gene expression was quantified across individuals, and genes were subsequently grouped into expression quartiles to calculate mean expression and expression relative standard deviation (RSD) within each quartile (**Fig. 2c**). Promoter histone 0 was only weakly associated with gene expression (Spearman’s p = 0.25 for H3K27ac and p = 0.12 for H3K4me3) and showed little relationship with inter-individual transcriptional variability (Spearman’s p = −0.08 for H3K27ac and p = −0.09 for H3K4me3). We next asked whether loci exhibiting variable promoter 0 also displayed variable transcription. Restricting our analysis to genes with a transcriptional RSD ≥ 100 and a mean RPKM ≥ 1 across individuals identified 468 associations, of which 431 involved promoter- and enhancer-associated active histone marks (Additional file 1: **Fig. S6**). Several had known relations to mammary gland function and disease (e.g., *ADORA1* [58], *ANKRD12* [59], *ANXA1* [60–64], *AZGP1* [65], *CXCL10* [66–69], *CYP24A1* [70,71], *EPSTI1* [72–75], *IGFBP5* [76,77], *MMP7* [50,78–80], *MUCL1* [81,82], *PDGFRA* [83–87], *PI3* [88,89]*, SAA2* [90], *SERPINE1* [91], *SFRP1* [92], and *TPM2* [93]). For example, *CXCL10* exhibited high inter-individual variation in both promoter H3K27ac and gene expression in LPs, with promoter H3K27ac levels strongly correlated with transcript abundance (Pearson’s r = 0.70; **Fig. 2d**, left), a relationship that was not observed in LCs or BCs. As *CXCL10* has been implicated in cell proliferation, invasion, and immune cell infiltration in breast cancer, these findings suggest that its regulatory activity may be modulated in a cell type-specific manner [66]. Likewise, *EPSTI1* displayed substantial variation in promoter H3K27ac and gene expression in LCs (**Fig. 2d**, right). Given its reported role in degradation of the mammary duct basal membrane during breast cancer progression, the concordance between promoter H3K27ac and gene expression suggests cell type-specific regulatory control in luminal cells.

Whereas variable histone marking identifies regulatory elements that differ between individuals, stable histone marking is expected to define the core regulatory architecture of each breast cell type. We therefore asked whether promoters exhibiting stable histone marking were enriched for genes and transcriptional programs underlying shared and cell type-specific functions. To test this, we examined genes associated with stable promoter histone marking, focusing on H3K27ac. Comparison of stable and variable H3K27ac regions using monaLisa [94] showed that variable regions were broadly depleted for TF binding motifs across all cell types relative to stable regions (Additional file 1: **Fig. S7**), consistent with stable regulatory elements harboring the principal transcriptional programs, whereas variable regions may represent more environmentally responsive regulatory activity. Consistent with this model, stably H3K4me3-marked promoters in LPs were enriched for lactation-associated genes, potentially reflecting lineage priming related to parity [95] (Additional file 1: **Fig. S8**). H3K4me3-marked promoters in LCs were enriched for *SERPINA1*-associated functions, whereas stably H3K27ac-marked promoters in LPs were enriched for RAGE receptor binding, both linked to breast cancer biology [96–100]. Stable promoter marking also coincided with cell type-specific expression of numerous genes implicated in mammary development and breast cancer, including *AFF3*, *PTHLH*, *LTF*, *RASAL1*, *PAK5*, *MAMDC2*, and *COL17A1* (Additional file 1: **Fig. S9a–e**) [101–115].

Because stable chromatin states are expected to reflect lineage-defining regulatory programs [116], we next examined TF motif enrichment within stable and highly variable H3K27ac regions using HOMER. Stable regions were enriched for established lineage TFs, including EHF in LPs and TP63 in BCs, consistent with previous studies [50,117], whereas SCs were enriched for PITX1:EBOX motifs previously associated with vertebrate stromal cells [118] (Additional file 1: **Fig. S10**). LCs showed enrichment for AP-1 family motifs, including FRA1, FRA2, FOS, and JUNB. By contrast, highly variable H3K27ac regions exhibited substantially less cell type specificity, with AP-1 family motifs predominating across all cell types (Additional file 1: **Fig. S11**).

### Chromatin state concordance reveals extensive inter-individual epigenomic variation

To determine how inter-individual histone mark variability is organized within higher-order regulatory elements, we characterized chromatin states across all breast cell types using a unified ChromHMM segmentation framework (see Methods). We then quantified chromatin state concordance between individuals using pairwise Jaccard similarity, comparing both matched cell types across individuals and each cell type against the remaining three (**Fig. 3a**). To assess whether these patterns extended beyond the breast epithelium, we performed the same analysis in naïve CD4^+^ and CD8^+^ T cells from the IHEC EpiATLAS (Additional file 1: **Fig. S12**).

**Fig. 3.**
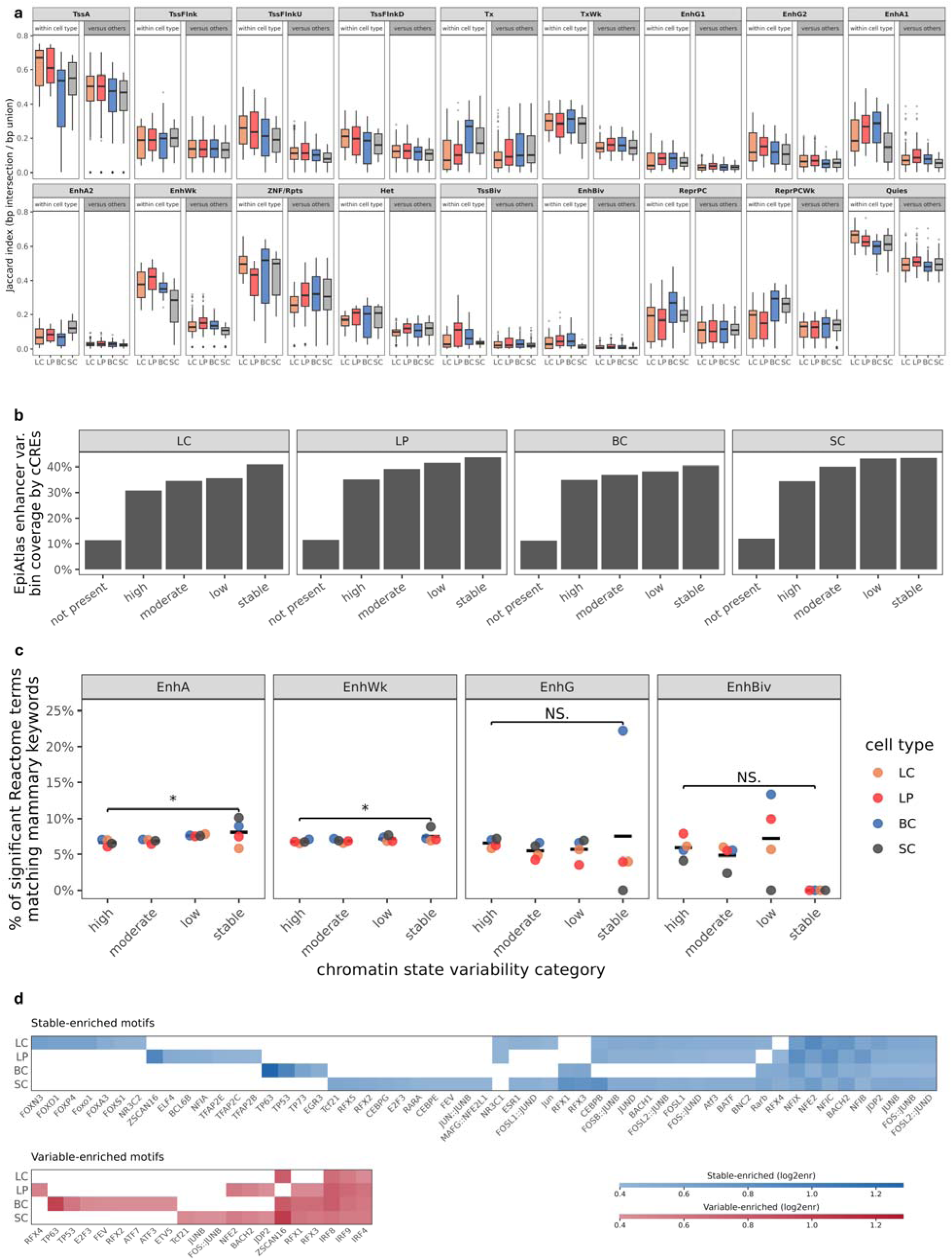
Chromatin state concordance across individuals and functional characterization of stable versus variable enhancer segments. **a**, Boxplots of base-pair-weighted Jaccard index for each of 18 EpiAtlas chromatin states, comparing within-cell type and cell type-versus-other-three pairwise individual comparisons across breast cell types. **b**, Fraction of ChromHMM enhancer segment genomic territory (merged across all enhancer states: EnhA1, EnhA2, EnhG1, EnhG2, EnhWk, EnhBiv) overlapping ENCODE Registry v4 enhancer-class cCREs, stratified by inter-individual variability category (not present: no individuals call an enhancer state; high: ≤ 25%; moderate: 25–50%; low: 50–75%; stable: > 75%). **c**, Percentage of significant GREAT Reactome terms matching a curated mammary relevance keyword set is shown across variability bins, with points representing individual cell types and crossbars indicate the cross-cell type mean. JT tests were applied across the four variability bins with BH correction (* for *p* < 0.05, ** for *p* < 0.01, and *** for *p* < 0.001). **d**, TF motifs enriched in stable versus variable enhancer chromatin across breast cell types. Motifs reaching adjusted *p* < 0.05 and log_2_ enrichment > 0.4 in at least one cell type are shown (55 stable, 21 variable). Motifs are grouped by the set of cell types in which they reach significance and sorted within blocks by mean log_2_ enrichment.

Chromatin state concordance varied markedly across functional chromatin compartments (**Fig. 3b**). Quiescent chromatin and active promoters were the most conserved chromatin states between individuals within each cell type, whereas enhancer states showed substantially lower concordance. Across all enhancer states combined, the median within-cell type Jaccard index was only 0.13, indicating that most enhancer-associated genomic intervals differ between any two individuals despite sharing the same cell identity. Bivalent chromatin exhibited the lowest concordance of all chromatin states, consistent with its dynamic regulatory nature [119].

Despite these differences, chromatin state concordance was significantly higher within matched cell types than between different cell types for all 18 chromatin states (BH-adjusted *p* < 0.001; Additional file 1: **Fig. S13**), demonstrating that cell type remains the dominant determinant of chromatin state organization. Together, these analyses identify enhancer and bivalent chromatin states as the principal source of inter-individual epigenomic variation, whereas promoter and quiescent states constitute a stable regulatory framework that is largely conserved across individuals.

To further characterize the level of chromatin variation at enhancers, we annotated enhancer segments with the proportion of individuals calling any enhancer state and assigned it to one of four variability categories (high variability, > 0 to ≤ 25% of individuals; moderate variability, > 25 to ≤ 50%; low variability, > 50 to ≤ 75%; stable, > 75%), then assessed how each category distributed across cell types. Strikingly, we found that stable enhancers were the most widely shared, with 36.1% of stable enhancer regions shared with enhancers of any stability in all three other cell types, while 14.7% were specific to a single cell type. By contrast, only 17.0% of highly variable enhancer regions were also present across all cell types, and 33.6% present in only one cell type (Additional file 1: **Fig. S14a**). Enhancers that were consistently stable across individuals in one breast cell type were therefore more likely to be shared across other cell types, albeit at lower levels of stability consistency, and, on average, 53.9% of any cell type’s stable enhancer territory was uniquely stable in that cell type. Enhancer state composition further tracked this partition, with the share of enhancer state calls assigned to the active EnhA1 state rising from 7.6% in the most variable category, to 23.7% in the stable category. By contrast, weak (EnhWk) and bivalent (EnhBiv) calls declined (76.6% to 70.8% and 1.6% to 0.2%, respectively; Additional file 1: **Fig. S14b**).

### Stable enhancer states define core mammary regulatory programs

Having established that enhancer states account for most inter-individual epigenomic variation, we next asked whether enhancer stability reflects regulatory function. Stable enhancer positions were more likely to overlap independently annotated enhancer-class candidate *cis*-regulatory elements (cCREs) from the ENCODE Registry [120] than variable enhancers, although fewer than half of stable enhancer intervals overlapped an annotated cCRE (40–44% across breast cell types; **Fig. 3b**, Additional file 1: **Fig. S15**). Variable enhancer positions showed progressively lower cCRE overlap and encompassed substantially more genomic territory (Additional file 1: **Fig. S16**), indicating that much of the variable enhancer compartment remains uncatalogued.

Among overlapping cCREs, enhancer stability was positively associated with regulatory strength, measured by H3K27ac maximum Z-score (Jonckheere–Terpstra trend test, *p* < 0.0001 in all cell types; Additional file 1: **Fig. S17a**), with the strongest relationship observed for active enhancers (Additional file 1: **Fig. S17b**). Enhancer state calls depend on histone mark signal crossing a detection threshold in each individual, so a weakly marked region is called inconsistently between individuals and is also more likely to go undetected in another cell type. Because enhancer stability and regulatory strength covary, the enhancer sharing described above could be generated by detection sensitivity alone rather than by any difference in how broadly enhancers are used. To disambiguate this, we matched enhancers of different variabilities by H3K27ac maximum Z-score, and this showed that within every Z-score decile, stable enhancers remained more widely shared between cell types compared to highly variable enhancers (25.3% versus 11.2% in the lowest decile, 52.6% versus 26.3% in the highest; Additional file 1: **Fig. S18**). Functional enrichment analysis showed that stable active and weak enhancers were progressively enriched for mammary-specific biological pathways, whereas bivalent chromatin became depleted for mammary-relevant functions with increasing stability (**Fig. 3c**, Additional file 1: **Fig. S19**).

Together, these findings support a two-compartment model of the enhancer epigenome. Stable enhancer states define the core regulatory architecture of each breast cell type: they preferentially overlap independently annotated regulatory elements, exhibit stronger regulatory signatures, and are enriched for mammary-specific biological functions. By contrast, variable enhancer states comprise a larger, weaker, and functionally less resolved regulatory compartment that likely captures context-dependent regulatory activity, including environmental responses and epigenomic drift.

### Stable and variable enhancer compartments encode distinct regulatory programs

To define the regulatory logic underlying the two enhancer compartments, we compared transcription factor motif enrichment and associated gene programs between stable and variable enhancer states (see Methods). Stable enhancers were consistently enriched for lineage-defining transcription factors characteristic of each breast cell type (**Fig. 3d**). LCs were enriched for forkhead, ETS, AP-2, and nuclear receptor families, basal cells for TP63/TP53, and stromal cells for TCF21, RFX, and CEBP family motifs, recapitulating established mammary lineage regulators [50]. Across all four cell types, stable enhancers were additionally enriched for NFI family motifs, extending their established role in differentiated mammary epithelium to the stromal compartment [121]. In contrast, variable enhancers displayed remarkably little cell type specificity. The only transcription factor family consistently enriched across all four breast cell types was the interferon regulatory factor (IRF) family, while AP-1 motifs were shared between both compartments, consistent with their established roles in both lineage maintenance and inducible stress responses [122,123]. These findings indicate that stable enhancers encode cell identity programs, whereas variable enhancers preferentially capture environmentally responsive regulatory activity.

Pathway enrichment analysis mirrored this partitioning (Additional file 1: **Fig. S20**; Additional file 1: **Table S5**). Stable enhancers were associated with canonical lineage-specific biological processes, including estrogen signaling in mature LCs, Notch signaling in LPs, TGF-β signaling in BCs, and extracellular matrix organization in SCs. In contrast, variable enhancers were enriched for broadly shared, inducible pathways, including interferon, TP53, nuclear receptor, and immune-response signaling, with relatively little cell type specificity.

Together, these findings provide a regulatory framework for the two-compartment model of the breast enhancer epigenome. The stable enhancer compartment concentrates lineage-defining transcription factors and their associated biological pathways, thereby encoding the core regulatory architecture of each breast cell type. Conversely, the variable enhancer compartment is dominated by broadly shared, environmentally responsive regulatory programs that likely underlie inter-individual epigenomic diversity.

### Genetic variation modulates cell type-specific regulatory architecture

Having established the two-compartment organization of the breast enhancer epigenome, we next investigated the contribution of genetic variation to inter-individual epigenomic diversity by identifying *cis* histone binary trait loci (*cis*-hBTLs) (Additional file 1: **Fig. S21**; see Methods).

Using this approach, we identified 245,435 *cis*-hBTLs across the four breast cell types (Additional file 7: **Table S6**). Nearly 90% were detected in only a single cell type, with fewer than 0.1% shared across all four cell types (**Fig. 4a**; Additional file 1: **Fig. S22;** Additional file 8: **Table S7**), demonstrating that genetic effects on histone deposition are highly cell type-specific. We further assessed whether the degree of cell type-specificity of *cis*-hBTLs varied by histone mark, finding that H3K27ac, H3K4me1, and H3K27me3 exhibited the greatest cell type-specificity.

**Fig. 4.**
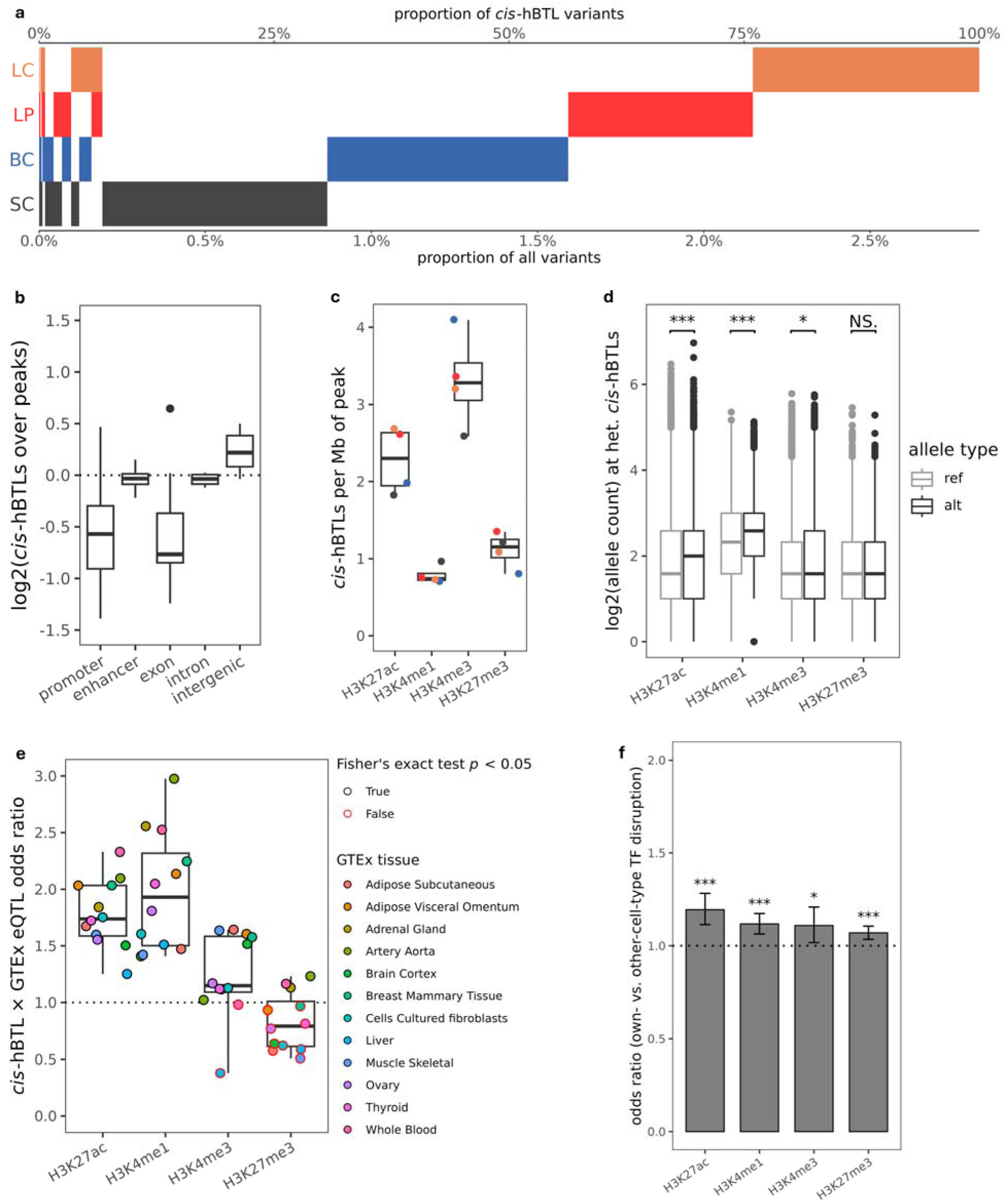
Association between genomic and epigenomic variation across breast cell types. **a**, Breakdown of *cis*-hBTLs by cell type (y axis) and percentage of *cis*-hBTLs (x upper axis) and total variants (x lower axis). **b**, Enrichment of sequence features within *cis-*hBTLs relative to associated histone modification peak sets. **c**, Box and whisker plot of *cis*-hBTL density normalized by overall peak coverage for each cell type (encoded by color) and histone modification. **d,** Allelic count imbalance of heterozygous *cis*-hBTLs within associated histone mark peaks (* for *p* < 0.05, ** for *p* < 0.01, and *** for *p* < 0.001). **e**, Enrichment of *cis-*hBTL sets within GTEx tissue-specific eQTL defined as the odds ratio of variants belonging to both the *cis*-hBTL set and the GTEx eQTL set, with the significance of each comparison by Fisher’s exact test represented as outline color. **f**, Odds ratios comparing the enrichment of *cis*-hBTLs from each breast cell type for disruption of JASPAR2024 CORE vertebrate TF binding motifs corresponding to the leading cell type-discriminating TFs identified in Pellacani et al. 2016 [50] Fig. 7B (* for *p* < 0.05, ** for *p* < 0.01, and *** for *p* < 0.001).

Consistent with a regulatory mechanism, *cis*-hBTLs were enriched within active regulatory chromatin, particularly H3K27ac and H3K4me3, showed significant allelic imbalance in ChIP-seq reads, and were enriched for GTEx eQTLs (**Fig. 4b–e**; Additional file 1: **Fig. S23–S25**). Motif disruption analysis further showed that active mark *cis*-hBTLs preferentially disrupted binding sites for transcription factors defining the same breast cell type in which the *cis*-hBTL was observed (**Fig. 4f**), whereas H3K36me3 and constitutive heterochromatin showed little or no such enrichment (Additional file 1: **Fig. S26**).

We next asked whether the localization of these variants is linked to the landscape of cross-cohort enhancer stability. Active mark *cis*-hBTLs were enriched within the enhancer regions of their own cell type across all of the previously-defined variability categories (log_2_ enrichment 2.25–2.80 in LCs, 2.61–3.04 in LPs, 2.47–2.82 in BCs, and 2.70–2.98 in SCs; Additional file 1: **Fig. S27**), demonstrating that genetic association with histone marks is distributed throughout the histone variability landscape. Averaged across cell types, same-cell type *cis*-hBTL enrichment was highest within the most variable enhancers, at 2.89 log_2_ units against 2.51, 2.67 and 2.64 in the moderately variable, low-variability and stable categories respectively. By contrast, cross-cell type enrichment rose monotonically across enhancer variabilities, from 1.58 log_2_ units in the highly variable category through to 1.90 in the stable category. *Cis*-hBTLs are therefore most sharply restricted to same-cell type enhancers where those enhancers are least consistently called, with the rising cross-cell type enrichment likely stemming from the increased sharing of stable enhancer regions between cell types that was previously observed. However, despite their highly cell type-specific genomic localization, *cis*-hBTLs were associated with broadly shared biological pathways rather than lineage-specific gene programs (Additional file 1: **Fig. S28–S29**). These findings indicate that genetic variation acts predominantly by modulating regulatory activity within cell type-specific enhancer architectures rather than fundamentally changing them.

To identify candidate causal variants for functional validation, we prioritized *cis*-hBTLs associated with significant changes in nearby gene expression. This yielded 1,907 expression-linked *cis*-hBTLs, from which nine high-confidence candidates were selected following manual inspection (Additional file 9: **Table S8**). Consistent with purifying selection, variants associated with the largest transcriptional effects were typically of low population frequency, with 73% exhibiting a gnomAD allele frequency below 10% (Additional file 1: **Fig. S30**).

### Functional validation identifies a causal *cis*-hBTL at the *ANXA1* locus

To establish whether *cis*-hBTLs directly influence chromatin state and transcription, we prioritized expression-associated variants for functional validation (see Methods). Among the manually curated candidates, heterozygous rs75071948 in Individual 7, located within the first intron of *ANXA1*, was associated with the presence of a de novo H3K4me3 peak and increased *ANXA1* expression in LCs (**Fig. 5a–b**). The variant lies within an ENCODE candidate *cis*-regulatory element and was predicted to alter the binding affinity of multiple transcription factors expressed in breast epithelial cells (Additional file 1: **Fig. S31**). Consistent with the primary tissue observations, breast cell lines carrying the alternate allele exhibited both increased H3K4me3 deposition and higher *ANXA1* expression, together with allelic imbalance in intronic RNA reads (Additional file 1: **Fig. S32**).

**Fig. 5.**
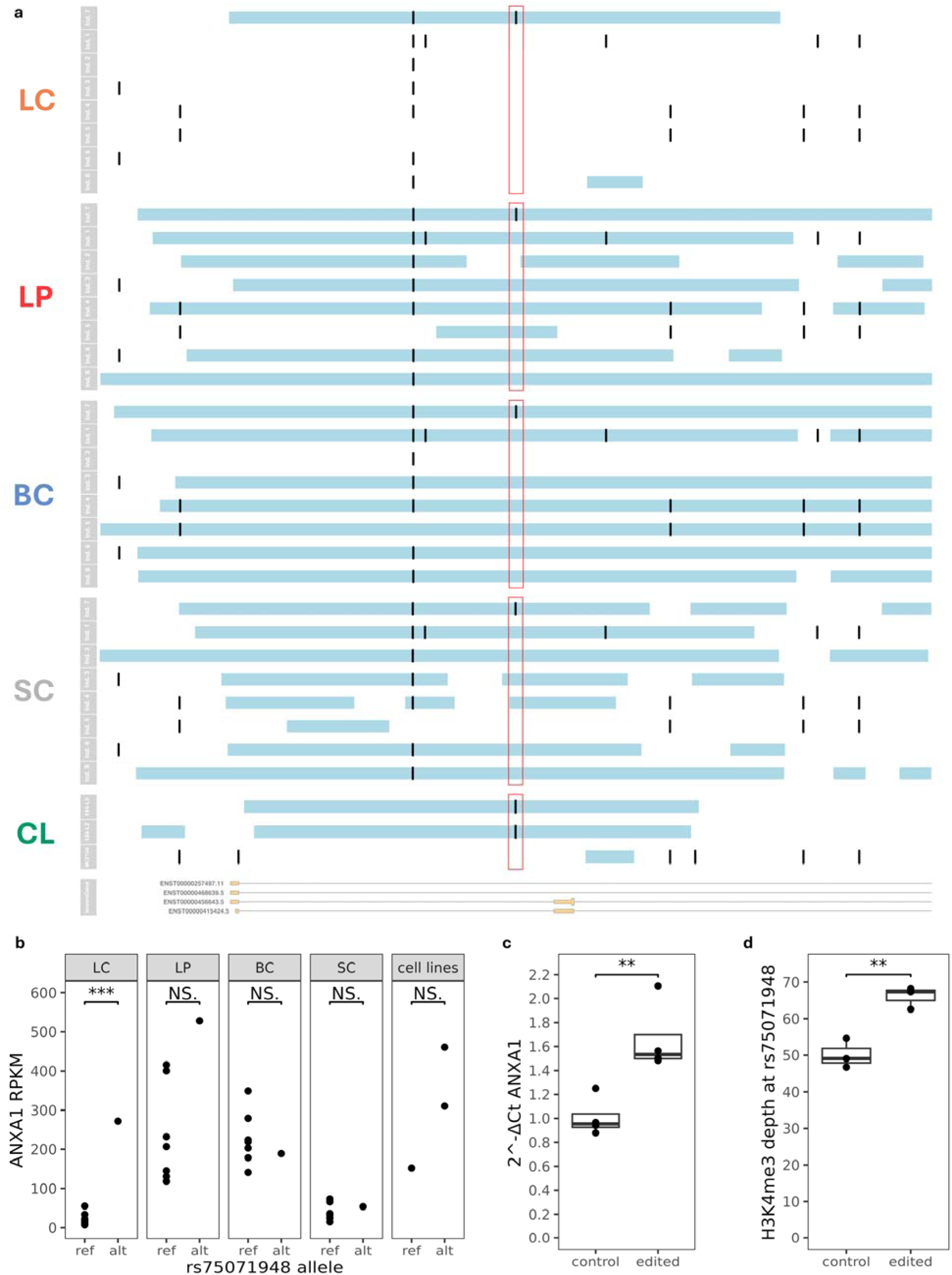
**Association between rs75071948, histone mark peaks and *ANXA1* expression by cell type**. **a,** MACS2 H3K4me3 peak calls (horizontal blue bars) and single-nucleotide variants (black vertical bars) along the first intron of *ANXA1* in chromosome 9, centered on rs75071948 ± 3 kb. **b,** *ANXA1* expression (RPKM) by rs75071948 genotype across all cell types (* for *p* < 0.05, ** for *p* < 0.01, and *** for *p* < 0.001). **c**, Normalized qPCR of *ANXA1* expression in control (parental) MCF10A and rs75071948-edited MCF10A (* for *p* < 0.05, ** for *p* < 0.01, and *** for *p* < 0.001). **d,** Normalized H3K4me3 read depth at rs75071948 in control (parental) MCF10A and rs75071948-edited MCF10A (* for *p* < 0.05, ** for *p* < 0.01, and *** for *p* < 0.001).

We therefore introduced the rs75071948 alternate allele into MCF10A cells, which carry the reference allele, by CRISPR-mediated genome editing (see Methods). Since rs75071948 is a single T > G transversion, we designed a CRISPR/Cas9 guide RNA targeting near rs75071948, as well as an ssODN repair template carrying the variant allele of interest to facilitate HDR [124–126] (Additional file 1: **Fig. S33**). Following sequence validation and clonal derivation, *ANXA1* RT-qPCR was performed on edited MCF10A clonal lines carrying the rs75071948-G alternate genotype along with parental controls and a significant increase in expression was observed as predicted by our model (**Fig. 5c**, Welch’s two-sample *t-*test on C_t_ values *p* = 4.6 × 10^-3^). ChIP-seq for H3K4me3 was then performed on edited and parental MCF10A clonal lines and H3K4me3 normalized read density was found to be significantly increased in association with rs75071948-G (**Fig. 5d**, Welch’s two-sample *t*-test *p* = 6.9 × 10^-3^). These findings establish rs75071948 as a functional variant that modulates *ANXA1* expression and chromatin state, highlighting its potential regulatory role and relevance for further mechanistic and disease-related studies.

## Discussion

Our study identifies a previously unrecognized two-compartment organization of the human breast enhancer epigenome. Stable enhancer states define the core regulatory architecture that underpins mammary cell identity, whereas a complementary variable enhancer compartment captures inter-individual epigenomic diversity arising from genetic, environmental, and stochastic influences. By integrating histone modification landscapes, chromatin state segmentation, transcription factor binding logic, and genotype, our findings provide a unified framework for understanding how epigenomic variation is organized within normal human tissues.

Although the overall epigenomic landscapes of the four breast cell types closely recapitulated their established developmental relationships [45,50,127,128], substantial inter-individual variation was evident within each cell type. Active promoter states were generally highly conserved, whereas active distal regulatory states and H3K27me3 states displayed considerably greater variability, suggesting that enhancers and repressive chromatin are more permissive to inter-individual divergence. Extending these observations to chromatin states demonstrated that enhancer chromatin constitutes the principal locus of inter-individual epigenomic variation. Promoter and quiescent chromatin showed the highest concordance between individuals, while enhancer and bivalent states exhibited markedly lower concordance. Together, these findings indicate that cell identity is maintained through a remarkably stable promoter architecture, whereas enhancer landscapes provide the principal substrate for individual variation.

Multiple independent analyses converged on the same biological model. Stable enhancers preferentially overlapped independently annotated ENCODE candidate *cis*-regulatory elements, exhibited stronger regulatory signatures, and were enriched for mammary-specific biological pathways and lineage-defining transcription factor motifs. Conversely, variable enhancers showed reduced overlap with annotated regulatory elements, were enriched for interferon regulatory factor and AP-1 family motifs, and were associated with broadly shared inflammatory and environmentally responsive pathways. Rather than representing poorly defined regulatory elements, these regions appear to constitute a dynamic regulatory compartment through which developmental identity is integrated with lifetime environmental exposures.

Somewhat counterintuitively, stable enhancer positions were the most widely shared between the four breast cell types, and high variability positions the most cell type-specific. This points to cell type-specificity of enhancers being a measure of the consistency with which a given cell type engages a specific enhancer region from a broader set of candidates, together with the TF grammar and biological programs encoded within it. The fact that this relationship survived matching on regulatory strength indicates genuine differences in how broadly enhancer regions are engaged, rather than the greater detectability of strongly marked regions in any one cell type. I observed that while 85.3% of a cell type’s stable enhancer space was also called as an enhancer in at least one other cell type, only 46.1% was stable in any other cell type, leaving 53.9% uniquely stable in that cell type. Cell type-specific motif enrichments observed in stable enhancers therefore derive from the majority of stable enhancer positions unique to each cell type. Consistent with this partition, the one family enriched in the stable compartment of all four cell types was NFI, whereas the lineage-defining families were each confined to the cell type whose identity they specify. This relationship is consistent with observations in other systems. In brain, heart, muscle and lung, regulatory elements shared by all four tissues were enriched for promoter annotations, while tissue-restricted elements were enriched for enhancers [42], matching my observations in different chromatin classes, where promoter and quiescent states were both the most concordant between individuals and the most shared between cell types.

Within the enhancer-called chromatin, equivalent ordering by inter-individual variability has been described in lymphoblastoid lines, where bivalent enhancers were the most individual-specific, followed by weak enhancers, and finally strongly active enhancers [37], concurring precisely with the ranking I recovered in purified primary breast cell types. Consistency of deployment across individuals and breadth of use across cell types therefore appear to be linked attributes of a shared landscape of regulatory potential, observable both between chromatin classes and within the enhancer compartment itself.

The distinction between stable and variable enhancers may also reflect two complementary biological timescales encoded within the enhancer landscape. Stable enhancers likely represent regulatory programs established during development and maintained by evolutionary constraint to preserve cell identity. In contrast, the variable enhancer compartment appears to capture regulatory programs that remain responsive throughout life to hormonal exposure, immune activation, ageing, and other environmental influences. Such a model provides an explanation for how genetically identical cell types maintain stable lineage identity while remaining capable of adapting to changing physiological conditions.

The genetic analyses further refine this framework. We identified more than 245,000 *cis*-hBTLs, nearly 90% of which were unique to a single breast cell type, demonstrating that genetic effects on chromatin state are highly cell type-specific. However, these variants did not preferentially regulate lineage-defining biological pathways. Instead, *cis*-hBTLs localized within the enhancer landscape of the cell type in which they were identified, at every level of within-cohort concordance, while influencing biological processes broadly shared across breast cell types. This apparent paradox suggests that inherited genetic variation rarely rewires the regulatory circuitry responsible for establishing cell identity. Rather, genetic variants fine-tune regulatory activity within an existing lineage-specific enhancer architecture, modulating the magnitude or responsiveness of conserved biological programs without fundamentally altering cell fate.

This interpretation is further supported by the distinct behavior of active and repressive histone modifications. Active marks exhibited greater densities of *cis*-hBTLs, significant allelic imbalance, and preferential disruption of lineage-relevant transcription factor binding motifs, whereas repressive chromatin showed comparatively weak genetic associations. Previous histone QTL studies have largely focused on active chromatin, and our findings suggest that the mechanisms linking genotype to chromatin state differ fundamentally between active regulatory elements and repressive chromatin domains. Moreover, the enrichment of environmentally responsive transcription factor motifs within the variable enhancer compartment suggests that many genotype-dependent chromatin effects may only become phenotypically manifest under particular environmental or inflammatory contexts.

The functional validation of rs75071948 at the *ANXA1* locus provides direct experimental support for this model. Introduction of the alternate allele was sufficient to establish de novo H3K4me3 deposition and increase *ANXA1* transcription, confirming that naturally occurring genetic variation can directly modify both chromatin state and gene expression. The observation that, in primary tissue, this effect was detected in LCs, while the edited allele was also sufficient to have an impact in MCF10A cells, further emphasizes that the functional consequence of a regulatory variant depends on the regulatory architecture of the cell in which it is assessed. Given the diverse and sometimes contradictory associations between *ANXA1* expression and breast cancer subtypes, these findings further illustrate how the same regulatory variant may have distinct biological consequences depending on cellular context.

More broadly, our findings carry important implications for interpreting non-coding genetic variation. Increasing evidence demonstrates that disease-associated variants frequently reside within regulatory DNA, yet predicting their functional consequences remains challenging. Our results suggest that regulatory variants should not be considered independently of the epigenomic landscape in which they occur. Instead, cell type-specific enhancer landscapes provide the regulatory scaffold upon which both inherited genetic variation and environmentally responsive regulatory programs act. Integrating genotype with cell type-resolved epigenomic maps therefore offers a powerful framework for interpreting non-coding variation in normal biology, disease susceptibility, and cancer initiation.

## Conclusions

We define a two-compartment organization of the human breast enhancer epigenome in which stable enhancers encode the lineage-defining regulatory architecture of each cell type, whereas a variable enhancer compartment captures environmentally responsive and genetically influenced regulatory activity. Genetic variation acts predominantly within this regulatory framework to modulate chromatin state and gene expression in a highly cell type-specific manner, as demonstrated by functional validation of rs75071948 at the *ANXA1* locus. Together, these findings provide a conceptual framework for understanding how cell identity, genotype, and environmental exposure interact to shape the human epigenome and influence disease susceptibility.

## Methods

### Breast tissue dissociation and cell sorting

Breast tissue was obtained as previously described [50,129]. Briefly, histologically normal breast tissue was obtained with informed consent from eight healthy premenopausal adult female donors undergoing reduction mammoplasty surgeries and used according to procedures approved by the Research Ethics Board of the University of British Columbia. Fresh tissue was viably cryopreserved as enzymatically isolated organoids and then thawed and dissociated into single-cell suspensions and the four defined breast cell types isolated by fluorescence-activated cell sorting (FACS). After sorting, cells were spun to remove sheath fluids and acquire a cell pellet.

Cells were then resuspended in ChIP-seq lysis buffer (0.1% Triton, 0.1% Deoxycholate, 1X PIC) prior to snap freezing before ChIP-seq or DNA extraction. For RNA-seq extraction, after aspirating supernatant from cell sorting, cells were directly snap frozen without resuspension.

### Whole genome library preparation and sequencing

All extraction, library preparation, and sequencing were performed following the guidelines formulated by IHEC (http://www.ihec-epigenomes.org). One cell type was selected per individual for WGS (LP for individual 1, BC for individuals 2 through 5, and LC for individuals 6 through 8). Standard procedures for ChIP-seq, and RNA-seq, are available at https://thisisepigenomics.ca/for-scientists/protocols-and-standards. For whole genome sequencing, library bias and coverage gaps associated with PCR amplification of high GC or AT-rich regions were minimized through implementation of a version of the TruSeq DNA PCR-free kit (New England Biolabs, cat. no. E6875-6877B-GSC), automated on a Microlab NIMBUS liquid handling robot (Hamilton Robotics). Briefly, 500 ng of genomic DNA was arrayed in a 96-well microtiter plate and subjected to shearing by sonication (Covaris, cat. no. LE220).

Sheared DNA was end-repaired and size selected using paramagnetic PCRClean DX beads (ALINE Biosciences, cat. no. C-1003-450) targeting a 300 to 400 bp fraction. After 3’ A-tailing, full length TruSeq (Individuals 1 to 5) or GSC TSV3 (Full) UMI (Individuals 6 to 8) adapters were ligated. Libraries were purified using paramagnetic beads (ALINE Biosciences, cat. no. C-1003-450). PCR-free genome library concentrations were quantified using a qPCR Library Quantification kit (KAPA, cat. no. KK4824) prior to sequencing with paired-end 150 base reads on the Illumina platform.

### Variant calling

FASTQC (Andrews *et al.*, unpublished) and fastp [130] were used to verify library quality. BWA MEM [47] v0.7.6a -M was used to align sequences from each sequencing lane to an hg38 genome with ALT contigs removed, and sequencing lane alignments were sorted and merged for each sample using SAMtools [131] v1.10. WGS data from all samples were variant called on GATK v4.1.4.1 using recommended best practices [132]. Duplicate reads were marked using GATK MarkDuplicatesSpark, and base quality score recalibration was performed with GATK BaseRecalibrator and ApplyBQSR using dbSNP SNPs [49], 1000 Genomes Project Phase 3 indels [133], and Mills indels [134] as gold standards. GATK HaplotypeCaller was used to call variants, and SNVs were selected and filtered per GATK recommendations for hard-filtering germline short variants. Variants were then phased using WhatsHap [135].

### ChIP sequencing

Native chromatin immunoprecipitation (N-ChIP) was prepared from FACS sorted cells with > 90% purity for the target cell type (∼100,000 cells per immunoprecipitation). Cells were lysed with lysis buffer (0.1% Triton X-100, 0.1% Deoxycholate, 10mM sodium butyrate) plus protease inhibitor for 20 minutes on ice. The extracted chromatin was then digested with 90U of MNase enzyme (New England Biolabs, cat. no. M0247S) for 6 minutes at 25°C, the reaction was quenched by adding 5.5 µL of 250 µM EDTA. A mix of 1% Triton X-100 and 1% Deoxycholate was then added to the digested samples and the 96-well plate of samples was chilled on ice for 20 minutes. The digested chromatin was then pooled and 12 µL of chromatin was reserved as input control. The rest of the digested chromatin was pre-cleared in IP buffer (20 mM Tris-HCl [pH 7.5], 2 mM EDTA, 150 mM NaCl, 0.1% Triton X-100, 0.1% Deoxycholate) plus protease inhibitor with 20 µL of pre-washed Protein A/G Dynabeads (Invitrogen, cat. no. 100-02D) at 4°C for 1.5 hours. Supernatants were removed from the beads and transferred to a 96-well plate containing the antibody-bead complex. The plate was sealed and incubated at 4°C on a rotating platform overnight. The reaction plate containing the immunoprecipitation samples was placed on a magnetic plate and samples were washed twice with low salt buffer (20 mM Tris-HCl [pH 8.0], 0.1% SDS, 1% Triton X-100, 2 mM EDTA, 150 mM NaCl) and twice with high salt buffer (20 mM Tris-HCl [pH 8.0], 0.1% SDS, 1% Triton X-100, 2 mM EDTA, 500 mM NaCl). DNA-antibody complexes were eluted in 30 µL Elution Buffer (100 mM NaHCO3, 1% SDS), incubated at 65°C for 1.5 hours with mixing speed at 1350 rpm on a thermomixer. Protein digestion using 1.75 µL of Qiagen Protease was performed on the eluted DNA samples at 50°C for 30 minutes with mixing at 600 rpm on a thermomixer. ChIP DNA was then purified using Sera-Mag beads (Fisher Scientific, cat. no. 09-981-123) with 30% PEG before library construction.

Library construction was prepared by following a modified paired-end library protocol (Illumina). Briefly, the DNA was subject to end-repair and phosphorylation by T4 DNA polymerase with Klenow DNA Polymerase and T4 polynucleotide kinase, respectively, in a single reaction. 3’ A overhangs were generated using Klenow fragment (3’ to 5’ exo minus) and ligated to Illumina PE adapters (Individuals 1 to 5) or TruSeqv3 short adapters with UMI (Individuals 6 to 8), that contain 5’ T overhangs. The adapter-ligated products were purified using PCR Clean DX beads (ALINE Biosciences, cat. no. C-1003-450), then PCR-amplified with NEBNext Ultra II Q5 master Mix (New England Biolabs, cat. no. M0544) in 8 cycles (for H3K4me1, H3K4me3, H3K9me3, H3K27me3, H3K36me3) and 10 cycles (for H3K27ac) using Illumina’s PE primer set (Illumina). PCR product was purified using PCR Clean DX beads (ALINE Biosciences, cat. no. C-1003-450), and the DNA quality was assessed and quantified using the Caliper LabChip GX DNA High Sensitivity assay (PerkinElmer, cat. no. 760568) and the Quant-iT dsDNA high sensitivity assay (Thermo Fisher Scientific, cat. no. Q33120).

Libraries were normalized and pooled and the final concentration of the pooled library was determined by Qubit dsDNA HS Assay Kit using a Qubit fluorometer (Thermo Fisher Scientific). Clusters were generated on the Illumina cluster station, and sequencing was run on the Illumina platform following the manufacturer’s instructions.

### ChIP-seq analysis

Library quality was verified as for variant calling, and BWA MEM [47] v0.7.6a -M was used to align sequences (75 bp paired-end reads) from each sequencing lane to hg38, with reads below mapping quality 10 filtered out. WASP [51] was used to further filter out reads whose alignment position shifted when aligned over individual-specific haplotypes versus reference haplotypes from hg38. Resulting alignments were peak called using MACS2 [53] v2.2.7.1 callpeak with a *q*-value cutoff of 0.01 for H3K4me3, H3K27ac, and H3K4me1, and --broad mode with a *q*-value cutoff of 0.05 for H3K9me3, H3K27me3, and H3K36me3.

### RNA sequencing

Qualities of total RNA samples were determined using an Agilent Bioanalyzer RNA Nanochip or Caliper RNA assay and arrayed into a 96-well plate (Thermo Fisher Scientific, cat. no. 14222327). Polyadenylated (PolyA+) RNA was purified using the NEBNext Poly(A) mRNA Magnetic Isolation Module (New England Biolabs, cat. no. E7490L) from 500 ng total RNA. Messenger RNA selection was performed using NEBNext Oligo d(T)_25_ beads (New England Biolabs, cat. no. S1419S) incubated at 65°C for 5 minutes followed by snap-chilling at 4°C to denature RNA and facilitate binding of poly(A) mRNA to the beads. mRNA was eluted from the beads in Tris Buffer incubated at 80°C for 2 minutes then held at 25°C for 2 minutes. RNA binding buffer was added to allow the mRNA to re-bind to the beads, mixed 10 times and incubated at room temperature for 5 minutes. The sample plate was placed on the magnet and the supernatant discarded. The mRNA bound beads were washed twice then cleared again on magnet. The supernatant was again discarded, and mRNA eluted from the beads in 20 µL Tris buffer incubated at 80°C for 2 minutes. mRNA was transferred to a new 96-well plate.

First-strand cDNA was synthesized from heat-denatured purified mRNA using a Maxima H Minus First Strand cDNA Synthesis kit (Thermo Fisher Scientific, cat. no. K1651) and random hexamer primers at a concentration of 200 ng/µL along with a final concentration of 40 ng/µL Actinomycin D, followed by PCR Clean DX (ALINE Biosciences, cat. no. C-1003-450) bead purification on a Microlab NIMBUS robot (Hamilton Robotics). The second strand cDNA was synthesized following the NEBNext Ultra Directional Second Strand cDNA Synthesis protocol (New England Biolabs, cat. no. E7550S) that incorporates dUTP in the dNTP mix, allowing the second strand to be digested using USER enzyme (New England Biolabs, cat. no. M5505) in the post-adapter ligation reaction and thus achieving strand specificity.

cDNA was fragmented by Covaris LE220 sonication to achieve 250-300 bp average fragment lengths. The paired-end sequencing library was prepared following the BC Cancer Genome Sciences Centre strand-specific, plate-based library construction protocol on a Microlab NIMBUS robot (Hamilton Robotics). Briefly, the sheared cDNA was subject to end-repair and phosphorylation in a single reaction using an enzyme premix (New England Biolabs) containing T4 DNA polymerase, Klenow DNA Polymerase and T4 polynucleotide kinase, incubated at 20°C for 30 minutes. Repaired cDNA was purified in 96-well format using PCR Clean DX beads (ALINE Biosciences, cat. no. C-1003-450), and 3’ A-tailed (adenylation) using Klenow fragment (3’ to 5’ exo minus) and incubation at 37°C for 30 minutes prior to enzyme heat inactivation. Illumina TruSeq adapters were ligated at 20°C for 15 minutes. The adapter-ligated products were purified using PCR Clean DX beads, then digested with USER enzyme (1 U/µL, New England Biolabs, cat. no. M5505) at 37°C for 15 minutes followed immediately by 10 cycles of indexed PCR using NEBNext Ultra II Q5 Master Mix (New England Biolabs, cat. no. M0544) and Illumina’s primer set. PCR parameters: 98°C for 1 minute followed by 10 cycles of 98°C for 15 seconds, 65°C for 30 seconds and 72°C for 30 seconds, and then 72°C for 5 minutes. The PCR products were purified and size-selected using a 1:1 PCR Clean DX beads-to-sample ratio (twice), and the eluted DNA quality was assessed with Caliper LabChip GX for DNA samples using the High Sensitivity Assay (PerkinElmer, cat. no. 760568) and quantified using a Quant-iT dsDNA High Sensitivity Assay Kit (Thermo Fisher Scientific, cat. no. Q33120) on a Qubit fluorometer (Invitrogen) prior to library pooling and size-corrected final molar concentration calculation for Illumina paired-end sequencing.

### RNA-seq analysis

RNA-seq analysis was performed as previously described [50,55] using Ensembl v94 [136]. Briefly, adaptor sequences were stripped, and reads were aligned to a transcriptome reference consisting of genomic sequence (GRCh37-lite, July 2010) supplemented by read-length-specific exon–exon junction sequences. A custom RNA quality control and analysis pipeline was used to generate a report and calculate a normalization constant for computing RPKM values (reads per kilobase per million mapped reads). Resulting RPKMs for Ensembl genes were used in subsequent analysis steps.

### Analysis of cell type-specific histone modification cross-cohort variation

Using simulated data (Additional file 1: **Fig. S4**) as an illustrative example, latent modification statuses were randomly sampled using the group modification probability 0_t_ (gray). Latent modifications are either observed (true positives) or unobserved (false negatives) at sample-specific rates using the false-negative (ε_i_^fn^) parameters. False positives are randomly generated using the sample-specific false-positive (ε_i_^fp^) parameters. The observed data (black) is the sum of latent peaks (purple) and false positives (red). After computation of histone mark bin 0 estimates, ENCODE blacklist regions [137] were removed from consideration, and remaining genome-wide bins with nonzero values were classified into quartiles. The 50 bp autosomal bins assigned a posterior mean probability of peak presence 0 were then discretized as blacklisted (0 undefined), unoccupied (0 = 0), highly variable (0 < 0 ≤ 0.25), moderately variable (0.25 < 0 ≤ 0.50), slightly variable (0.50 < 0 ≤ 0.75) and stable (0 > 0.75). Genes whose promoters were uniquely covered with low-varying regions were used as inputs for GO term enrichment analysis using gprofiler2 [138]. Promoters were defined as 2,000 bp upstream and 200 bp downstream of transcription start sites, and enhancer definitions were based on a set previously defined for breast cell types [50].

### Chromatin state segmentation and inter-individual concordance analysis

Chromatin state segmentation was performed using ChromHMM [139]. Peak coordinates were converted to binary presence/absence calls in non-overlapping 200 bp genomic bins tiling the hg38 canonical autosomes and sex chromosomes. Each bin was scored as 1 (overlapping at least one peak) or 0 (no overlap). For the single instance where a histone mark assay was unavailable (H3K9me3 in one individual’s BC sample), bins were encoded as 2, ChromHMM’s designated missing-data value, which causes the model to marginalize over the missing mark during Viterbi decoding.

Chromatin states were assigned using the pretrained Roadmap Epigenomics 18-state “core + K27ac” expanded model, originally trained on 98 reference epigenomes profiled for the same six histone modifications [9,139]. The MakeSegmentation command was used to apply the Viterbi algorithm with fixed emission and transition parameters, producing a per-cell BED-format segmentation assigning each contiguous interval to one of 18 chromatin states with established mnemonic labels: TssA, TssFlnk, TssFlnkU, TssFlnkD, Tx, TxWk, EnhG1, EnhG2, EnhA1, EnhA2, EnhWk, ZNF/Rpts, Het, TssBiv, EnhBiv, ReprPC, ReprPCWk, and Quies. The pretrained model was parsed via the segmenter Bioconductor package.

To quantify inter-individual concordance, a base-pair-weighted Jaccard index was computed for each state between every pair of individuals: *J*(*A*_s_, *B*_s_) = Σ width(*A*_s_ ∩ *B*_s_) / Σ width(*A*_s_ ∪ *B*_s_), where intersection and union are base-pair-level set operations computed using the Bioconductor plyranges package’s reduce_ranges(), intersect_ranges(), and union_ranges() functions. Two comparison frameworks were defined: within-cell type (28 pairs per cell type, 112 total) and cell type-versus-other-three. One-sided Wilcoxon rank sum tests (alternative: within > between) were used to assess whether within-cell type concordance exceeded between-cell type concordance for each state, with BH correction across 18 tests (Additional file 1: **Fig. S13**).

### Enhancer segment intersection with ENCODE cCREs

To assess the relationship between inter-individual enhancer concordance and independently cataloged regulatory elements, ChromHMM segments were intersected with enhancer-class candidate *cis*-regulatory elements (cCREs) from the ENCODE Registry v4 [120]. The registry was filtered to distal enhancer and proximal enhancer cCRE classes. For each cell type, genome-wide segments produced and annotated with the proportion of individuals in which any enhancer ChromHMM state (EnhA1, EnhA2, EnhG1, EnhG2, EnhWk, or EnhBiv) was called and classified into variability categories: stable (> 75% of individuals), low (50–75%), moderate (25–50%), high (≤ 25%), and not present (0%). Enhancer cCRE coverage was computed per segment as the union base pair overlap with enhancer cCREs divided by segment width, using join_overlap_intersect() followed by GenomicRanges::reduce() to avoid double-counting overlapping cCREs. Among segments overlapping at least one enhancer cCRE, the weighted mean H3K27ac max Z score of the intersecting cCREs was computed, weighting by intersection width. JT trend tests (DescTools::JonckheereTerpstraTest, alternative = “increasing”) were used to assess monotonic trends in cCRE strength across variability categories. This analysis was also extended to CD4^+^ and CD8^+^ naive T cells from the IHEC EpiAtlas to assess generalizability.

Eight samples of each T cell subtype were selected from the IHEC metadata, and their pretrained 18-state ChromHMM segmentation BED files were downloaded from the IHEC post-processed data repository (Additional file 11: **Table S10**).

### Cross-cell type sharing of enhancer regions

To test for sharing of enhancer regions across cell types, the full enhancer territory of each cell type was first defined as all segments assigned to any variability category and merged into a set of non-overlapping ranges with GenomicRanges::reduce(). For each focal cell type and category in turn, that cell type’s category territory was pooled with the full enhancer territory of the other three, then partitioned using GenomicRanges::disjoin(), cutting pooled ranges at every breakpoint contributed by any of the four sets, and resulting atomic intervals were restricted to those falling inside the focal territory with IRanges::subsetByOverlaps(). Enhancer interval widths were obtained with width() and summed within each combination of focal cell type, variability category and sharing count, then expressed in megabases. Pooling across focal cell types was performed on these base-pair totals, and proportions were then computed within each variability category, giving the fraction of that category’s coverage shared with zero, one, two or three other cell types. Sharing was further computed on a category-matched basis, in which the four cell types’ regions in a given variability category were pooled, disjoined, and each interval labelled with the number of cell types assigning it to that same category. Because intervals shared by any number of cell types contribute to that grouping’s territory, proportions were reweighted by the number of cell types in that grouping to place them on the same per-cell type denominator as the presence measure.

Specific enhancer state compositions were also computed per cell type and variability category as the share of enhancer state calls assigned to each of the six enhancer states, weighting each segment by its width and by the number of individuals making each call, so that the result reports the composition of state calls rather than of segments.

To test whether the sharing gradient reflected differences in marking strength rather than breadth of use, segments overlapping an enhancer cCRE were assigned to deciles of ENCODE Registry H3K27ac maximum Z-scores computed across all cell types and categories jointly, and the proportion of territory shared with all three other cell types was recomputed within each decile. Sharing was computed on the disjoined partition as above, with each atomic interval inheriting the Z-score of the segment containing it.

### Pathway enrichment analysis of stable and variable enhancer segments

Pathway enrichment was performed using two complementary approaches. GREAT [140] was used for region-based enrichment, associating genomic intervals with nearby genes using the basal-plus-extension gene regulatory domain model with hg38 TSS annotations and testing for enrichment in Reactome pathways (MSigDB C2:CP:REACTOME). Enrichments were computed separately for each combination of cell type (LC, LP, BC, SC, CD4^+^ T cell, CD8^+^ T cell), enhancer state (merged enhancer and each individual enhancer state), and variability category (high, moderate, low, stable).

To quantify the degree to which enriched terms reflected expected cell type- and tissue-specific biology, a relevance scoring framework was developed. A curated pattern legend of 10 categories was defined, spanning mammary-specific terms (mammary/breast/ductal, lactation/milk, ductal branching morphogenesis), cell type-lineage terms (luminal, basal/myoepithelial, stromal/fibroblast), and broader developmental signaling terms (estrogen/progesterone, WNT/Notch/TGFβ/EGFR, epithelial development, ECM/basement membrane) (Additional file 2: **Table S1**). Each significant enrichment term (*p* < 0.05) was matched against the pattern legend using case-insensitive regular expressions. Hits were tallied as a percentage of all significant terms, and this percentage of significant terms matching the relevance pattern was computed per variability category and used to compare the functional specificity across enhancer stability (**Fig. 3c**, Additional file 1: **Fig. S19**).

To test for directional changes in mammary relevance across the variability gradient, JT tests for ordered alternatives were computed for each merged enhancer category, with observations pooled across all four breast cell types (n = 16 per enhancer type: 4 cell types × 4 variability bins ordered from high variability through moderate, low, and stable). The JT test evaluates whether the distributions of mammary relevance fractions across the four ordered variability groups are stochastically ordered, with the test statistic counting concordant pairs between groups of increasing stability rank. *P*-values were computed using the normal approximation with tie correction (DescTools::JonckheereTerpstraTest), which is appropriate for grouped data with n = 4 per group and accounts for tied fraction values across cell types within the same bin. *p*-values were corrected for multiple testing across the five enhancer categories using the BH method. Spearman’s p between the ordered bin rank and the mammary relevance fraction is reported alongside JT statistics as an interpretable effect-size measure of the monotonic trend.

### Binned transcription factor binding site motif enrichment analysis

To identify sequence-level features distinguishing stable from variable enhancer chromatin compartments, binned transcription factor binding site motif enrichment was performed using monaLisa [94]. For each cell type, ChromHMM enhancer segments (merged across EnhA1, EnhA2, EnhG1, EnhG2, EnhWk, and EnhBiv) were annotated with the enhancer_prop score, defined as the proportion of individuals in which any enhancer state was called at that locus. We tested the stable bin (enhancer_prop > 0.75) and the high-variability bin (enhancer_prop < 0.25) independently against a GC-matched random genomic background.

Segments were resized to a fixed 500 bp width centered on the segment midpoint using GenomicRanges::resize() with fix = “center”, filtered to the canonical autosomes and X chromosome, deduplicated, and sorted. Sequences were extracted from hg38 using BSgenome.Hsapiens.UCSC.hg38 via Biostrings::getSeq() and filtered for greater than 30% ambiguous base content. Motif enrichment was computed using calcBinnedMotifEnrR() against the full JASPAR2024 CORE vertebrate position weight matrix collection (879 motifs) [141] with GC content correction enabled. Motifs were classified as significantly enriched if they reached BH-adjusted *p* < 0.05 and log_2_ enrichment > 0.4 in at least one cell type and compartment. To reduce redundancy from highly similar PWMs, the full motif set was clustered using universalmotif::compare_motifs() with Pearson correlation coefficient ≥ 0.7 and average linkage, yielding 113 archetype clusters across 828 clustered motifs. The clustering approach is inspired by Vierstra *et al.* 2020 [142] but uses our own per-set clustering rather than published archetype assignments.

### Pathway-level enrichment of stable and variable enhancer compartments

Pathway-level enrichment of compartment-specific genomic regions was performed using rGREAT [143] against MSigDB C2:CP:REACTOME gene sets retrieved via msigdbr [144], using the default basal-plus-extension *cis*-regulatory association rules and hg38 RefSeq TSS annotations. Enrichments were computed independently for each cell type (LC, LP, BC, SC) and compartment (stable, variable), yielding 8 panel-level enrichment tables. Binomial *p*-values were Benjamini–Hochberg-corrected within each panel and the full enrichment table (one row per cell type × compartment × Reactome term, with BH-adjusted *p*, log_2_ enrichment, and observed region hit count) is provided as Additional file 6: **Table S5**.

### *Cis*-hBTL mapping, association with gene transcription changes and manual selection

We built an in-house R pipeline using bedr [145], GenomicRanges [146], and plyranges [147] to define and annotate loci where alleles overlapped consistently with peak presence/absence across the cohort of eight individuals for each cell type and histone mark combination. Genomic loci where, across all eight individuals, individuals that carried a particular variant allele, whether reference or alternate, also had an overlapping histone mark peak, and, at the same locus, individuals that did not have that variant allele also had no overlapping histone mark peak (in other words, where variant alleles co-segregated with histone mark peaks) were counted as *cis*-hBTLs. *Cis*-hBTLs were linked to their nearest gene and corresponding RPKM value from each individual/cell type combination. At each *cis*-hBTL, genotype was used as the independent variable and RPKM as the dependent variable, and a Welch’s *t-*test was performed, with *p-*values adjusted for false discovery rate using the BH method [148]. Positions with the largest shift in transcription between genotypes were prioritized by computing RPKM variance at each *cis*-hBTL, and ChIP-seq peaks and treatment pileups surrounding *cis*-hBTL variants were manually verified on IGV to prioritize positions where individuals with peak absence had no nearby peaks.

### *Cis*-hBTL and peak set enrichment analyses

To assess enrichment with sequence features, UCSC knownGene [149] sequence features (transcripts, promoters, introns, exons, coding sequences, five prime UTRs, three prime UTRs, and intergenic regions) were intersected with *cis*-hBTLs and corresponding histone modification peak sets (H3K4me3, H3K27ac, H3K4me1, H3K9me3, H3K27me3, H3K36me3), and the fraction of each *cis*-hBTL set covered by a sequence feature was divided by the fraction of each corresponding peak set covered by a sequence feature, then log_2_ transformed.

To assess the enrichment of *cis*-hBTLs with known eQTL, GTEx v8 eQTL variant sets were downloaded from https://www.gtexportal.org. Variant calls from the breast cohort were annotated with *cis*-hBTL status, either present or not present, as well as eQTL status for all GTEx tissues. For each GTEx tissue, a Fisher’s exact test for *cis*-hBTL and eQTL status was performed, and the odds ratio for representation of eQTL within the *cis*-hBTL set was calculated.

To compute the relative enrichment of histone modification peak sets for a given histone mark across the eight individuals with *cis*-hBTLs, peak sets were tiled into unique sections of the genome with the potential to form a *cis*-hBTL, i.e. having any permutation of 1 to 7 individuals carrying a peak at a given locus. The fraction of the genome covered by tiles intersecting corresponding *cis*-hBTLs carried by the same permutation of individuals was computed for all histone mark and cell type combinations and normalized to the total genomic coverage for the corresponding mark.

### Allele-specific *cis*-regulation analysis

To quantify allelic imbalance in ChIP-seq read counts at heterozygous *cis*-hBTL positions, paired reference- and alternate-allele depths were extracted from the variant-aware (WASP-derived) ChIP-seq alignments using the VariantAnnotation Bioconductor package (readVcfAsVRanges). For each of the 192 unique (individual × cell type × histone mark) combinations available in the cohort, the corresponding sample’s WGS-derived high-confidence SNV call set was filtered to heterozygous positions, defined as positions with a phased allele frequency of exactly 0.5 in the GATK joint-called VCF. The intersection of this heterozygous SNV set with the cell type- and mark-matched *cis*-hBTL set defined the position pool eligible for allelic imbalance testing for that sample. Positions were further restricted to those at which the ChIP-seq treatment alignments produced at least one read carrying each allele (refDepth > 0 and altDepth > 0), excluding monoallelic pileups where allelic ratio is undefined.

For each retained position, reference and alternate read depths were tallied. Two complementary statistical tests were performed. First, per-position raw paired depths were tested across all positions within each histone mark using a one-sided paired *t*-test with the directional alternative testing whether the reference-allele depth was lower than the alternate-allele depth at *cis*-hBTL-driving positions, consistent with the prior expectation that the *cis*-hBTL-driving allele recruits the histone mark. Second, to control for the within-individual correlation among positions and to weight individuals equally regardless of how many heterozygous *cis*-hBTL positions each contributed, per-(individual, cell type) mean depths were computed for the reference and alternate alleles, and a one-sided paired Student’s *t*-test was performed on these aggregated means across the 32 (individual × cell type) combinations contributing to each histone mark. The aggregated form is the test reported in the Results text and used to annotate **Fig. 4d**, while the figure itself displays the per-position log_2_ depth distributions overlaid with significance brackets derived from the aggregated test.

### Enhancer overlap analysis of *cis*-hBTLs across variability categories

To assess whether *cis*-hBTLs preferentially overlapped enhancer regions of their own cell type, and whether this depended on the consistency of enhancer calls across individuals, the variability category assignments described above were used directly. The four categories partition each cell type’s entire enhancer territory. Because this enhancer stability value is computed on the merged enhancer class, a segment in the stable category is one at which > 75% of individuals received some enhancer state, not one at which every individual received the same enhancer state. *Cis*-hBTLs were then filtered to those uncovered in active, enhancer-associated histone marks (H3K27ac, H3K4me1, H3K4me3) and tested for overlap with each cell type’s enhancer sets. For each *cis*-hBTL source cell type by enhancer cell type combination, log_2_ enrichment over genome-wide expectation was computed as observed overlap count divided by expected count under a uniform null. Because *cis*-hBTLs are ascertained within histone mark peaks rather than distributed uniformly, this null inflates fold enrichment for all peak-like territory equally, so values are comparable between enhancer variabilities and cell types, but not calibrated in absolute terms.

### Cell type specificity analysis of *cis*-hBTL functional enrichment

To assess whether *cis*-hBTLs unique to each cell type preferentially enriched for that cell type’s biology at the functional pathway level, we applied GREAT [140] via the local rGREAT implementation [143] to *cis*-hBTL sets stratified by cell type and active histone mark (H3K27ac, H3K4me1, H3K4me3). Variants were reduced to non-overlapping intervals using GenomicRanges::reduce() to avoid double-counting in the GREAT binomial test. Enrichment was computed against two gene set collections: the MSigDB Hallmark (H) and Curated (C2) gene sets retrieved via msigdbr. TSS annotations from the TxDb.Hsapiens.UCSC.hg38.knownGene package were used to define GREAT regulatory domains under the default basal-plus-extension model (5 kb upstream, 1 kb downstream, extended up to 1 Mb). Enrichment tables were extracted with getEnrichmentTable() using min_region_hits = 2.

To classify enriched terms by their biological relevance to each breast cell type, a curated keyword reference table was constructed with regex patterns drawn from canonical marker studies of breast cell type identity [45,46,50,90,128] (Additional file 3: **Table S2**). The keyword set comprised 25 functional categories distributed across the four cell types, covering BC-associated terms (myoepithelial identity, contractile/smooth muscle, ECM/adhesion, basement membrane, polarity, epithelial barrier), LP-associated terms (progenitor/stemness markers, proliferation, ELF5/SOX9, hormone response sensitization, alveolar markers, lineage plasticity), LC-associated terms (luminal identity, hormone response, FOXA1/GATA3 transcription factors, secretory function, tight junction biology, milk protein production), and SC-associated terms (fibroblast identity, adipogenesis, angiogenesis, immune signaling, growth factor secretion, ion transport). A parallel cancer/pathology keyword set covered breast cancer specifically, general cancer hallmarks, cancer signaling pathways, EMT, and breast-specific pathology terms (Additional file 4: **Table S3**). Regex matching was performed case-insensitively against term descriptions with GO accession prefixes stripped.

For each *cis*-hBTL cell type by mark combination, the proportion of significant terms (FDR < 0.05) matching each of the four cell types’ keyword sets and the cancer/pathology keyword set was computed. A self-versus-other specificity metric was defined as log_2_((proportion matching own keyword set) / (mean proportion matching other cell types’ keyword sets)), where positive values would indicate preferential enrichment for own-cell type biology.

### Cell type-specific TF motif disruption by *cis*-hBTLs

To query for instances of TF binding site disruption mediating breast cell type *cis*-hBTLs, SNV-level motif disruption scoring was performed using motifbreakR [150] with JASPAR2024 [141] CORE vertebrate position weight matrices. The cell type-specific TF reference set was constructed from the leading cell type-discriminating TFs identified in Pellacani *et al.* 2016 [50] Fig. 7B, manually resolved to single HGNC gene symbols with matching JASPAR2024 PWMs. The reference set comprised LC-specific TFs (FOXA1, JUN, FOSL2, FOSL1, FOXP1), LP-specific TFs (EHF, ELF5, NFIB, ETS1), and BC-specific TFs (TP63, TP53, NFIB, TEAD4).

For all *cis*-hBTL positions across all six histone marks, reference and alternate allele information was retrieved from dbSNP build 155 via SNPlocs.Hsapiens.dbSNP155.GRCh38, filtering to single-nucleotide variants with unambiguous REF and ALT alleles. Motif disruption was computed with motifbreakR using the information-content (IC) scoring method, which quantifies the change in motif match score caused by the alternate allele relative to the reference allele, weighted by position-specific information content of the PWM. A *p*-value threshold of 1 × 10^-4^ was applied to identify motif matches, and variants producing “strong” or “weak” effect classifications (as defined by motifbreakR’s internal IC-based thresholds) were counted as disrupting the motif. IC scoring was chosen over the default log-odds scoring because it is more sensitive to subtle but biologically meaningful disruptions at high-information positions within the motif.

To test whether *cis*-hBTLs from a given cell type preferentially disrupted TF motifs of their own cell type, the motif disruption results were pivoted to a long-form design in which each *cis*-hBTL variant contributed one observation per focal TF epithelial cell type (LC, LP, or BC), yielding three rows per variant. For each row, a binary disruption outcome was defined as whether the variant disrupted at least one PWM belonging to the focal TF cell type’s reference set, and a binary predictor (*is_match*) was defined as whether the *cis*-hBTL’s source cell type matched the focal TF cell type.

Per-mark inference was performed by fitting a logistic generalized linear model (GLM) of the binary disruption outcome on *is_match* separately for each histone mark, using all three breast cell types with defined TF reference sets (BC, LP, LC). Because each variant contributes three rows to the long-form dataset, observations from the same variant are not independent. To account for this within-variant correlation, cluster-robust standard errors were computed using the sandwich HC0 estimator (sandwich::vcovCL, clustered by variant coordinate identifier), and coefficient inference was performed via the Wald *z*-test (lmtest::coeftest). The exponentiated GLM coefficient for *is_match* is reported as the odds ratio (OR), with 95% confidence intervals derived from the cluster-robust standard error. Two-sided Wald *p*-values were adjusted for multiple testing using the BH method across all six histone marks, and one-sided *p*-values (testing OR > 1) were derived from the Wald *z* statistic.

### Cell culture

MCF10A cells were a gift from the Connie Eaves Lab (Terry Fox Laboratories, BC Cancer Research Centre, Vancouver, BC, Canada). Cells were cultured in DMEM/F-12 (STEMCELL Technologies, cat. no. 36254), insulin (10 ug/ml) (Sigma-Aldrich, cat. no. I6634), hydrocortisone (0.5 ug/ml) (Sigma-Aldrich, cat. no. H4881), cholera toxin (100 ng/ml) (Sigma-Aldrich, cat. no. C8052), EGF (20 ng/ml) (Sigma-Aldrich, cat. no. E9644), and horse serum (5%). When cells were 80% confluent, they were trypsinized for 15 minutes, and cells were pelleted by spinning down at 150G for 10 minutes at 20°C. Pelleting in preparation for RT-qPCR or sequencing was done at 300G for 5 minutes at 4°C. Pellets were then flash frozen in an ethanol + dry ice mixture and stores at -80°C.

### CRISPR/Cas9-mediated HDR of MCF10A

Alt-R S.p. Cas9 Nuclease V3 (Integrated DNA Technologies, cat. no. 1081058) and Alt-R CRISPR-Cas9 sgRNA (Integrated DNA Technologies, custom design specified in Additional file 10: **Table S9**) ribonucleoprotein along with Alt-R HDR Donor Oligo (Integrated DNA Technologies, custom design specified in Additional file 10: **Table S9**) and Alt-R HDR Enhancer V2 (Integrated DNA Technologies, cat. no. 10007910) were electroporated into MCF10A using the Neon electroporation system (Thermo Fisher Scientific) according to manufacturer protocols. The region of interest surrounding the edit site was lifted out through PCR (primer design specified in Additional file 10: **Table S9**). To test the efficiency of electroporation, libraries were indexed, pooled, and sequenced with Illumina P3 reagents for 200 cycles (Illumina, cat. no. 20040560) using 100 bp paired-end sequencing chemistry on the Illumina NextSeq 2000 platform, following manufacturer protocols (Illumina). Reads were aligned to hg38 using BWA MEM [47] and the haplotypes of reads around rs75071948 were tallied using a custom R script. In the electroporated MCF10A population, around 1% of reads contained the rs75071948-G alternate genotype with no off-target edits within the amplicon window, with another 13% of reads carrying the alternate genotype as well as at least one off-target edit within the amplicon window. To acquire clones containing only the desired variant allele, a series of sorting steps were undertaken. Single-cell sorting did not yield any clones with the desired genotype in the 51 lines that were grown and sequenced. However, sorts of 100 cells per well yielded one line containing the desired genotype at a frequency of 31%, so additional single-cell sorts were cloned out of this line, yielding 10 clones with a biallelic knock-in of rs75071948-G and no off-target edits which were pelleted for downstream validation steps.

### Validation RT-qPCR

RNA was extracted from parental and rs75071948-G MCF10A frozen cell pellets using an AllPrep DNA/RNA Mini Kit (QIAGEN, cat. no. 80204) and quantified on an Agilent 2100 Bioanalyzer System using an RNA 6000 Nano Kit (Agilent, cat. no. 5067-1511). RT-qPCR was performed with TaqPath 1-Step RT-qPCR MM, CG (Thermo Fisher Scientific, cat. no. A15300), and TaqMan probes targeting *ANXA1* (Thermo Fisher Scientific, cat. no. 4453320, assay ID Hs00167549_m1), and *ANXA1* 2^-ΔCt^ in parental and rs75071948-G MCF10A lines was calculated against the parental mean.

### Validation ChIP-seq

H3K4me3 ChIP-seq libraries for parental and rs75071948-G MCF10A lines were constructed according to standard operating procedures available at https://thisisepigenetics.ca/for-scientists/protocols-and-standards. Cells were lysed in mild non-ionic detergents (0.1% Triton X-100 and Deoxycholate) and protease inhibitor cocktail (EMD Millipore, cat. no. 539134-1ML) in order to preserve the integrity of histones harboring epitope of interest during cell lysis. In brief, 100,000 cells were digested by micrococcal nuclease at room temperature for 6 minutes and 0.25mM EDTA was used to stop the reaction. Antibody H3K4me3 (Cell Signaling Technology, cat. no. 9751S) was incubated with anti-IgA magnetic Dynabeads (Invitrogen, cat. no. 10002D) for 2 hours. Digested chromatin was incubated with magnetic beads alone for 1.5 hours. Digested chromatin was separated from the beads and incubated with antibody-bead complex overnight in IP buffer (20mM Tris-HCl pH 7.5, 2mM EDTA, 150mM NaCl, 0.1% Triton X-100, 0.1% Deoxycholate). The resulting IPs were washed 2 times by low salt (20mM Tris-HCl pH 8.0, 2 mM EDTA, 150mM NaCl, 1% Triton X-100, 0.1% SDS) and then high salt (20 mM Tris-HCl pH 8.0, 2 mM EDTA, 500 mM NaCl, 1% Triton X-100, 0.1% SDS) wash buffers. IPs were eluted in elution buffer (1% SDS, 100mM Sodium Bicarbonate) for 1.5 hours at 65°C. Remaining histones were digested by QIAGEN Protease (QIAGEN, cat. no. 19155) for 30 minutes at 50°C and DNA fragments were purified using SeraMag magnetic beads in 30% PEG. Illumina sequencing libraries were generated by end repair, 3’ A-addition, and Illumina sequencing adaptor ligation (New England BioLabs, cat. no. E6000B-10). Libraries were then indexed, PCR amplified (8 cycles) and 150 bp paired-end reads were sequenced on 4 lanes of the Illumina Novaseq 6000 platform (Illumina) following manufacturer protocols. Adaptor sequencing trimming was performed with skewer-0.2.2 [151], and sequence reads were split by index and aligned to human reference genome GRCh38 with BWA (v0.7.6a) MEM [47]. Prior to testing for coverage at rs75071948, BAMs were normalized using DeepTools [152] bamCoverage --minMappingQuality 10 --samFlagExclude 3332 --normalizeUsing RPKM --exactScaling --extendReads --binSize 1 --effectiveGenomeSize 2634848028.

### Source code

All scripts used in this thesis can be found in the following repository: https://github.com/hauduc/cis-hbtl.

## Supplementary Information

Additional file 1: Supplementary figures.

Additional file 2: Table S1. Mammary cell type GREAT term pattern legend. Additional file 3: Table S2. Breast cell type keywords.

Additional file 4: Table S3. Cancer keywords.

Additional file 5: Table S4. Genes with variable expression and variably marked promoters. Additional file 6: Table S5. GREAT curated breast terms (**Fig. S20**)

Additional file 7: Table S6. *Cis*-hBTLs.

Additional file 8: Table S7. *Cis*-hBTL percentages.

Additional file 9: Table S8. Top manually verified *cis*-hBTLs and literature annotations. Additional file 10: Table S9. CRISPR/Cas9 designs.

Additional file 11: Data accessions.

## Declarations

### Ethics approval and consent to participate

This work was performed under the BC Cancer Agency Research Ethics Board certificate number H12-01767.

### Consent for publication

Not applicable.

### Availability of data and materials

The data sets used in this study are available in the Epigenome Reference Registry (EpiRR), and associated accessions can be found in Additional file 11: **Table S10** or upon reasonable request.

## Authors’ contributions

A.H. and M.H. designed the study. A.H. performed the analysis, interpreted the results, performed validation cell culture, and CRISPR editing. J.S. developed the Bayesian algorithm used to measure epigenomic variability across cell types. S.D. performed immunofluorescence and imaging on stromal cells. M.B., M.M. and Q.C. oversaw the generation and quality control of sequence data under the guidance of M.H. A.H. and M.H. wrote the manuscript. All authors read and approved of the final manuscript.

## Acknowledgements

We are grateful to Susanna Tan, who helped provide the MCF10A cell lines used in the validation work, and Ryan Ghorayeb, who gave advice on CRISPR/Cas9 experimental design. **Fig. 1a** was created in BioRender by A.H. https://BioRender.com/c27s373. **Fig. S33** was created in BioRender by A.H. https://BioRender.com/j82z532.

