## Additional file 1 for "Cell type-specific epigenomic variation and its association with genotype in the human breast"

### Slide 1
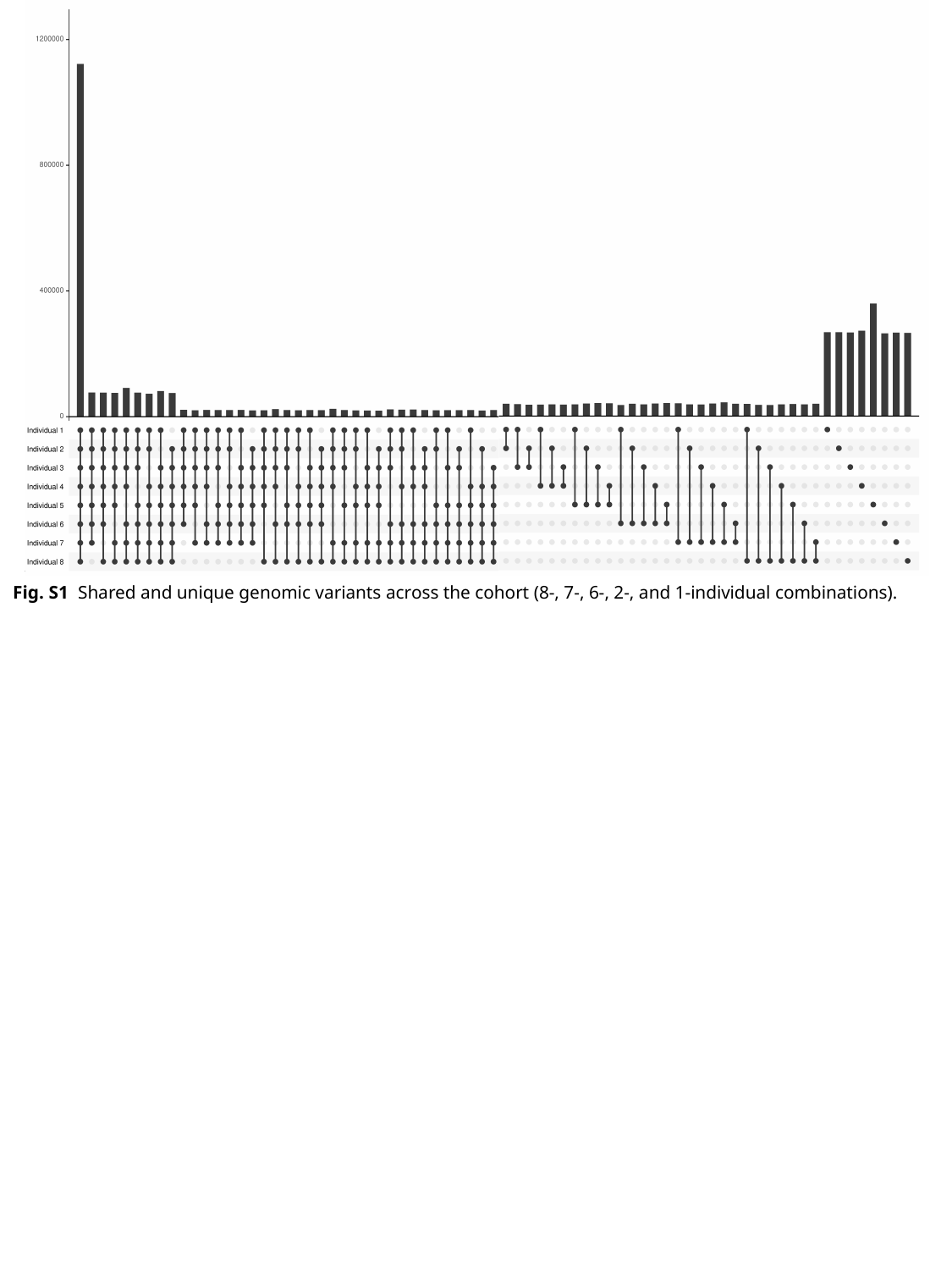

Fig. S1 Shared and unique genomic variants across the cohort (8-, 7-, 6-, 2-, and 1-individual combinations).

### Slide 2
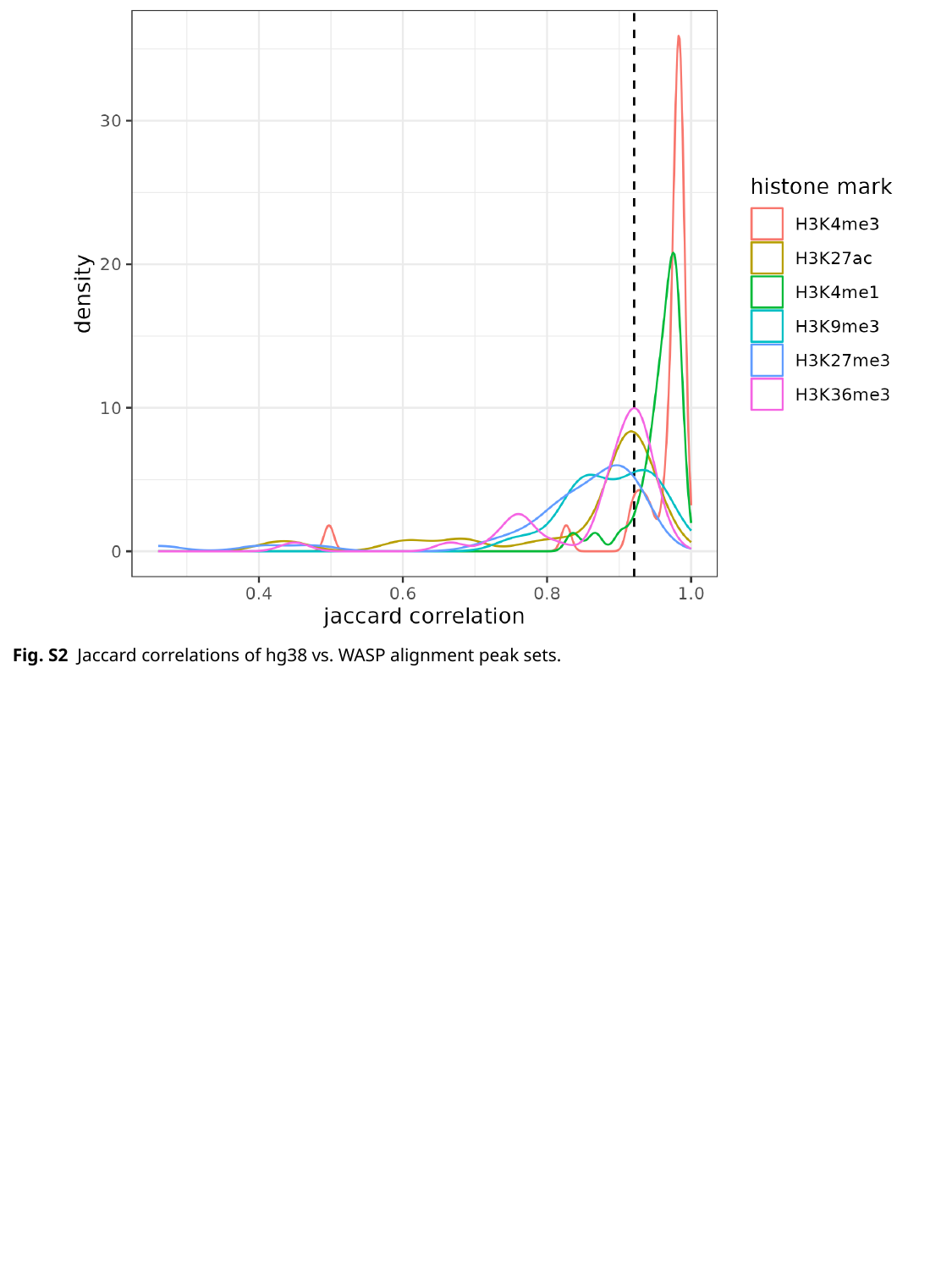

Fig. S2 Jaccard correlations of hg38 vs. WASP alignment peak sets.

### Slide 3
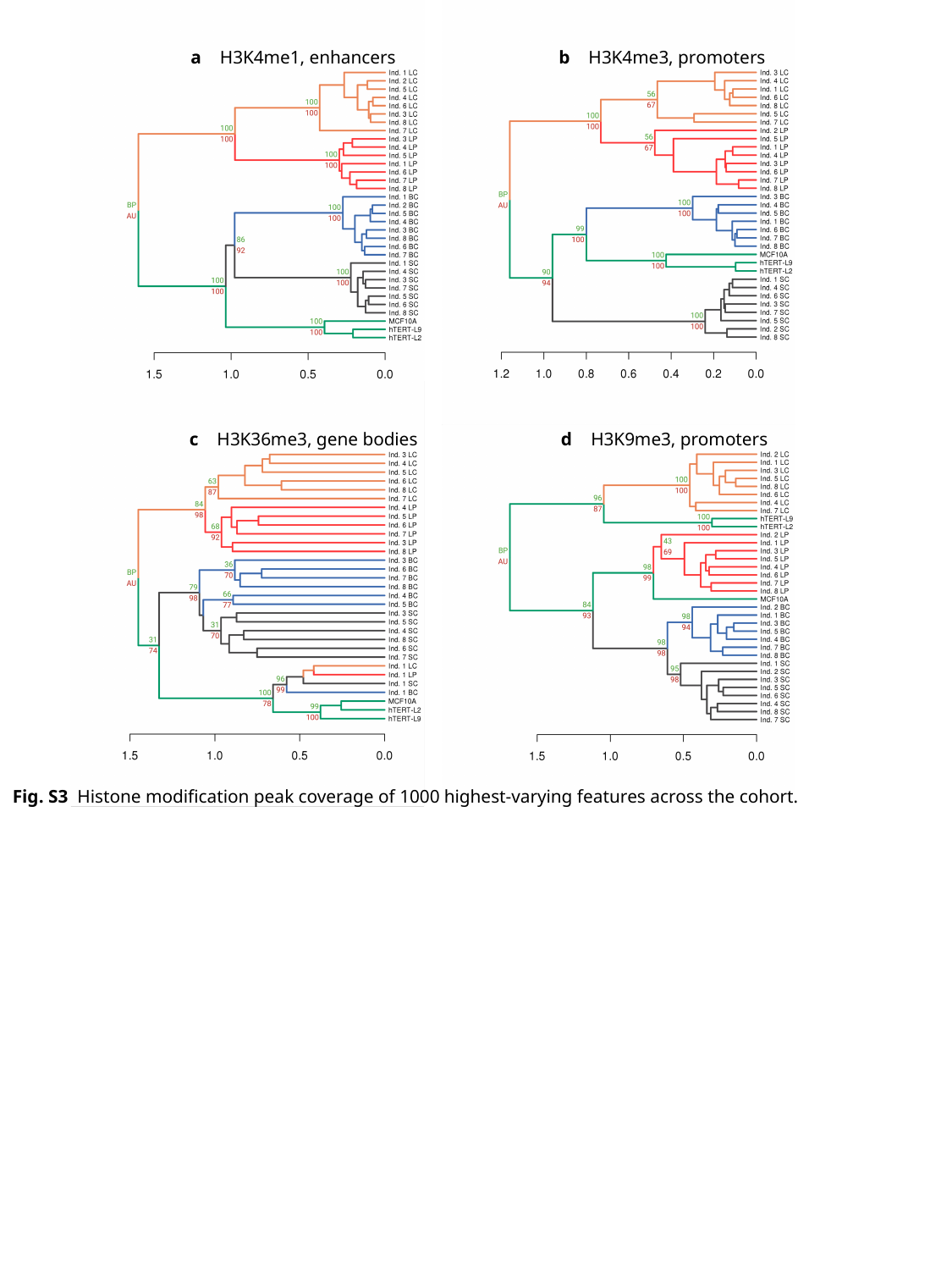

b H3K4me3, promoters
a H3K4me1, enhancers
c H3K36me3, gene bodies
d H3K9me3, promoters
Fig. S3 Histone modification peak coverage of 1000 highest-varying features across the cohort.

### Slide 4
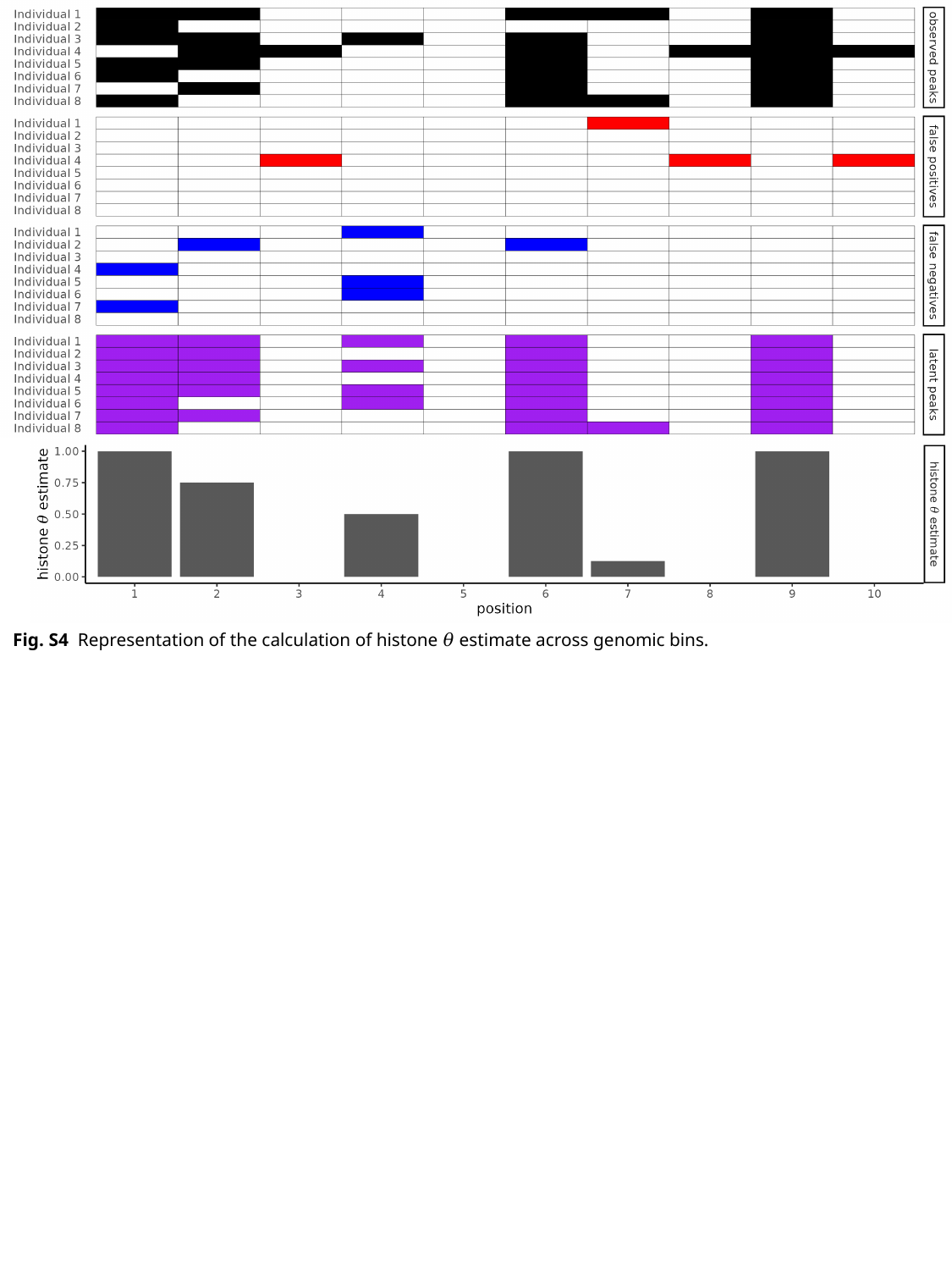

Fig. S4 Representation of the calculation of histone 𝜃 estimate across genomic bins.

### Slide 5
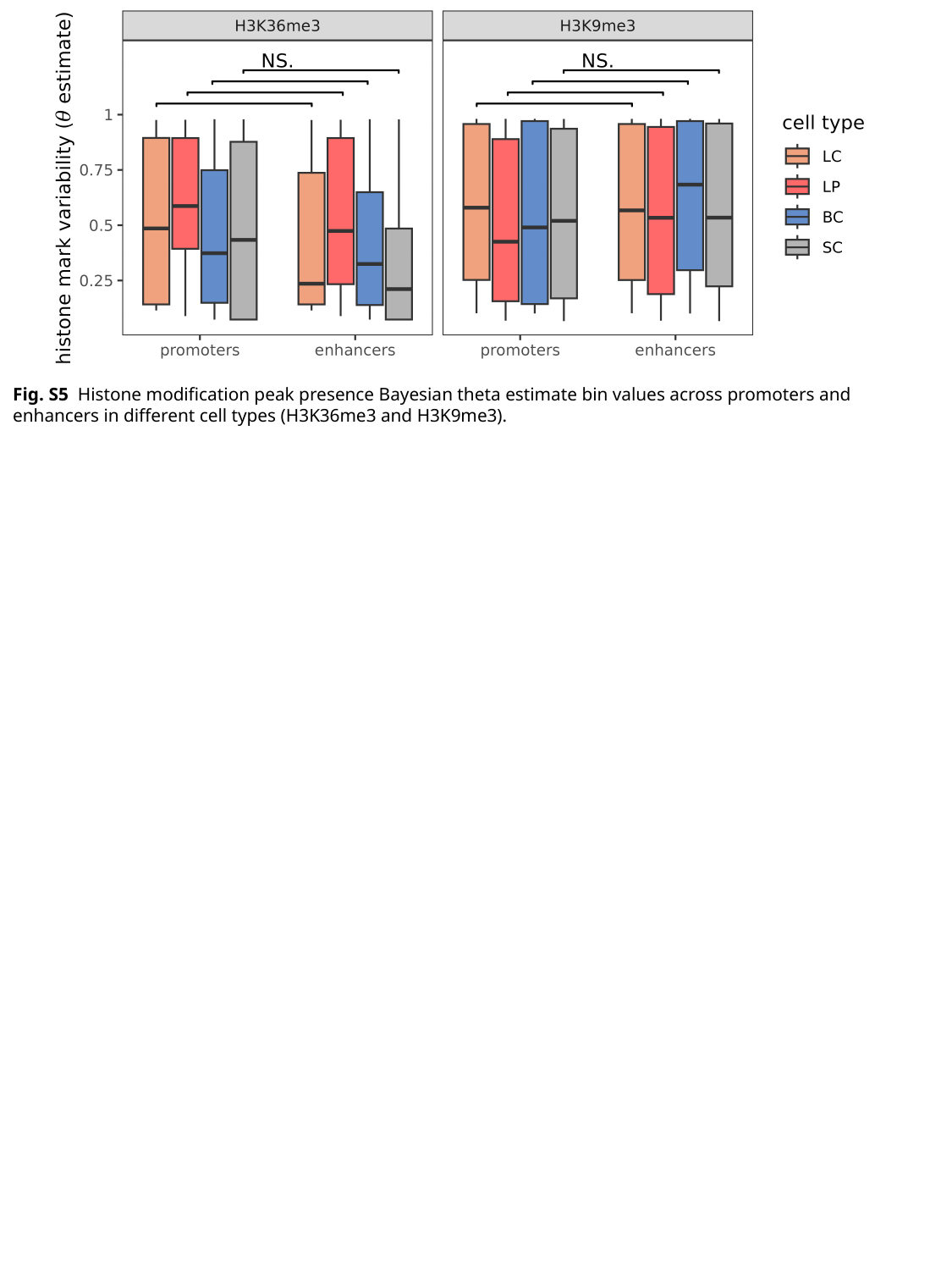

Fig. S5 Histone modification peak presence Bayesian theta estimate bin values across promoters and enhancers in different cell types (H3K36me3 and H3K9me3).

### Slide 6
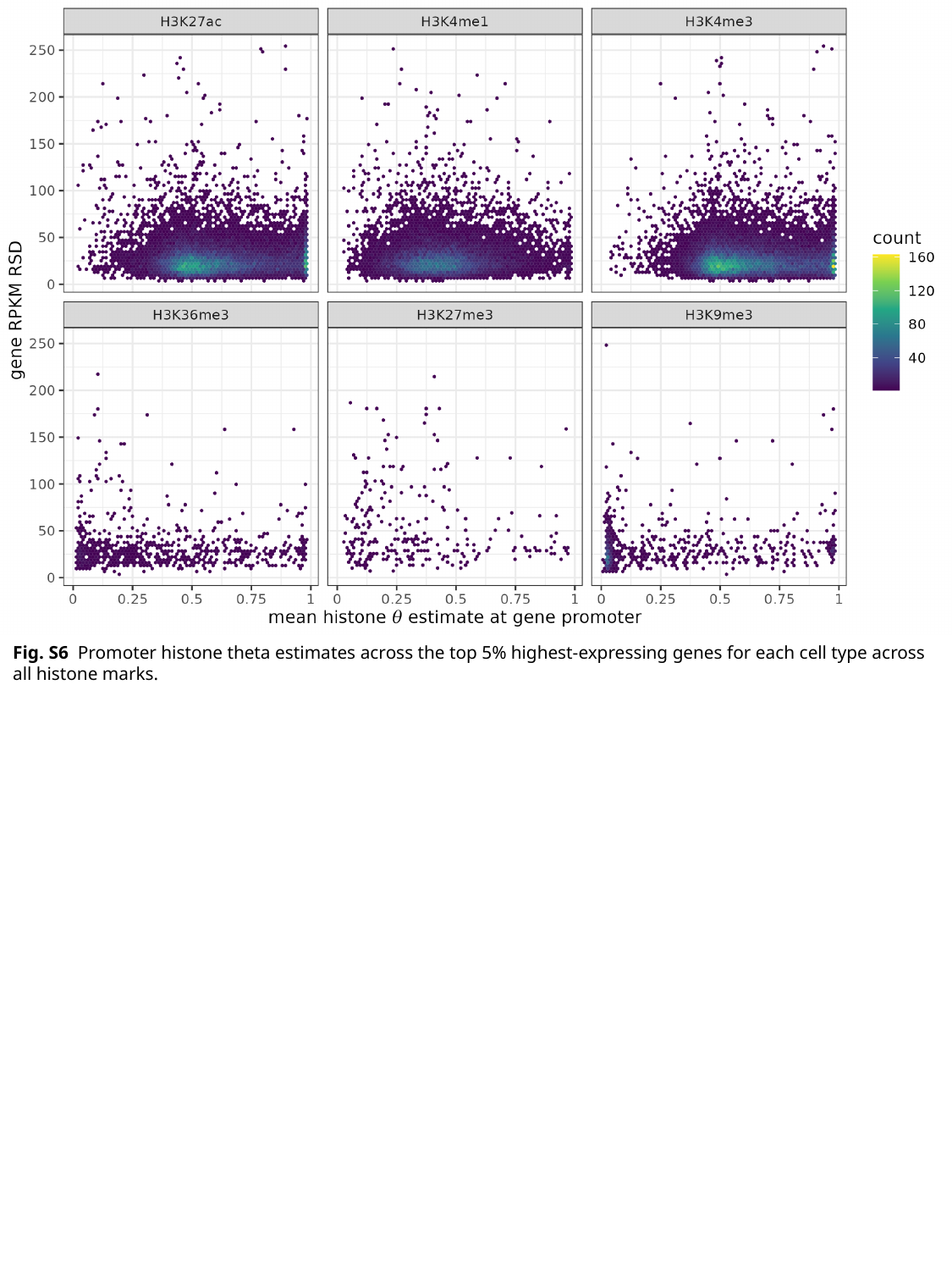

Fig. S6 Promoter histone theta estimates across the top 5% highest-expressing genes for each cell type across all histone marks.

### Slide 7
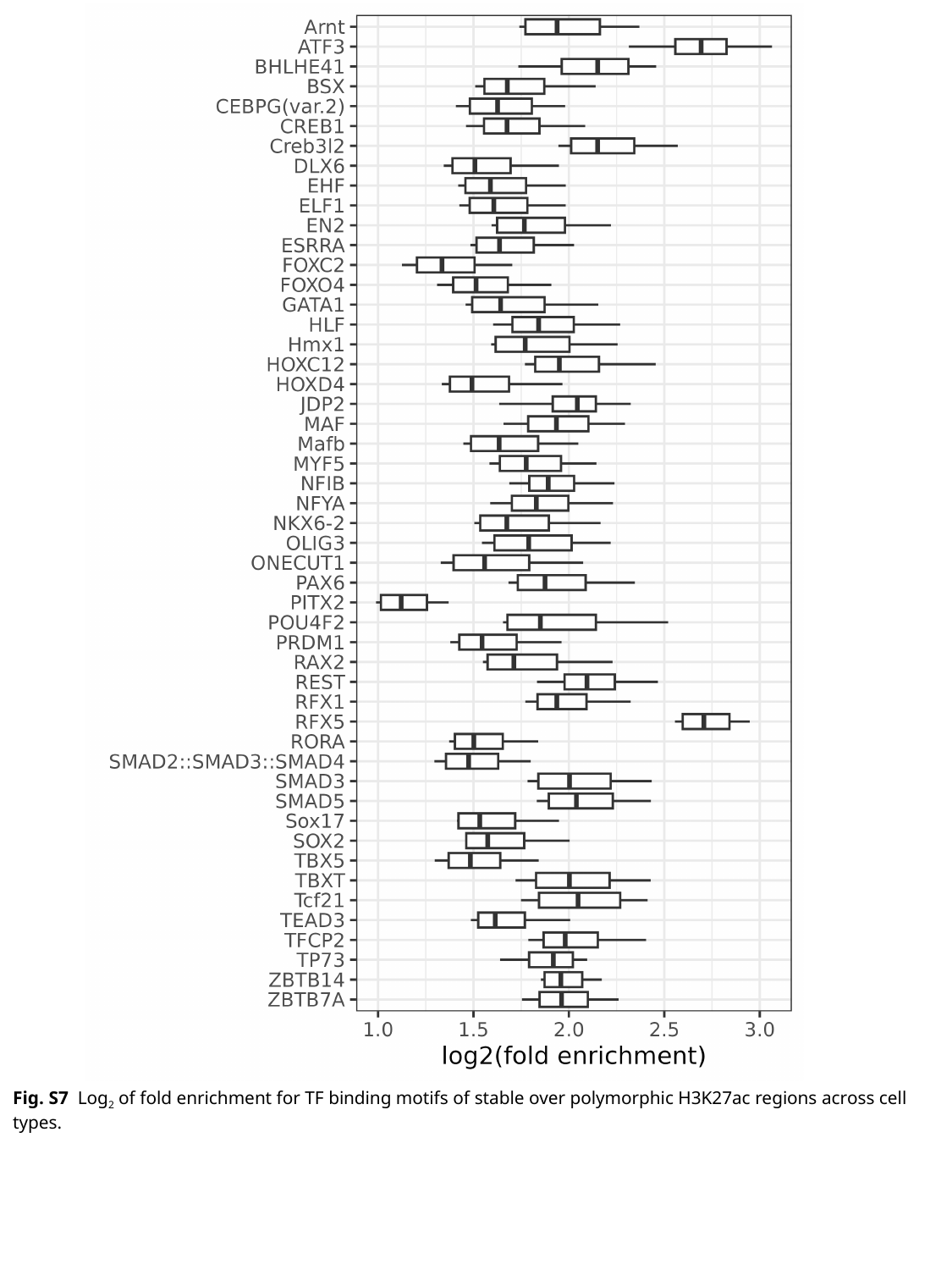

Fig. S7 Log_2_ of fold enrichment for TF binding motifs of stable over polymorphic H3K27ac regions across cell types.

### Slide 8
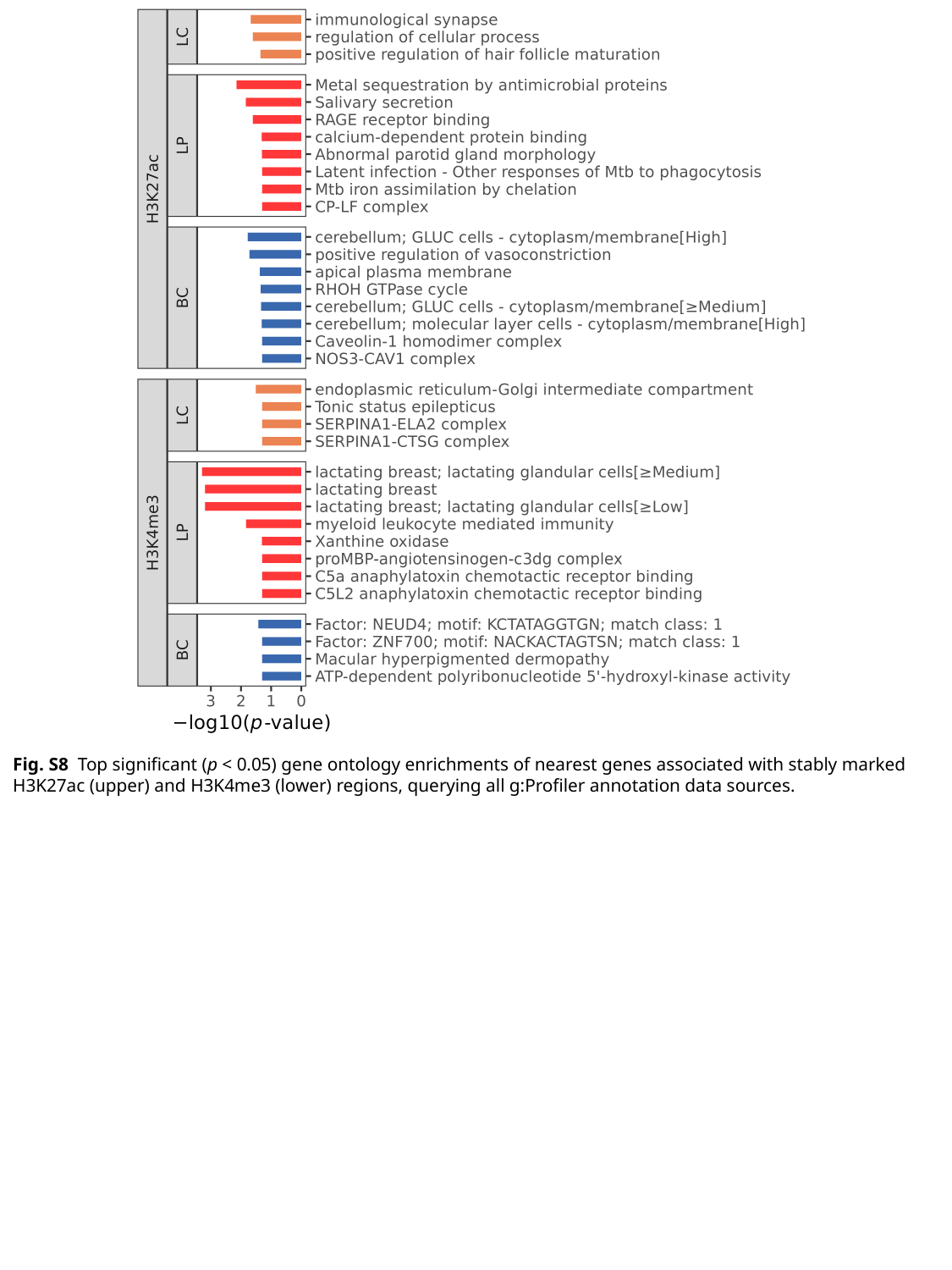

Fig. S8 Top significant (p < 0.05) gene ontology enrichments of nearest genes associated with stably marked H3K27ac (upper) and H3K4me3 (lower) regions, querying all g:Profiler annotation data sources.

### Slide 9
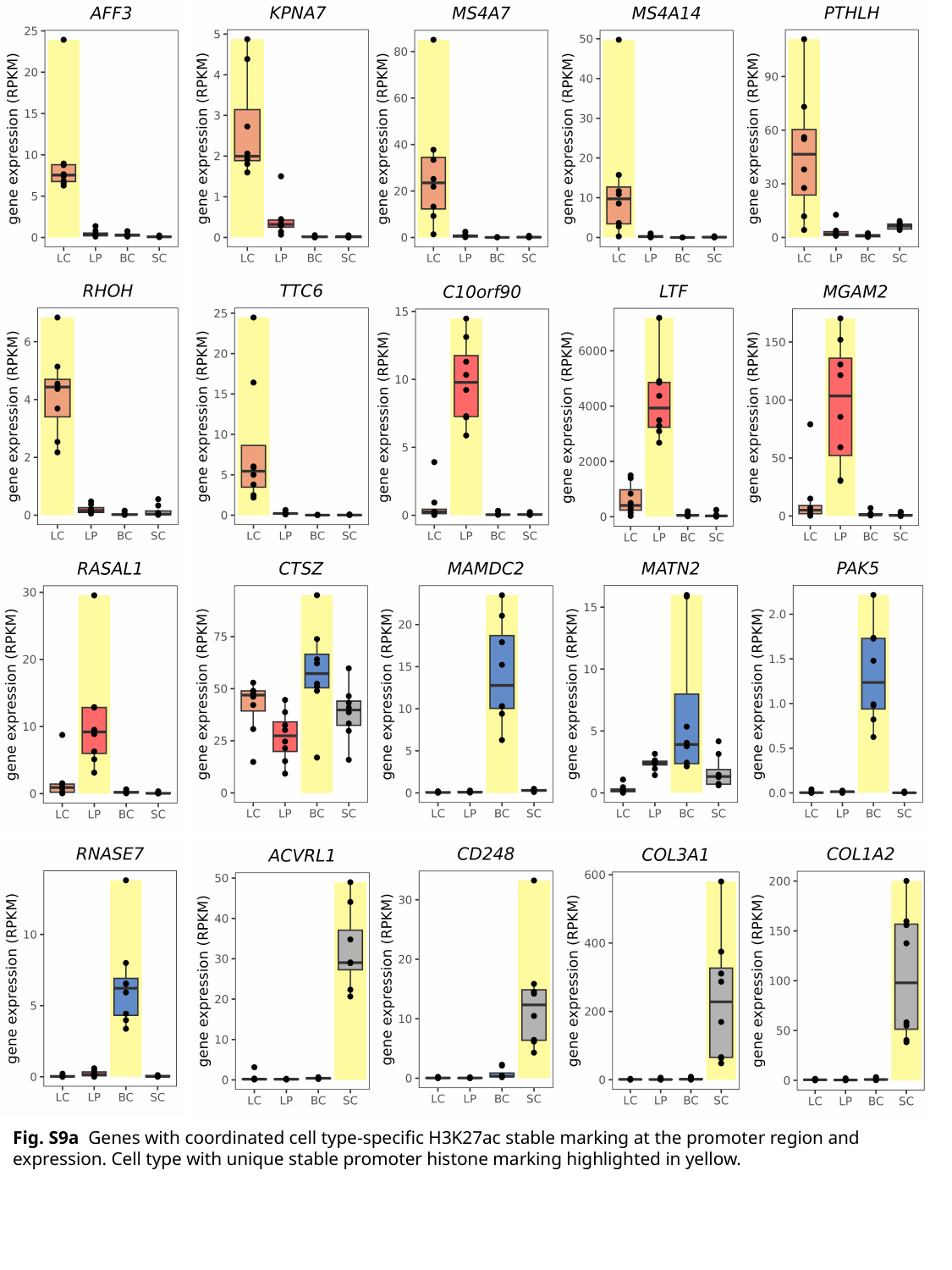

Fig. S9a Genes with coordinated cell type-specific H3K27ac stable marking at the promoter region and expression. Cell type with unique stable promoter histone marking highlighted in yellow.

### Slide 10
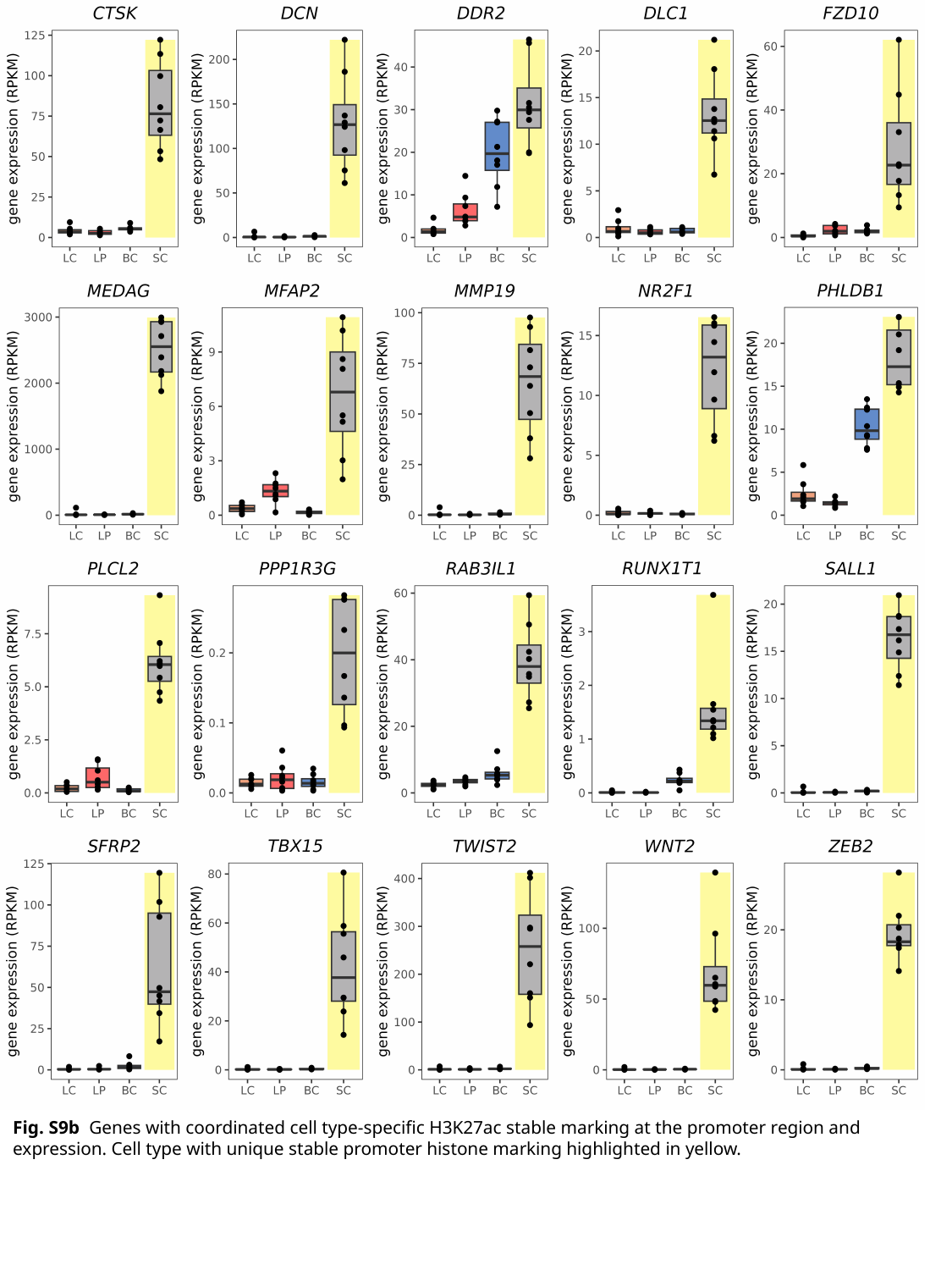

Fig. S9b Genes with coordinated cell type-specific H3K27ac stable marking at the promoter region and expression. Cell type with unique stable promoter histone marking highlighted in yellow.

### Slide 11
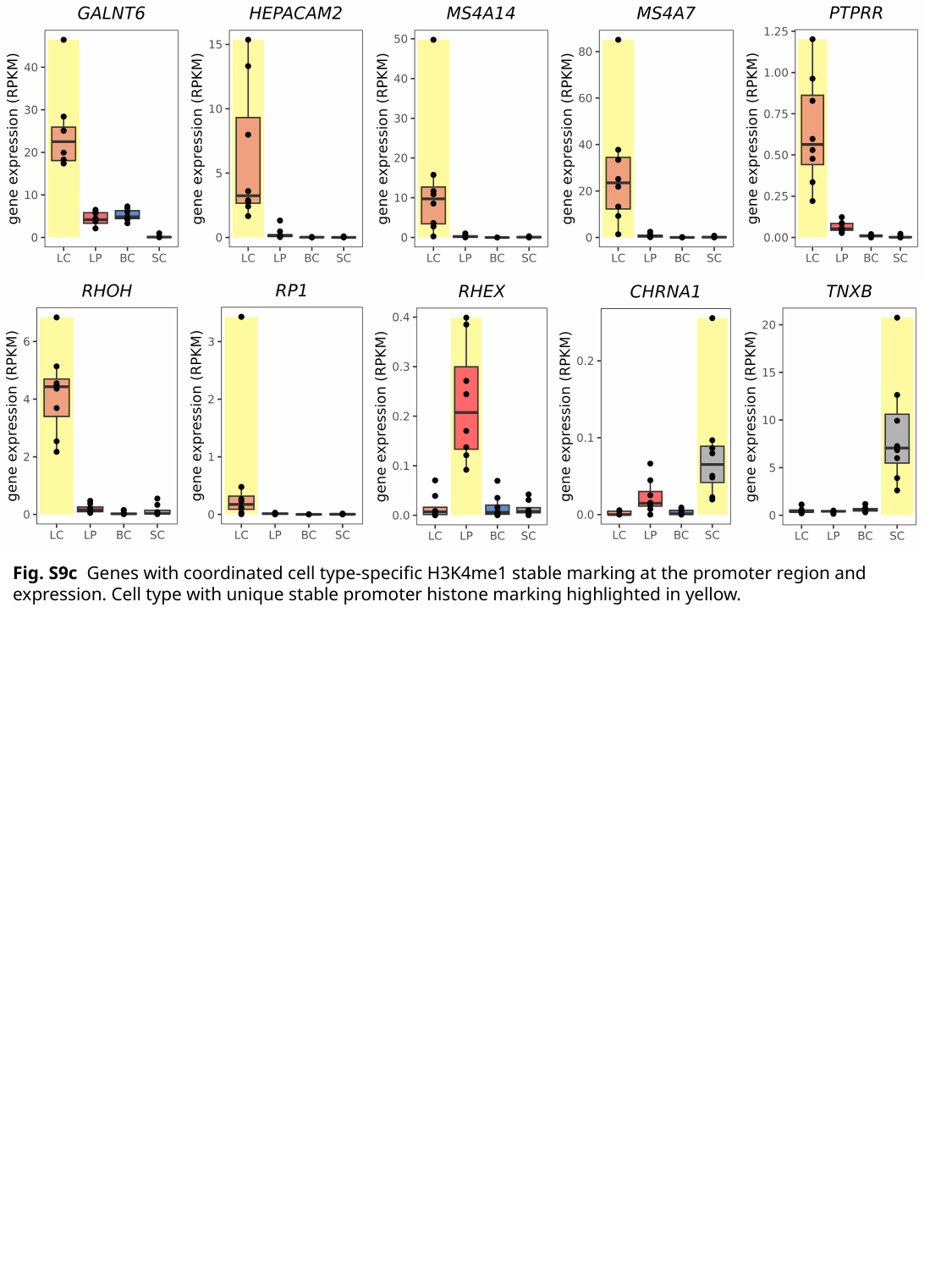

Fig. S9c Genes with coordinated cell type-specific H3K4me1 stable marking at the promoter region and expression. Cell type with unique stable promoter histone marking highlighted in yellow.

### Slide 12
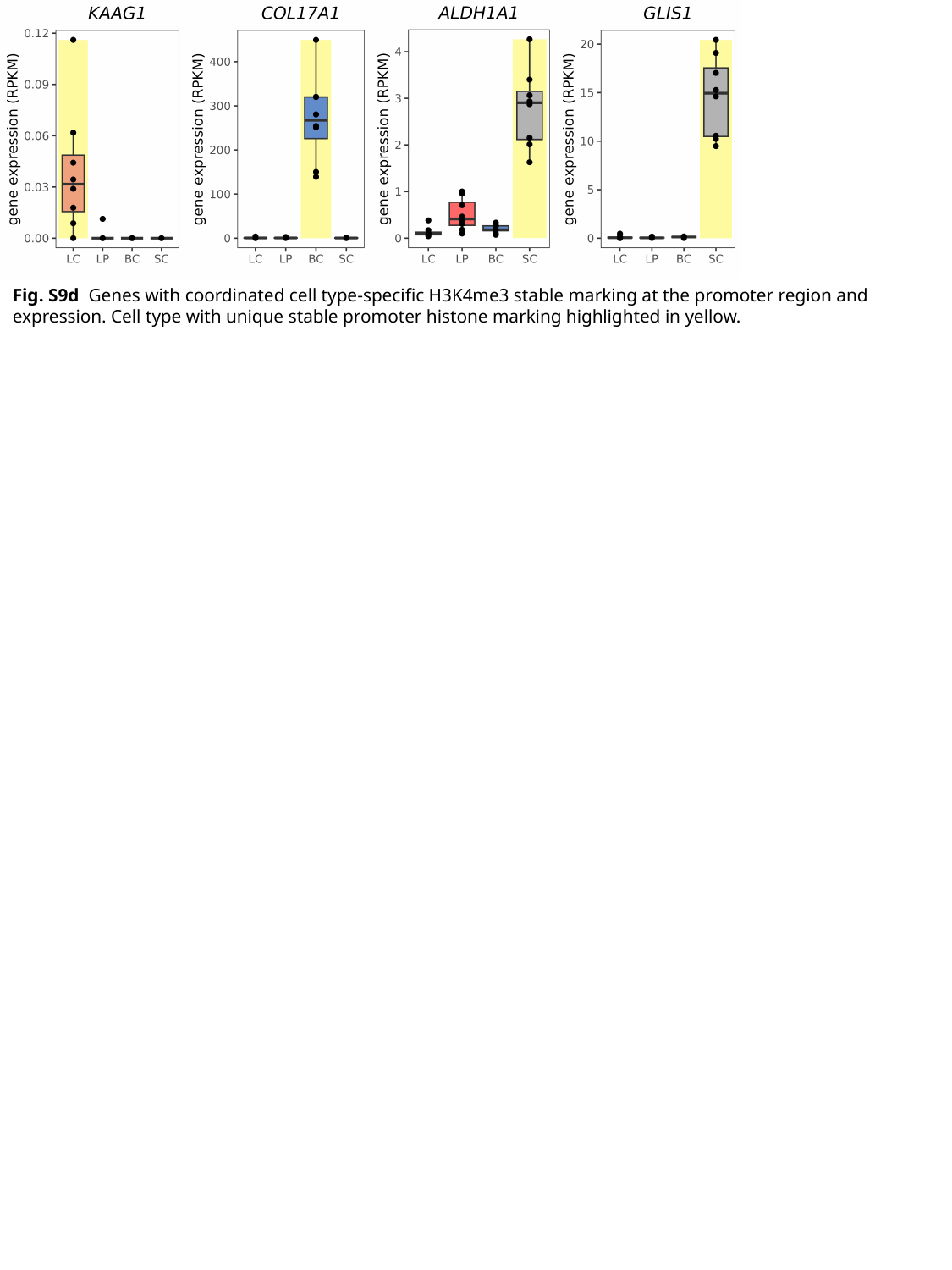

Fig. S9d Genes with coordinated cell type-specific H3K4me3 stable marking at the promoter region and expression. Cell type with unique stable promoter histone marking highlighted in yellow.

### Slide 13
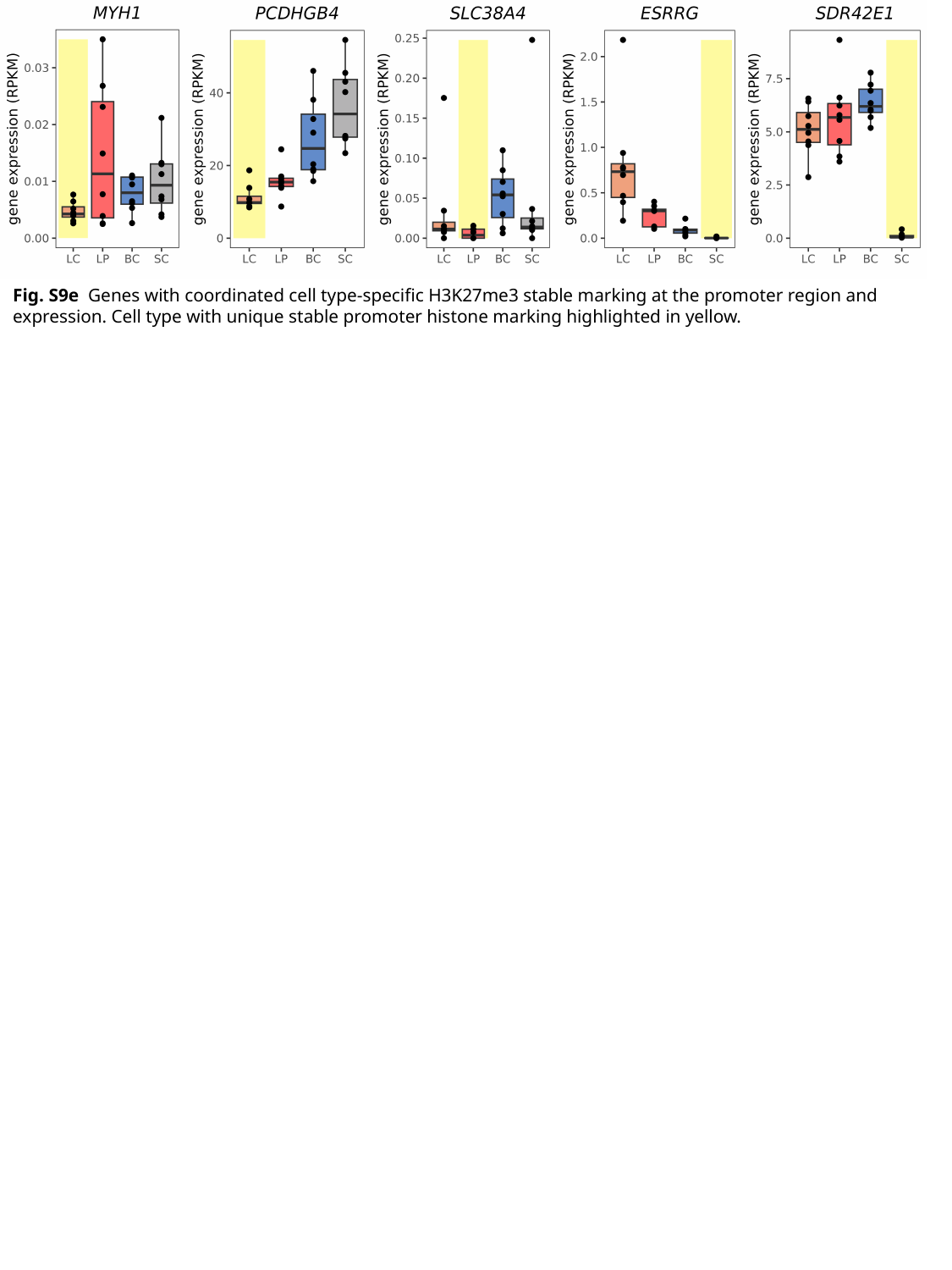

Fig. S9e Genes with coordinated cell type-specific H3K27me3 stable marking at the promoter region and expression. Cell type with unique stable promoter histone marking highlighted in yellow.

### Slide 14
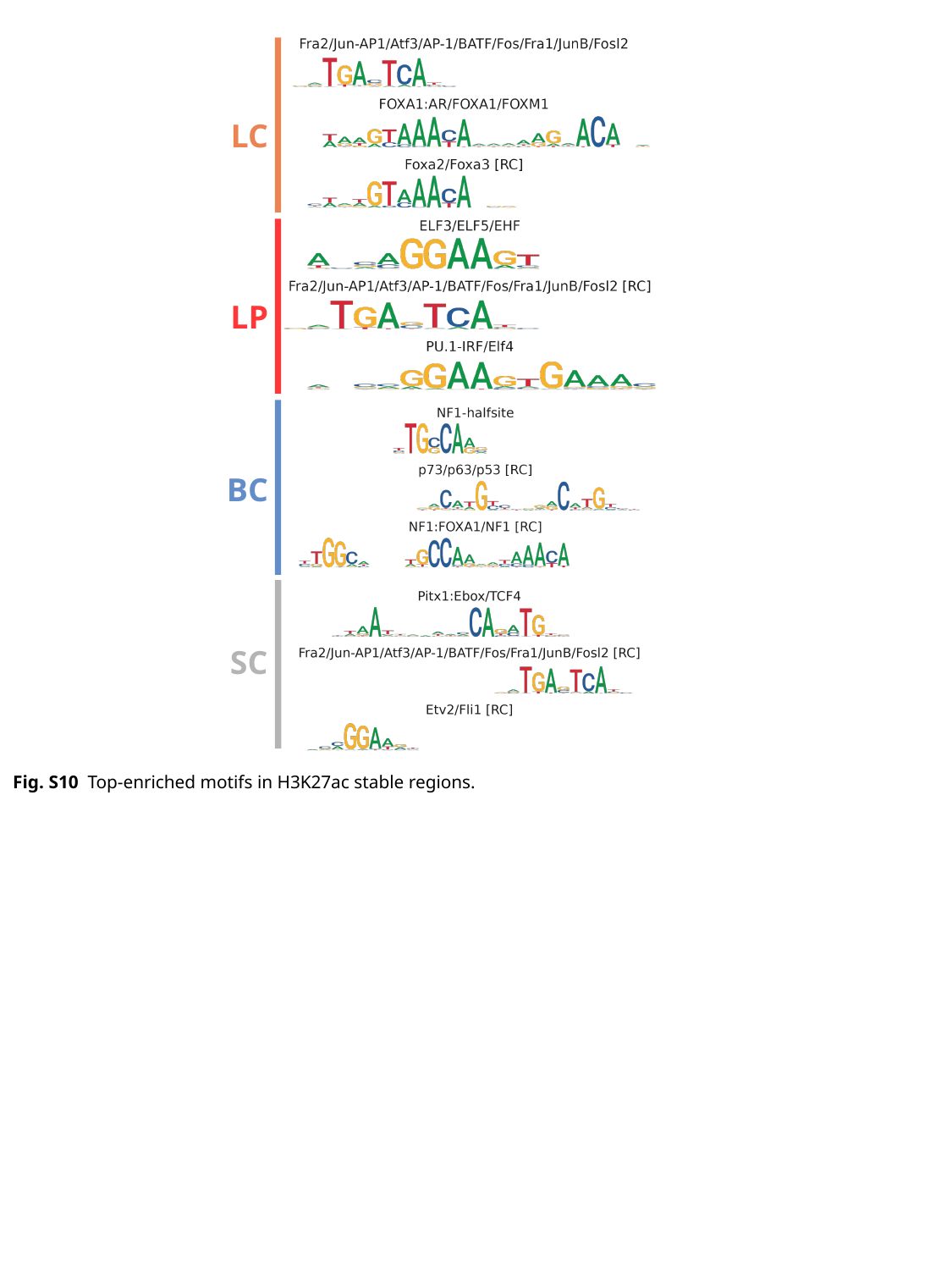

LC
LP
BC
SC
Fig. S10 Top-enriched motifs in H3K27ac stable regions.

### Slide 15
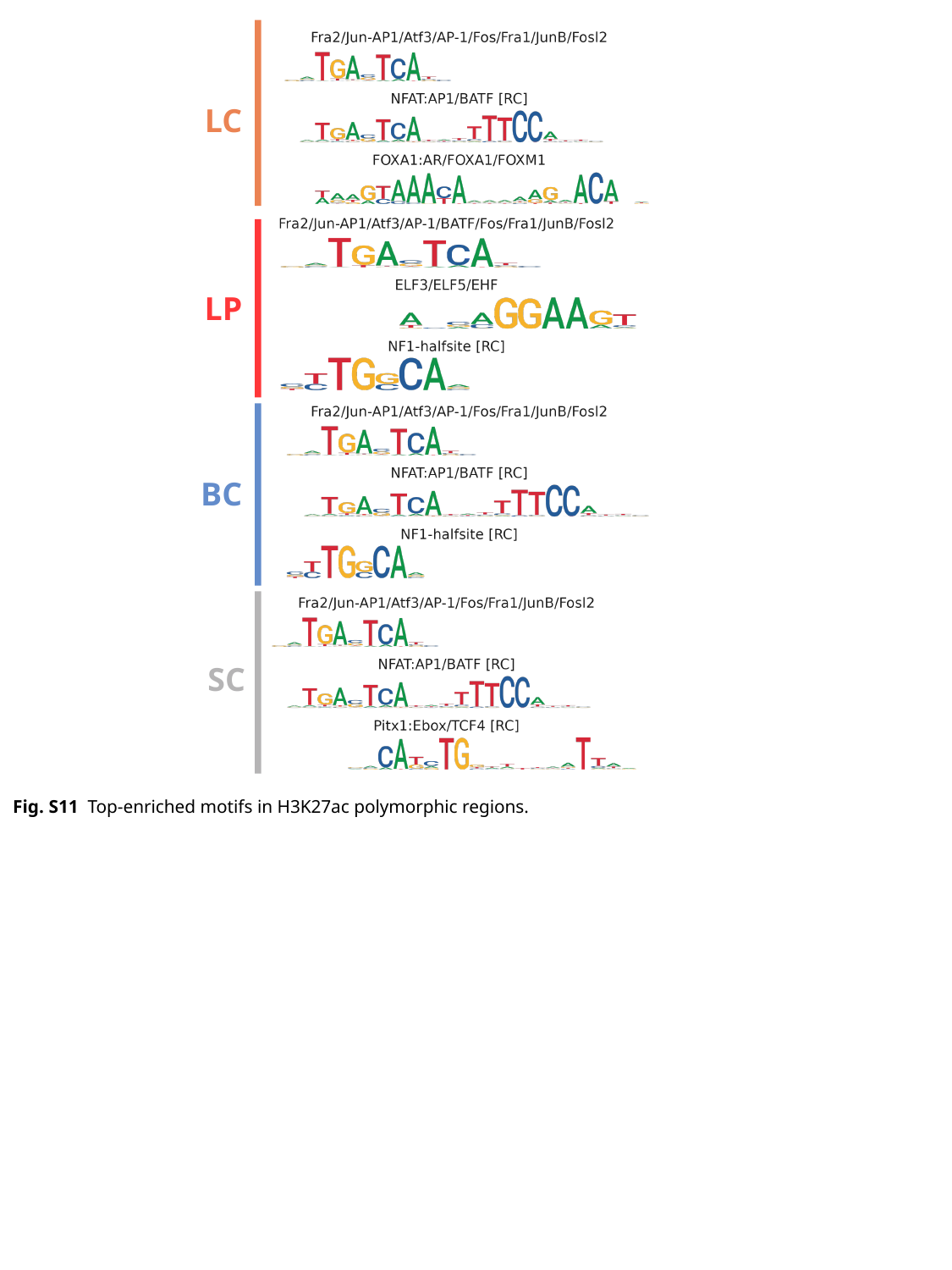

LC
LP
BC
SC
Fig. S11 Top-enriched motifs in H3K27ac polymorphic regions.

### Slide 16
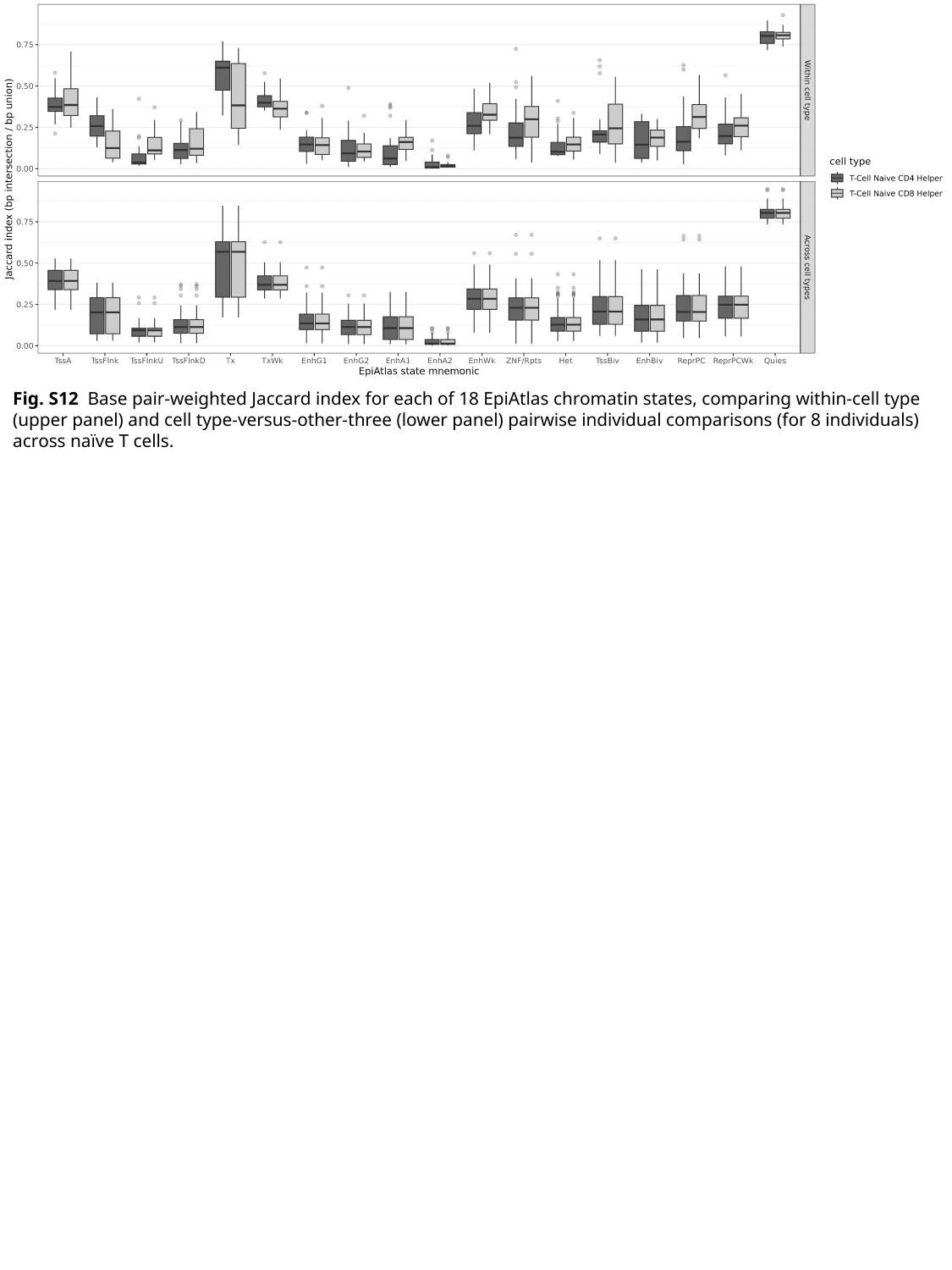

Fig. S12 Base pair-weighted Jaccard index for each of 18 EpiAtlas chromatin states, comparing within-cell type (upper panel) and cell type-versus-other-three (lower panel) pairwise individual comparisons (for 8 individuals) across naïve T cells.

### Slide 17
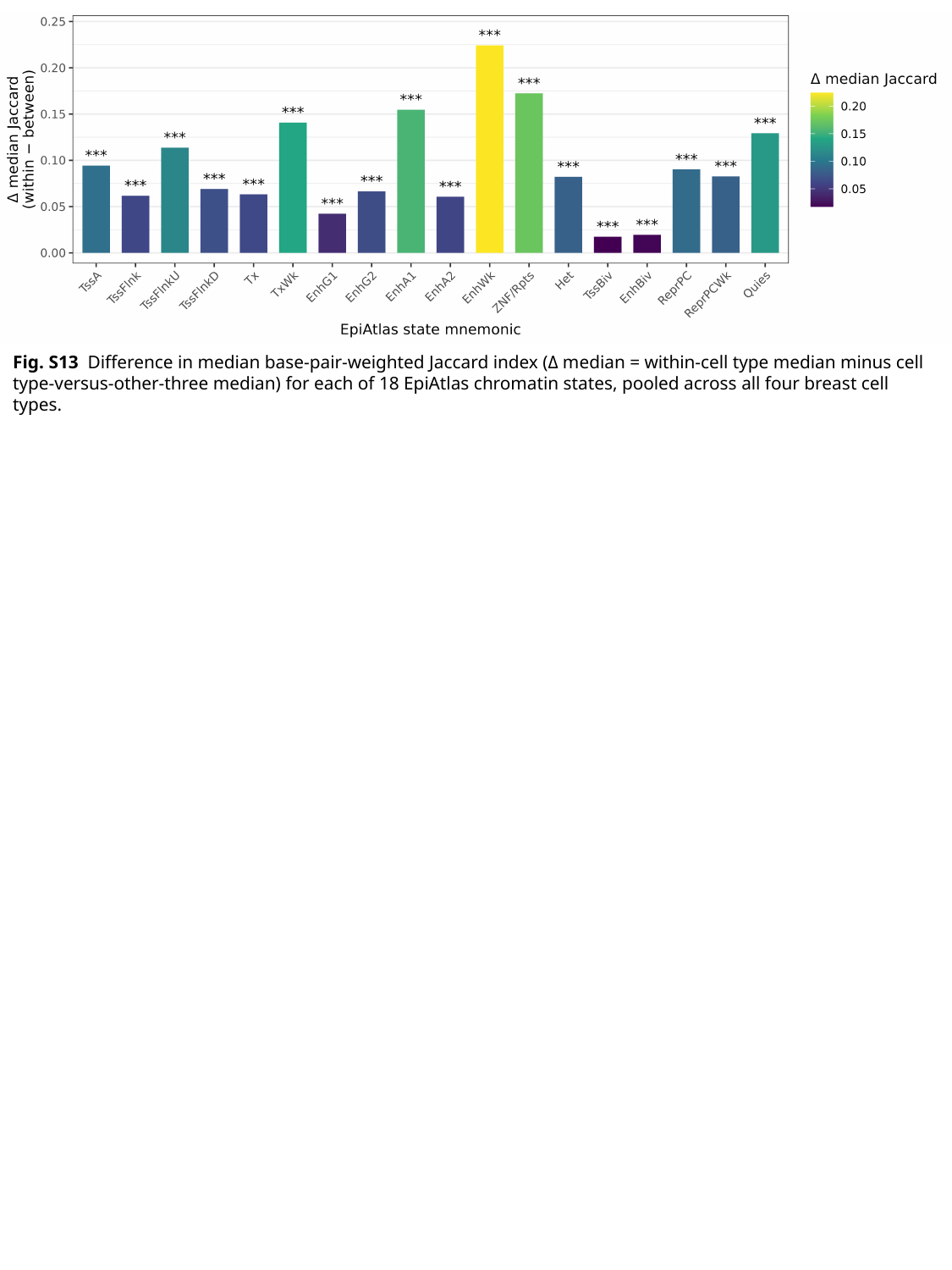

Fig. S13 Difference in median base-pair-weighted Jaccard index (Δ median = within-cell type median minus cell type-versus-other-three median) for each of 18 EpiAtlas chromatin states, pooled across all four breast cell types.
Fig. S13c Bar plot showing the Kruskal–Wallis χ² statistic for each of 18 EpiAtlas chromatin states, testing whether the distribution of within-cell-type Jaccard indices differs among the four breast cell types (LC, LP, BC, SC). Bar color encodes −log10(BH-adjusted p-value). The strongest cell type effects were observed for EnhA2 (χ² = 29.1, p < 0.001), ReprPCWk (χ² = 23.0, p < 0.001), and EnhA1 (χ² = 21.3, p < 0.001). Post-hoc pairwise Wilcoxon tests identified SCs as the primary outlier, with significantly lower enhancer state concordance than the epithelial cell types.

### Slide 18
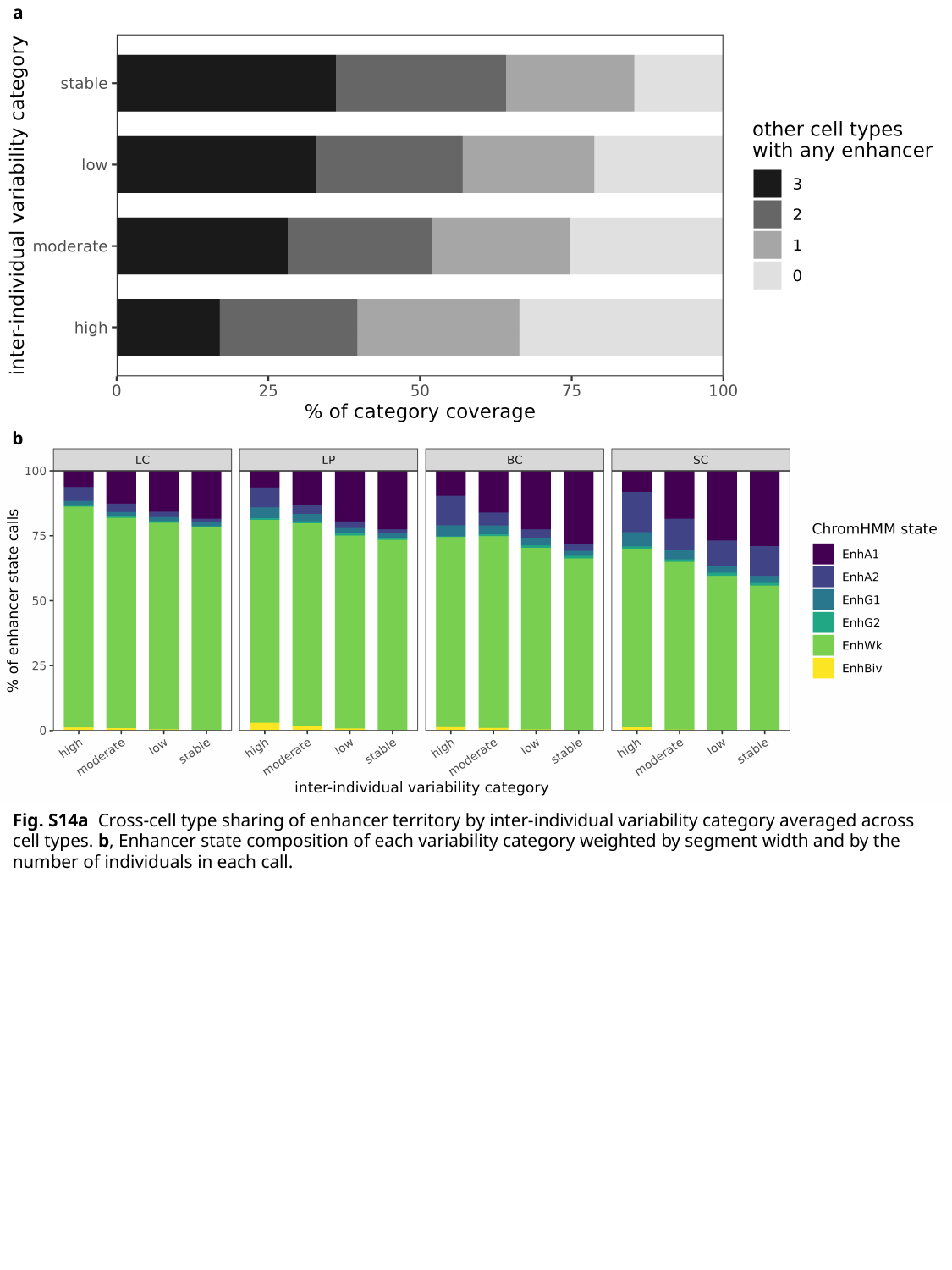

a
b
Fig. S14a Cross-cell type sharing of enhancer territory by inter-individual variability category averaged across cell types. b, Enhancer state composition of each variability category weighted by segment width and by the number of individuals in each call.

### Slide 19
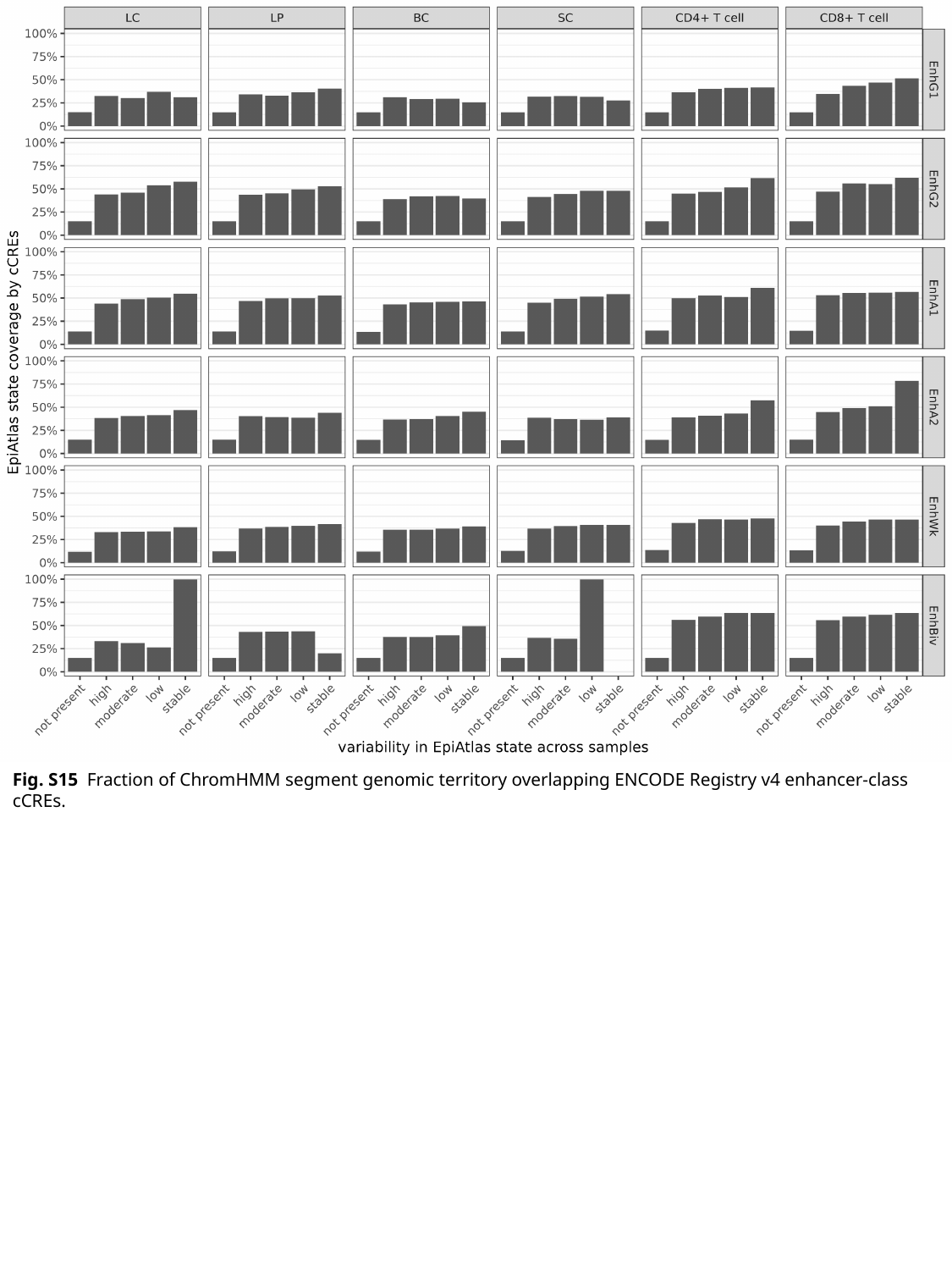

Fig. S15 Fraction of ChromHMM segment genomic territory overlapping ENCODE Registry v4 enhancer-class cCREs.

### Slide 20
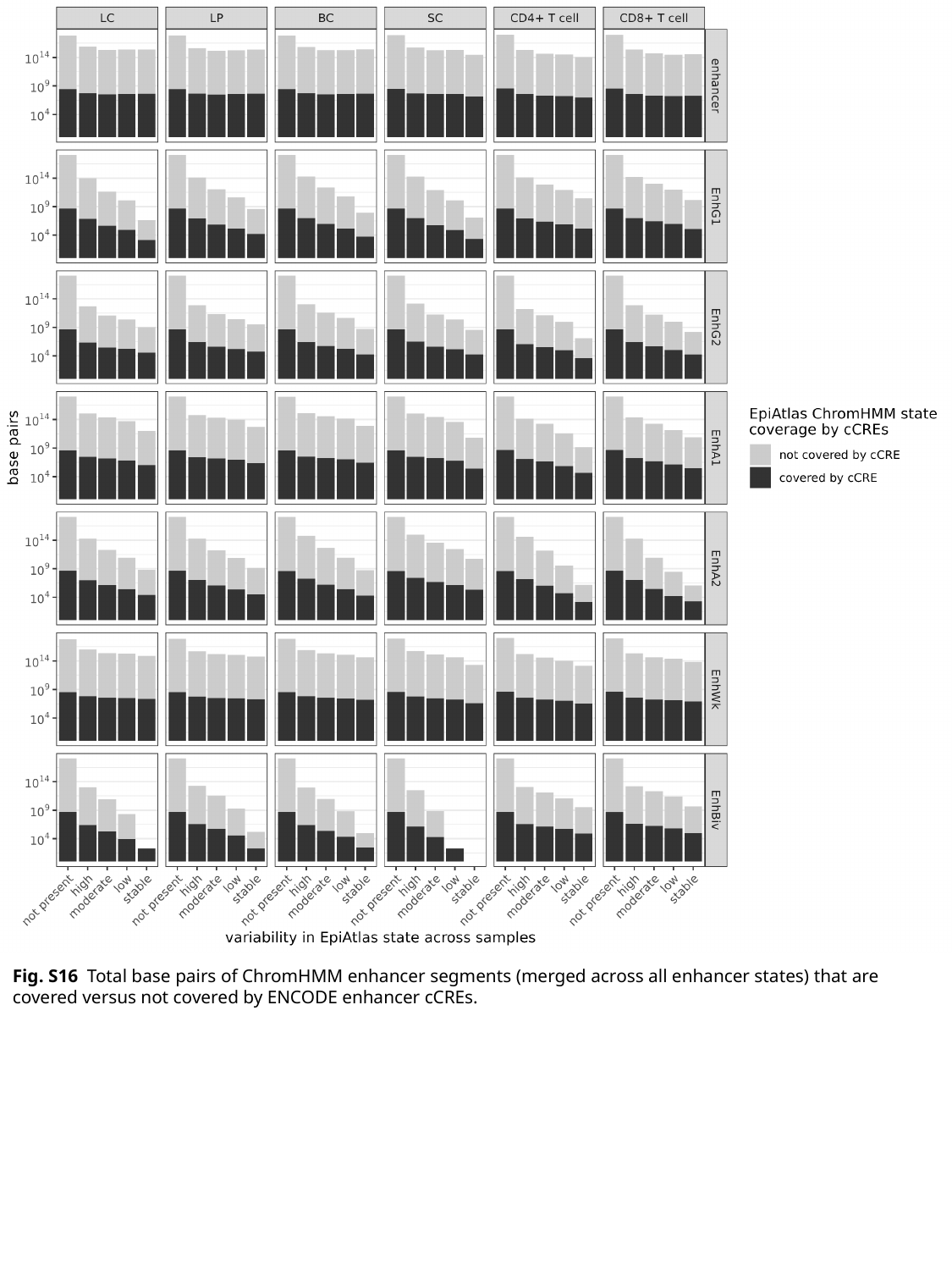

Fig. S16 Total base pairs of ChromHMM enhancer segments (merged across all enhancer states) that are covered versus not covered by ENCODE enhancer cCREs.

### Slide 21
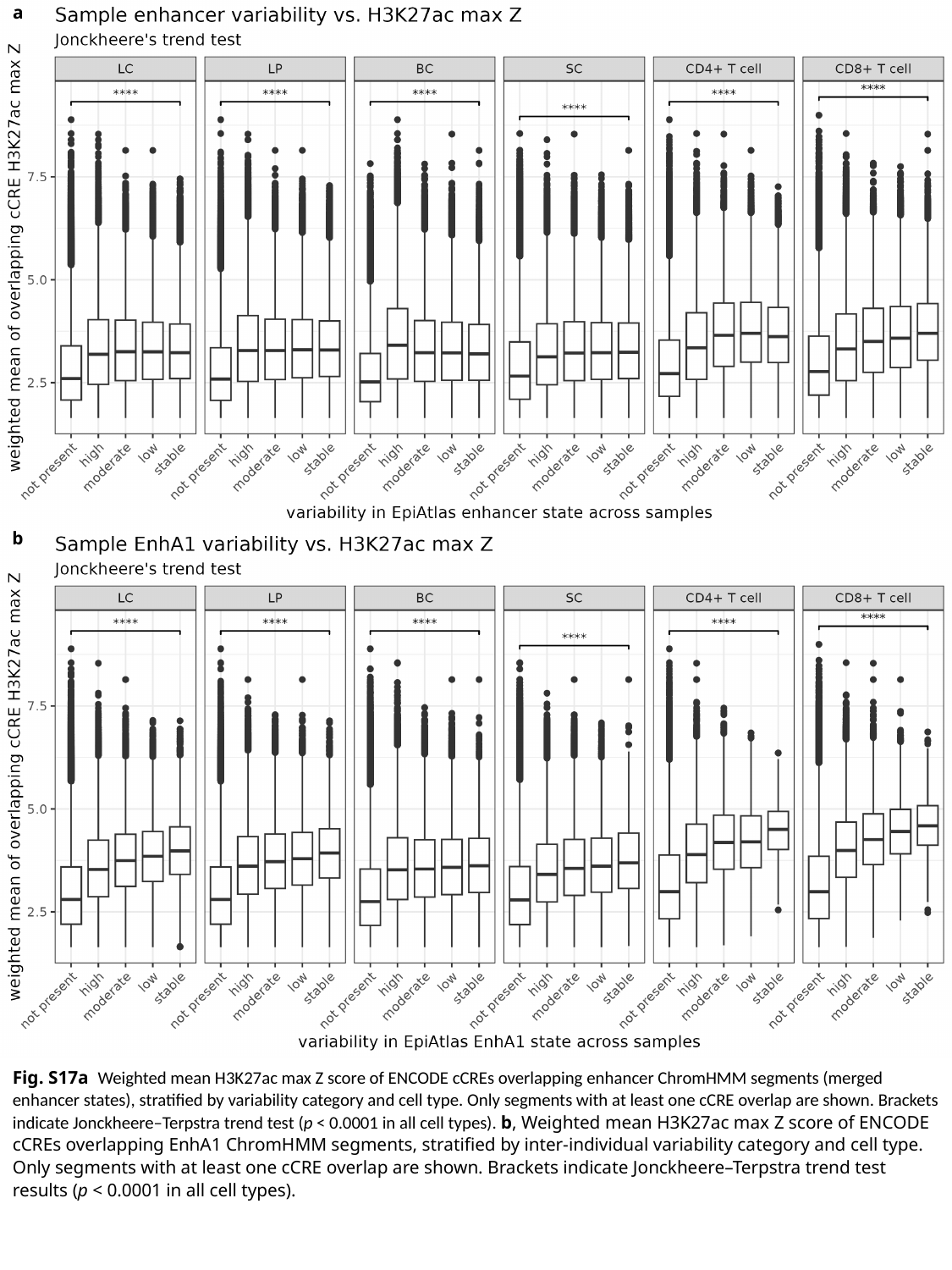

a
b
Fig. S17a Weighted mean H3K27ac max Z score of ENCODE cCREs overlapping enhancer ChromHMM segments (merged enhancer states), stratified by variability category and cell type. Only segments with at least one cCRE overlap are shown. Brackets indicate Jonckheere–Terpstra trend test (p < 0.0001 in all cell types). b, Weighted mean H3K27ac max Z score of ENCODE cCREs overlapping EnhA1 ChromHMM segments, stratified by inter-individual variability category and cell type. Only segments with at least one cCRE overlap are shown. Brackets indicate Jonckheere–Terpstra trend test results (p < 0.0001 in all cell types).

### Slide 22
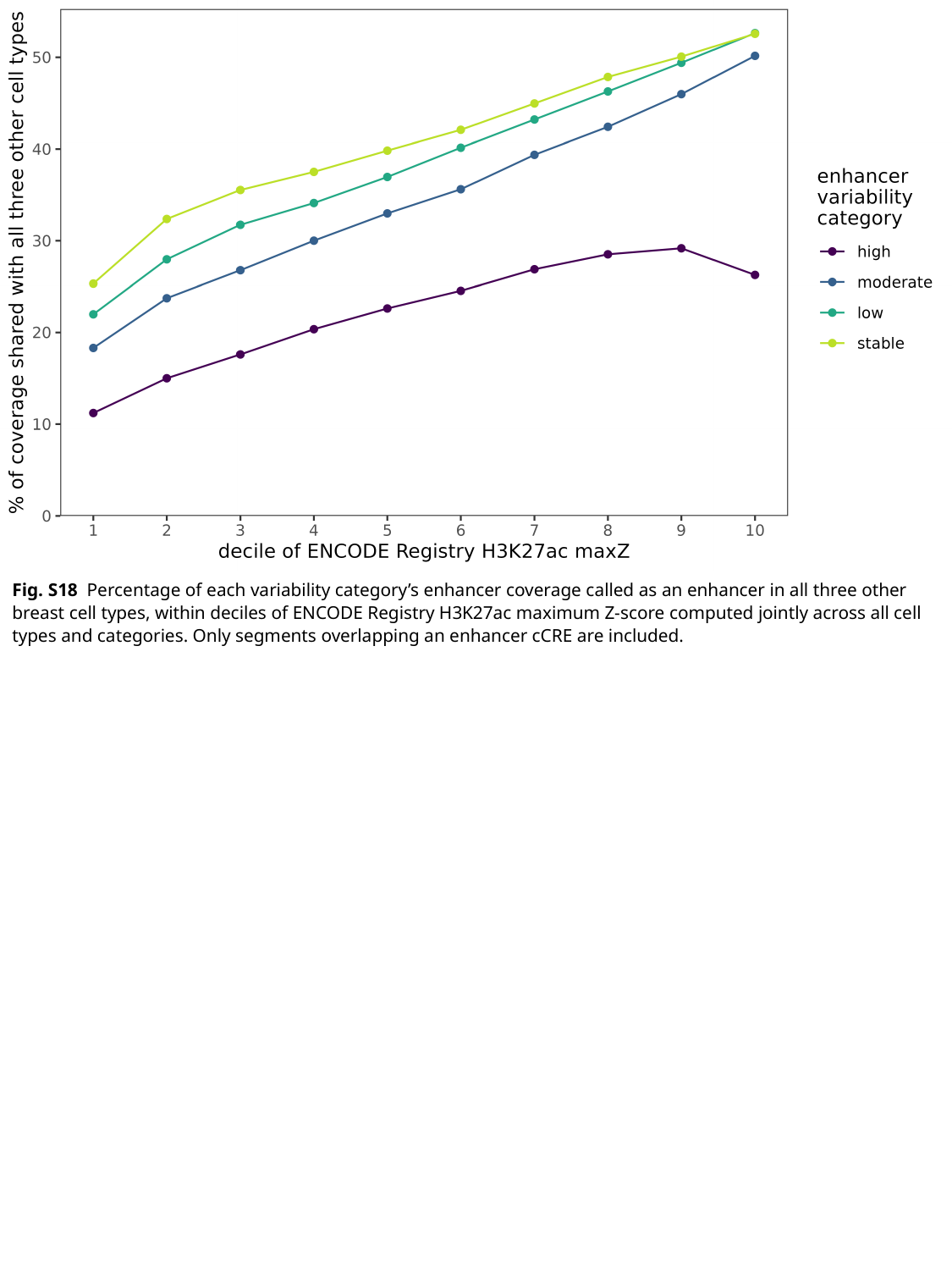

Fig. S18 Percentage of each variability category’s enhancer coverage called as an enhancer in all three other breast cell types, within deciles of ENCODE Registry H3K27ac maximum Z-score computed jointly across all cell types and categories. Only segments overlapping an enhancer cCRE are included.

### Slide 23
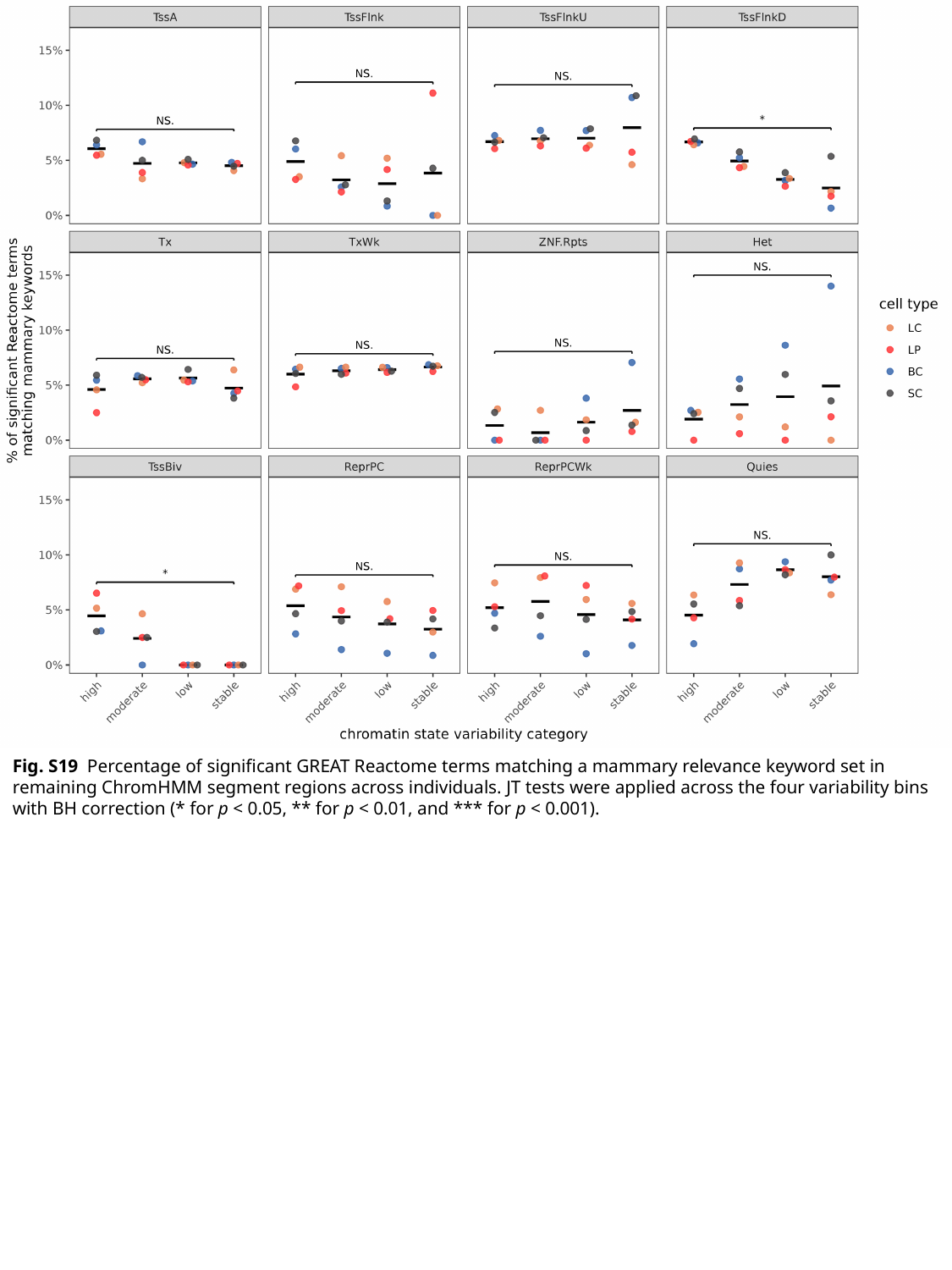

Fig. S19 Percentage of significant GREAT Reactome terms matching a mammary relevance keyword set in remaining ChromHMM segment regions across individuals. JT tests were applied across the four variability bins with BH correction (* for p < 0.05, ** for p < 0.01, and *** for p < 0.001).

### Slide 24
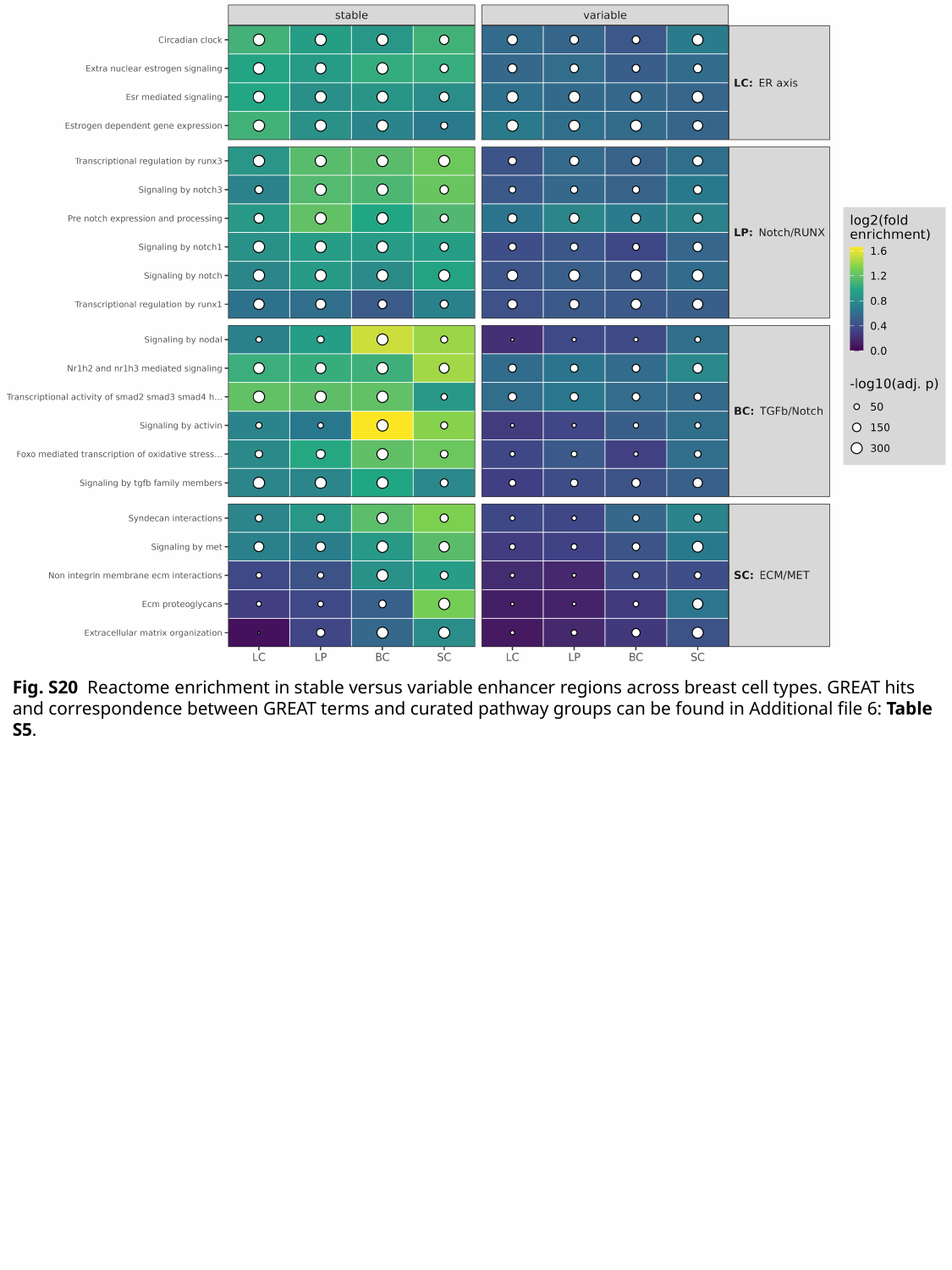

Fig. S20 Reactome enrichment in stable versus variable enhancer regions across breast cell types. GREAT hits and correspondence between GREAT terms and curated pathway groups can be found in Additional file 6: Table S5.

### Slide 25
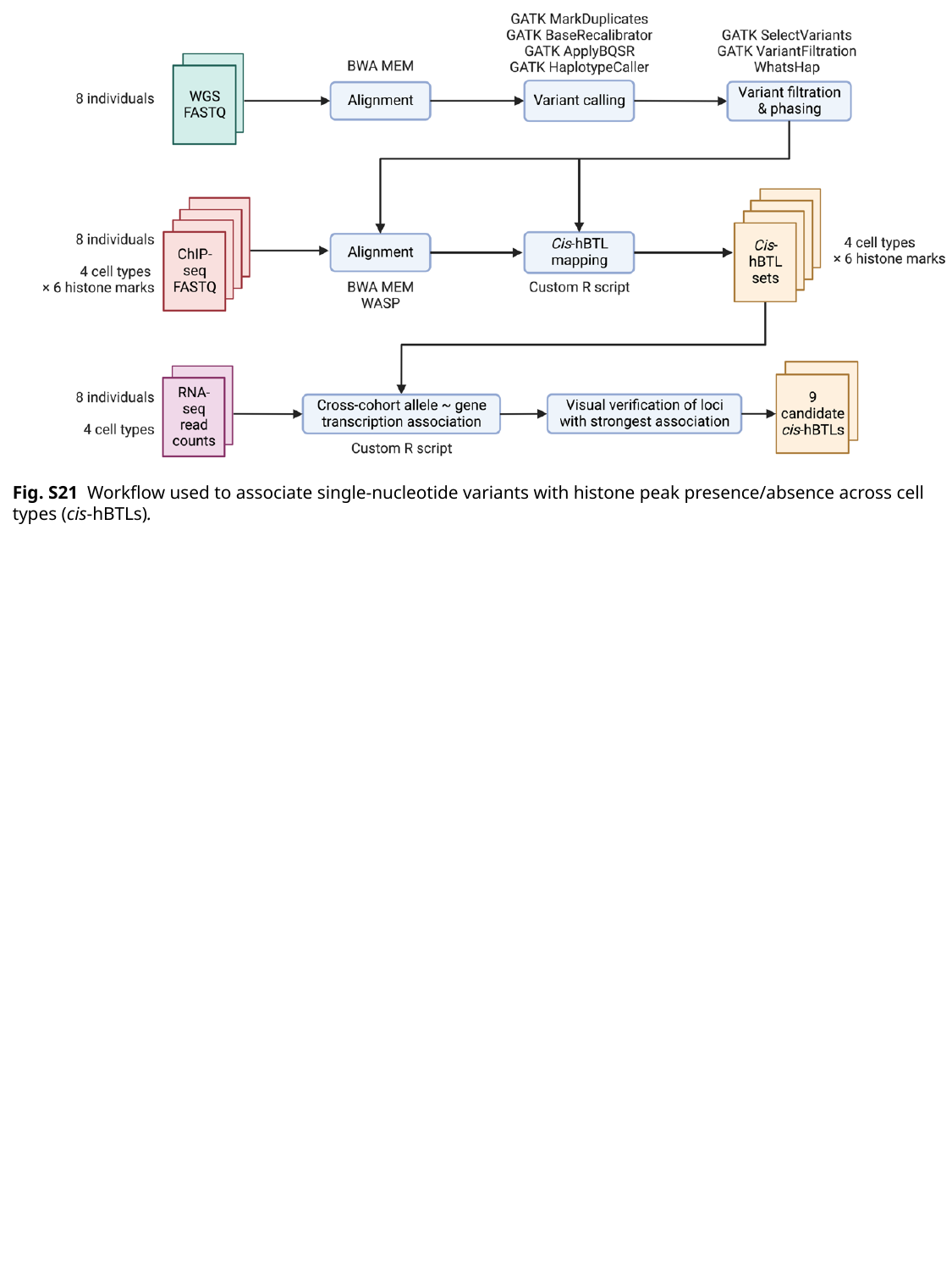

Fig. S21 Workflow used to associate single-nucleotide variants with histone peak presence/absence across cell types (cis-hBTLs).

### Slide 26
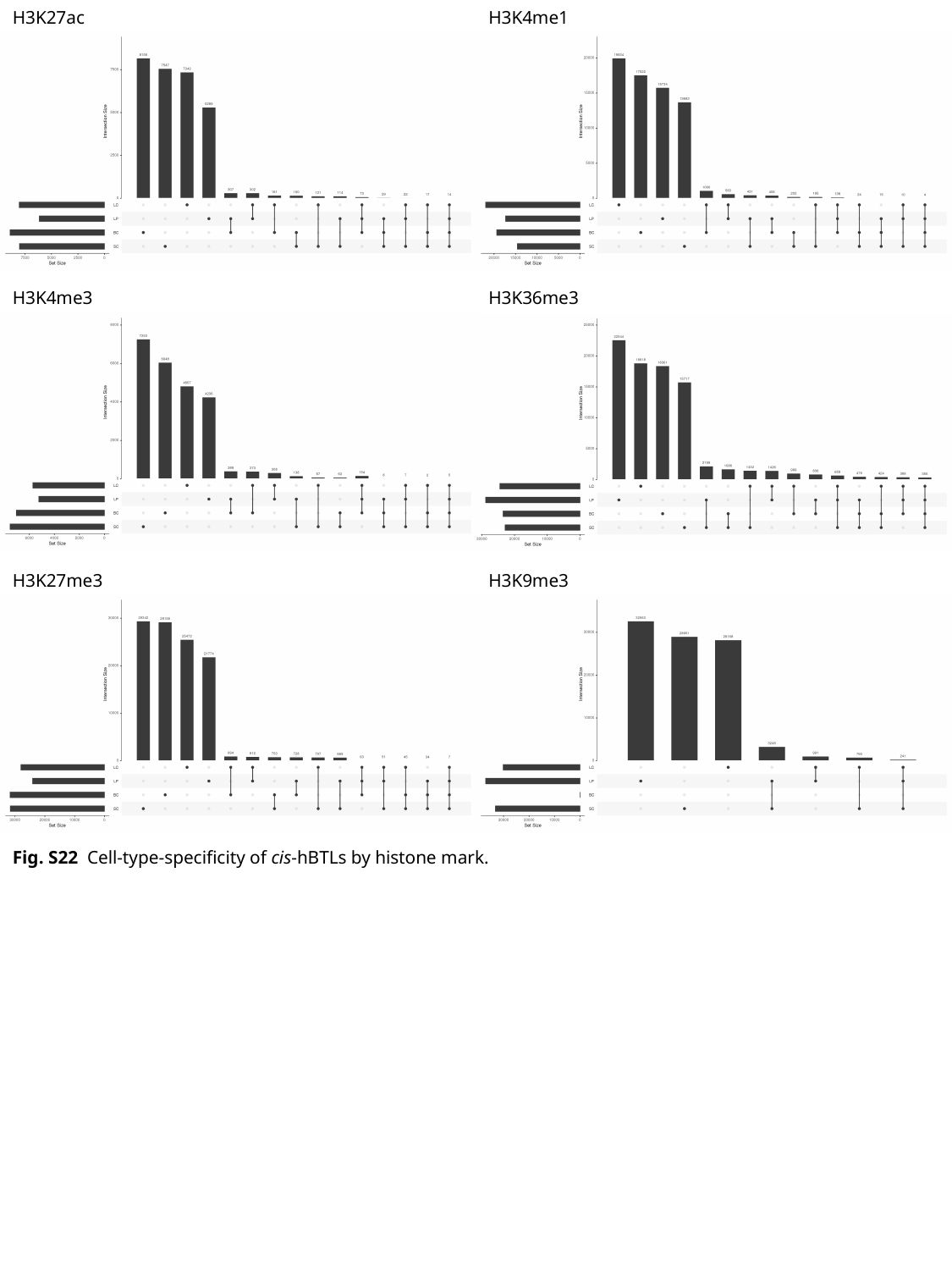

H3K27ac
H3K4me1
H3K4me3
H3K36me3
H3K27me3
H3K9me3
Fig. S22 Cell-type-specificity of cis-hBTLs by histone mark.

### Slide 27
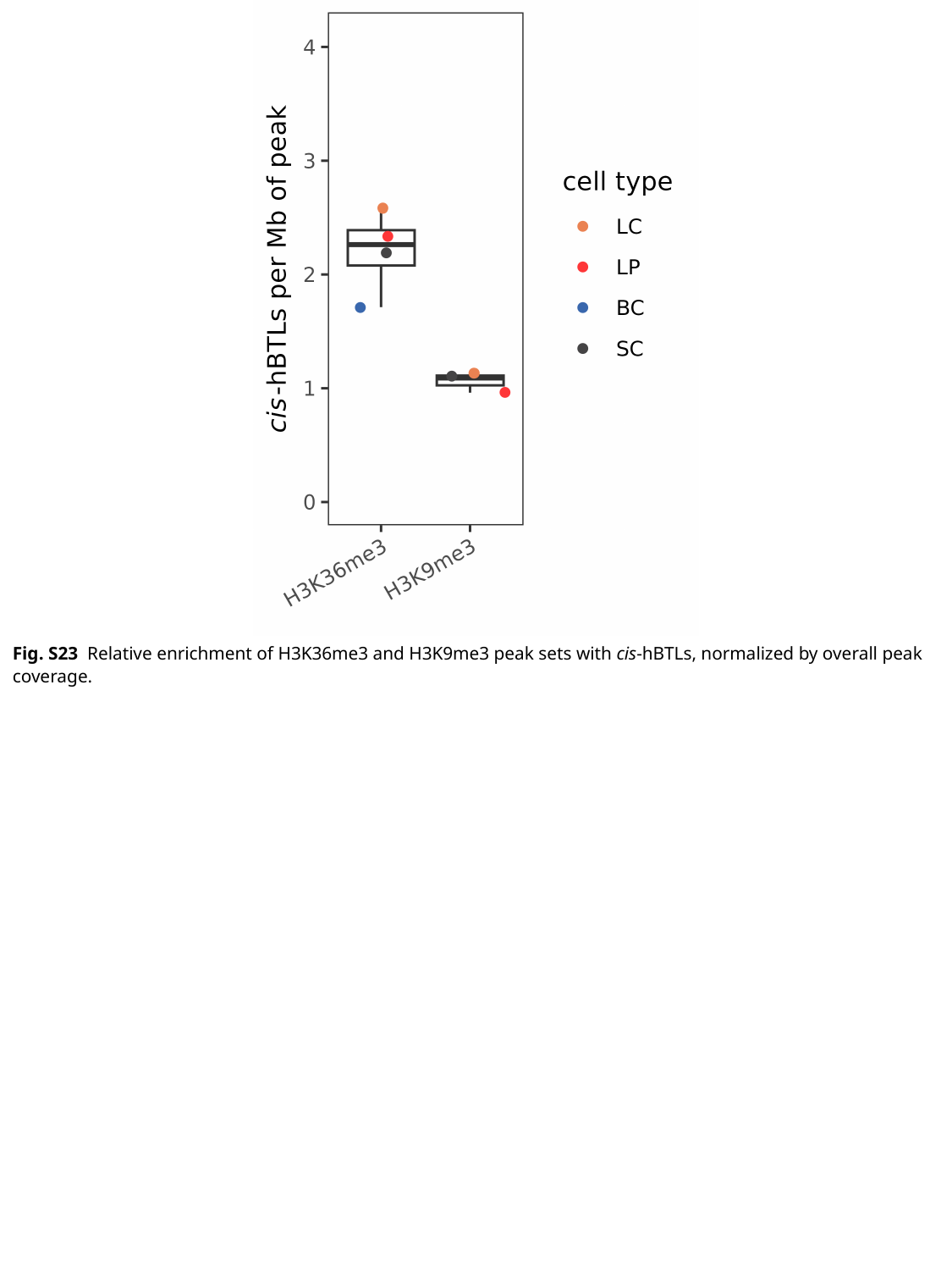

Fig. S23 Relative enrichment of H3K36me3 and H3K9me3 peak sets with cis-hBTLs, normalized by overall peak coverage.

### Slide 28
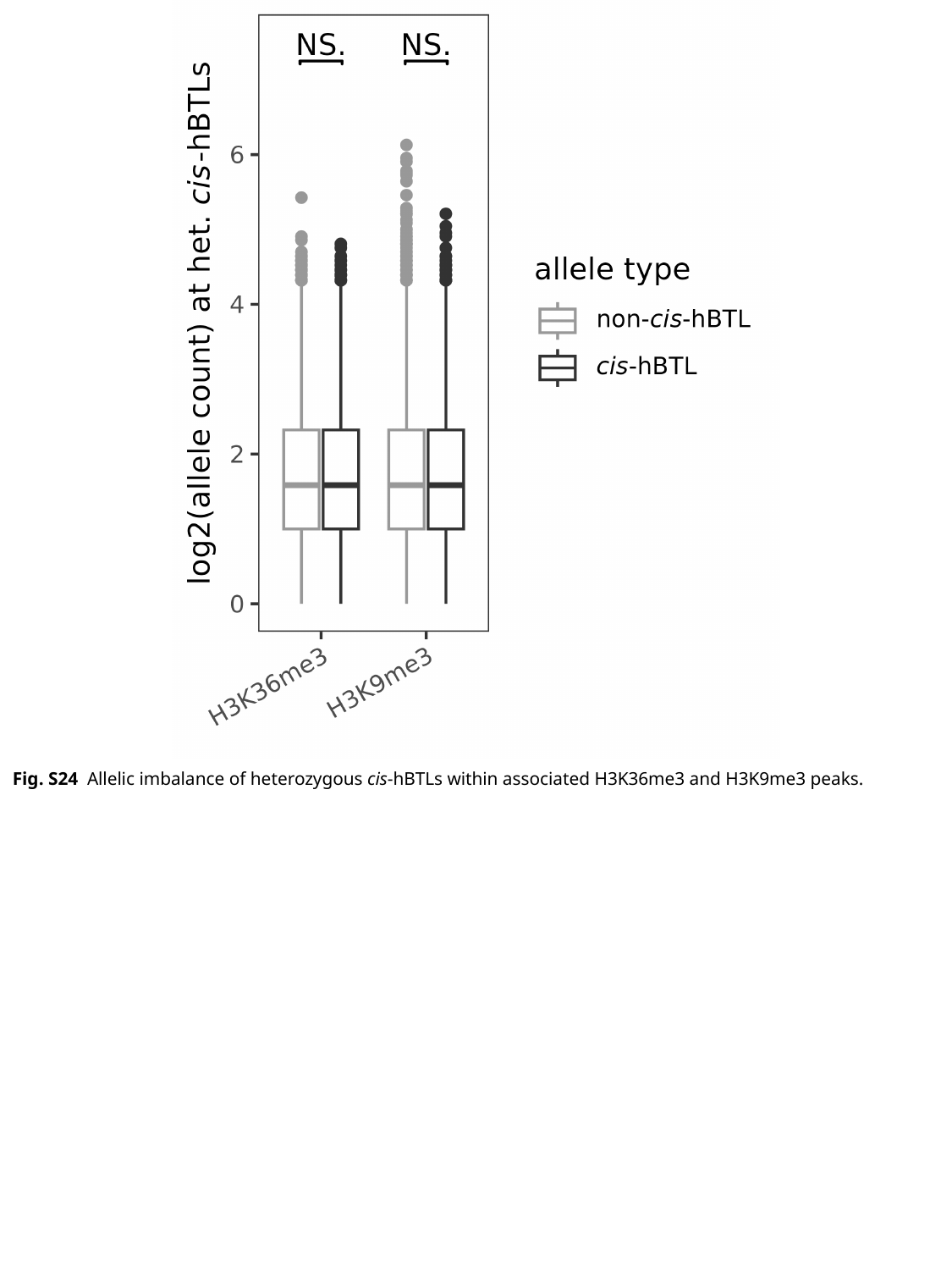

Fig. S24 Allelic imbalance of heterozygous cis-hBTLs within associated H3K36me3 and H3K9me3 peaks.

### Slide 29
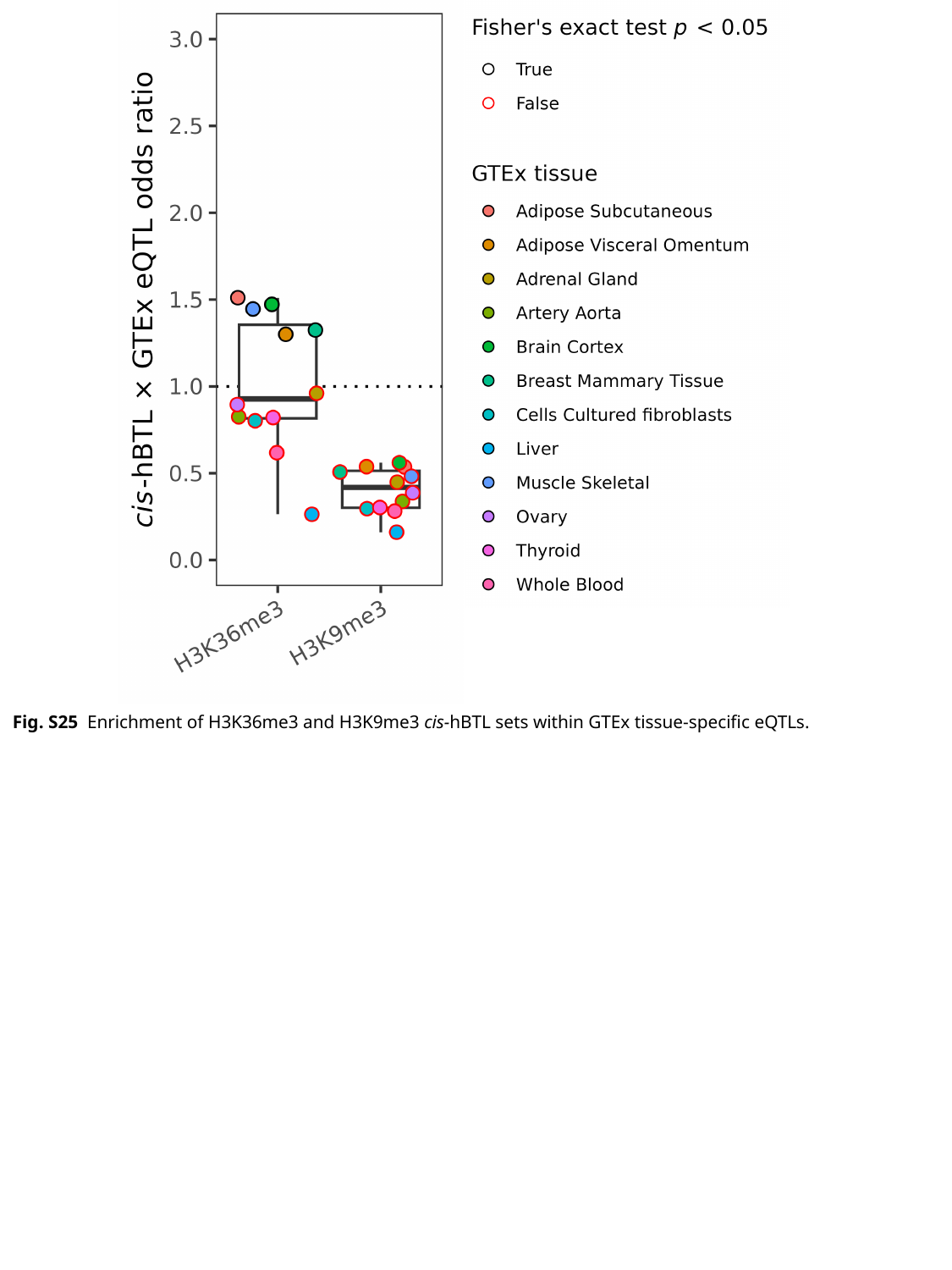

Fig. S25 Enrichment of H3K36me3 and H3K9me3 cis-hBTL sets within GTEx tissue-specific eQTLs.

### Slide 30
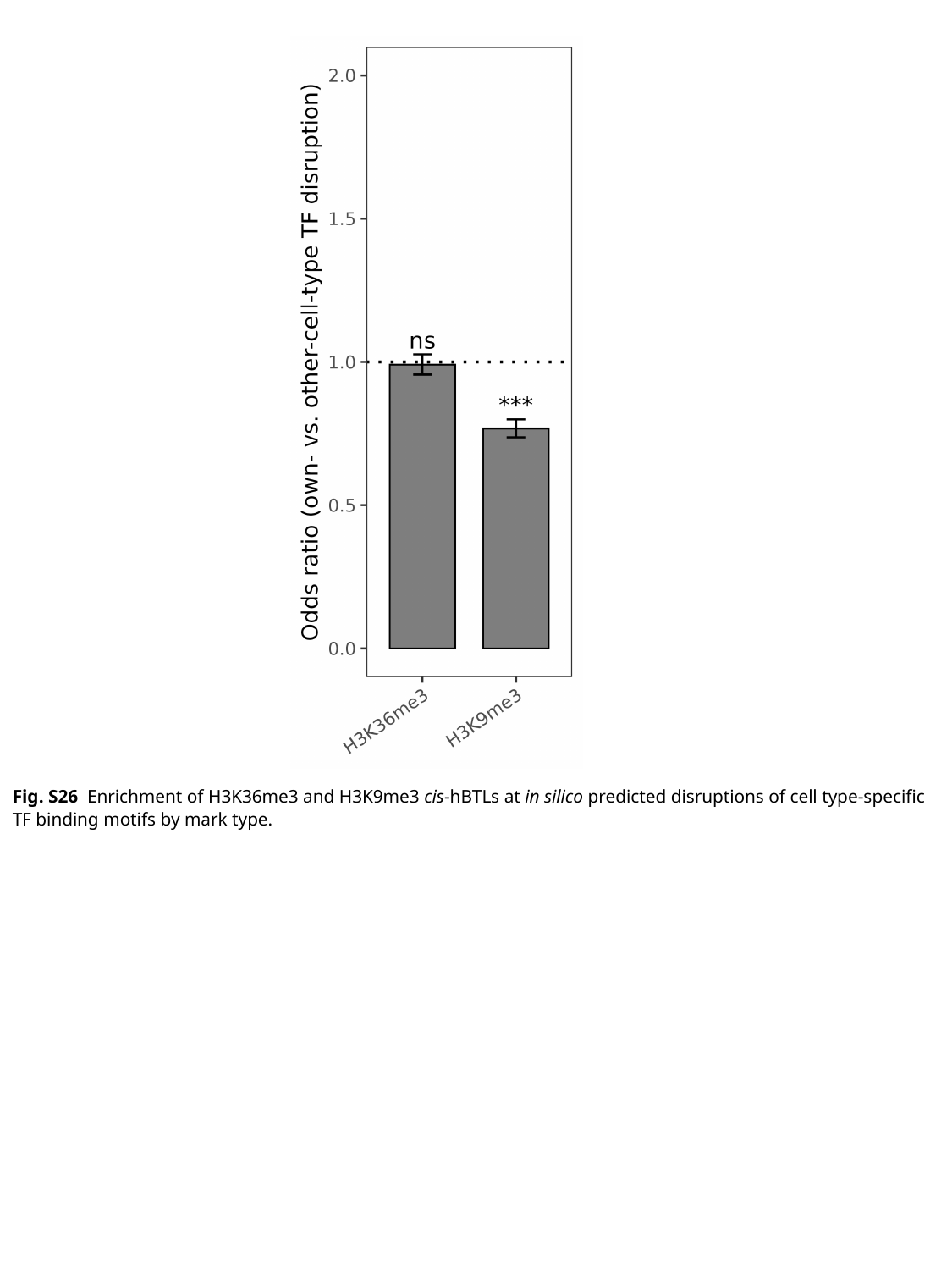

Fig. S26 Enrichment of H3K36me3 and H3K9me3 cis-hBTLs at in silico predicted disruptions of cell type-specific TF binding motifs by mark type.

### Slide 31
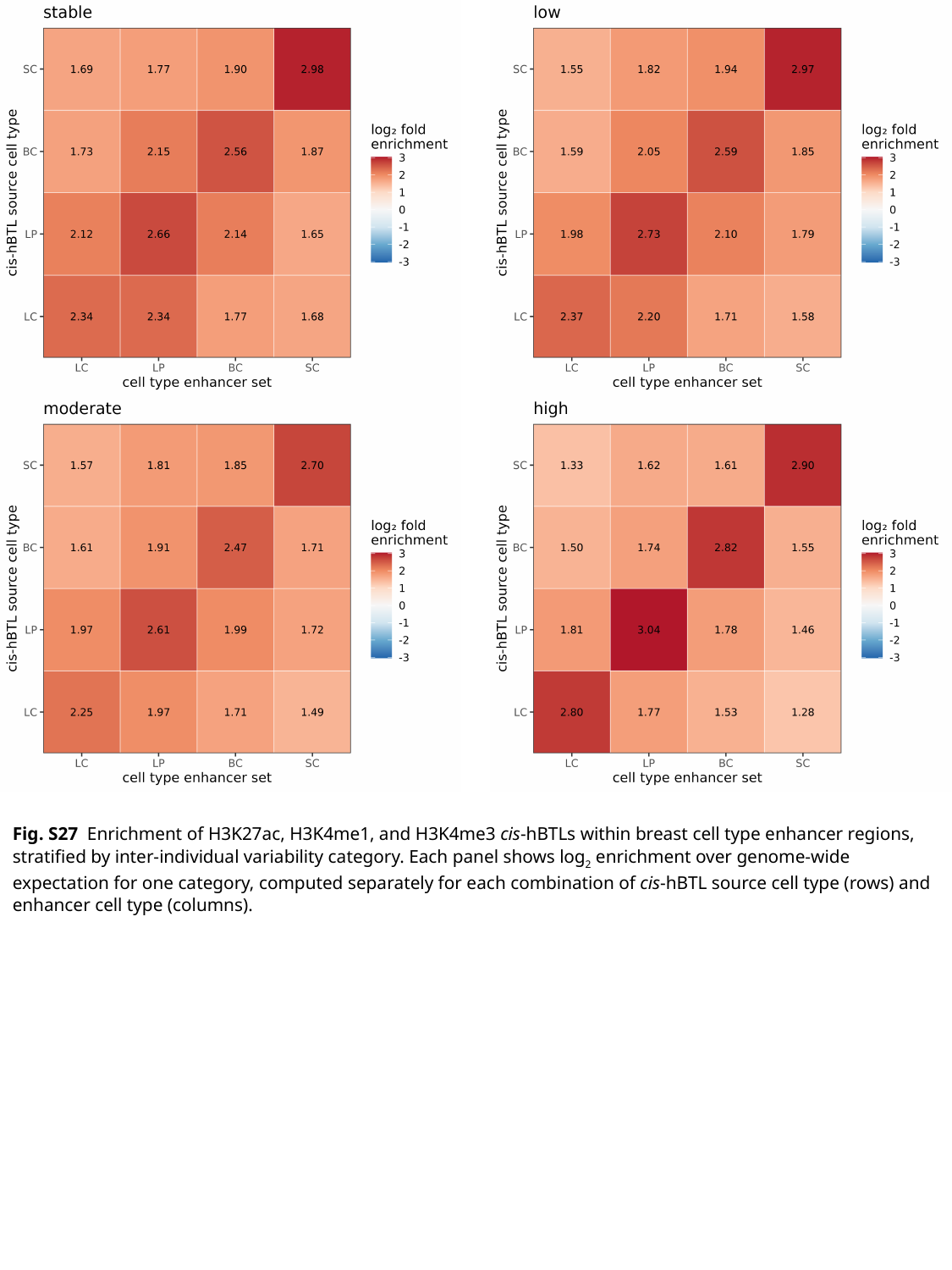

Fig. S27 Enrichment of H3K27ac, H3K4me1, and H3K4me3 cis-hBTLs within breast cell type enhancer regions, stratified by inter-individual variability category. Each panel shows log_2_ enrichment over genome-wide expectation for one category, computed separately for each combination of cis-hBTL source cell type (rows) and enhancer cell type (columns).

### Slide 32
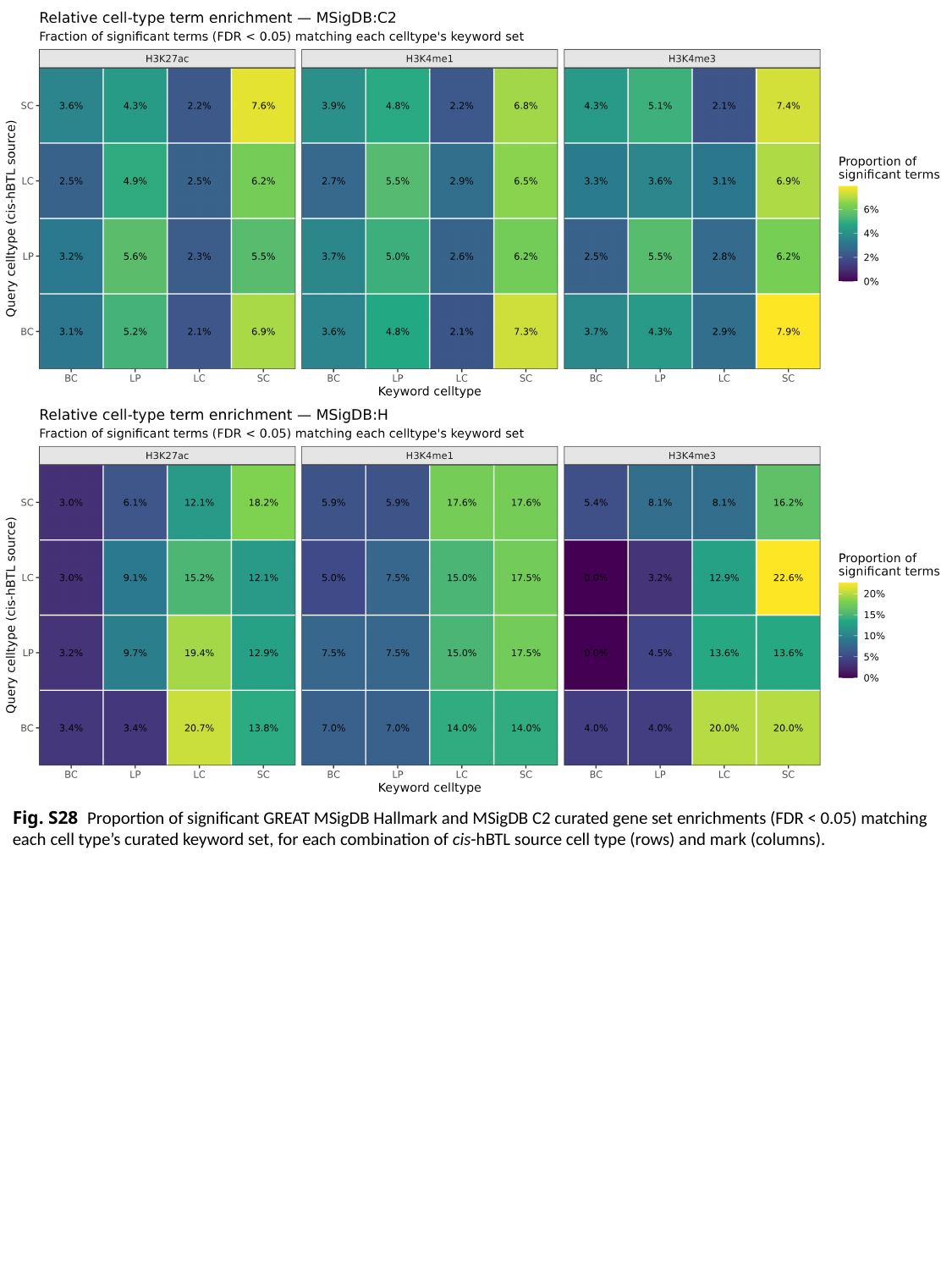

Fig. S28 Proportion of significant GREAT MSigDB Hallmark and MSigDB C2 curated gene set enrichments (FDR < 0.05) matching each cell type’s curated keyword set, for each combination of cis-hBTL source cell type (rows) and mark (columns).

### Slide 33
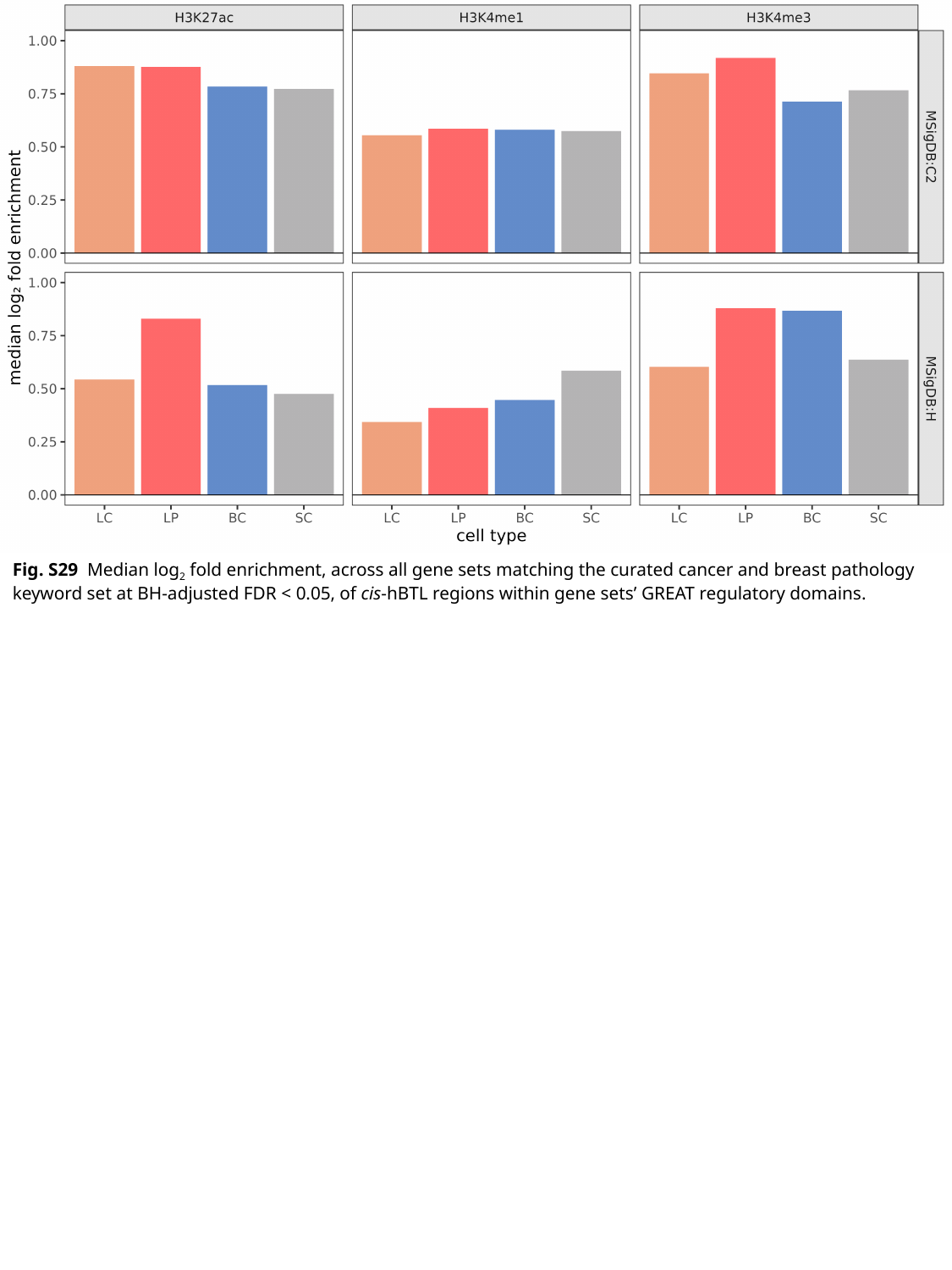

Fig. S29 Median log_2_ fold enrichment, across all gene sets matching the curated cancer and breast pathology keyword set at BH-adjusted FDR < 0.05, of cis-hBTL regions within gene sets’ GREAT regulatory domains.

### Slide 34
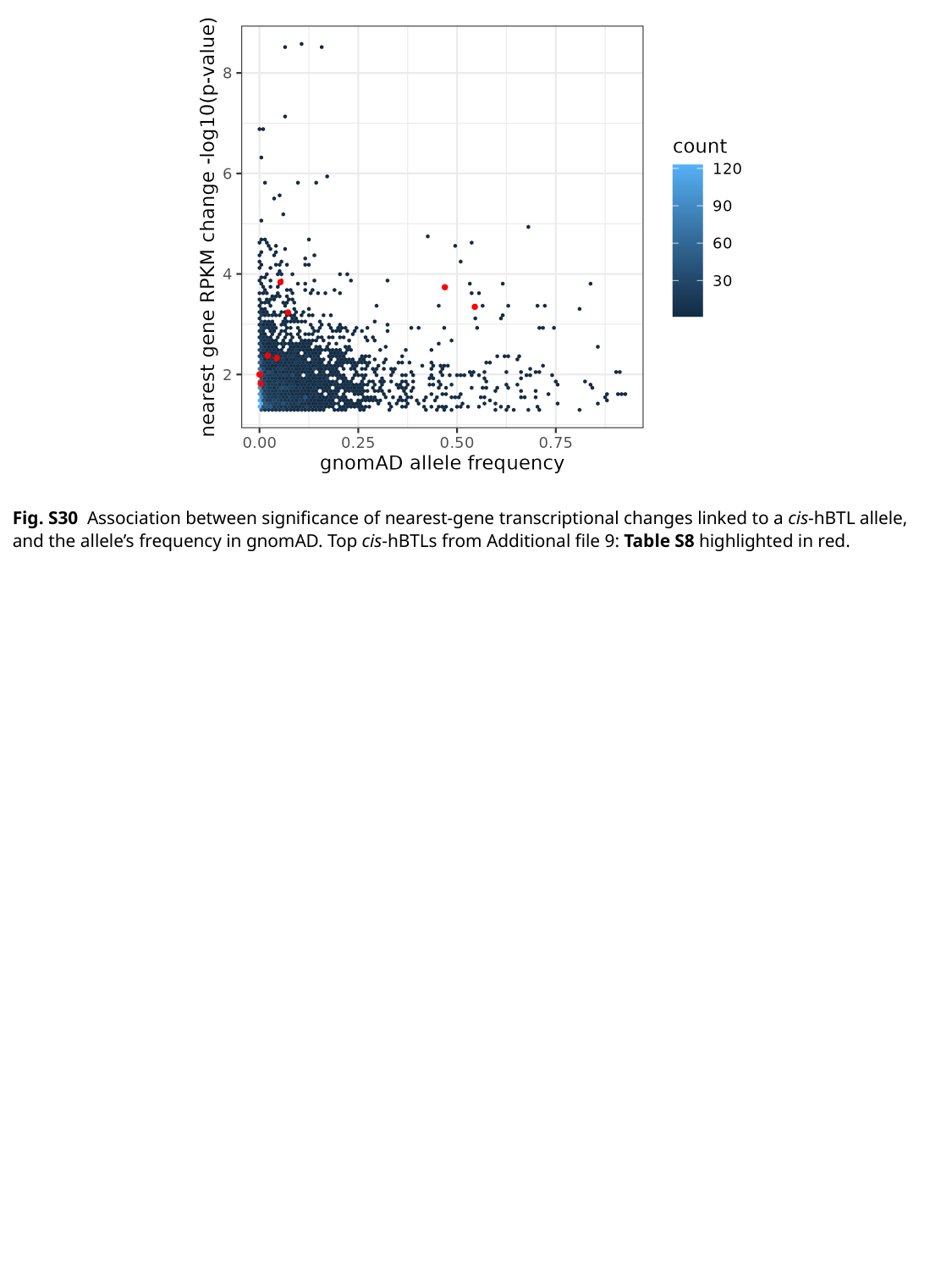

Fig. S30 Association between significance of nearest-gene transcriptional changes linked to a cis-hBTL allele, and the allele’s frequency in gnomAD. Top cis-hBTLs from Additional file 9: Table S8 highlighted in red.

### Slide 35
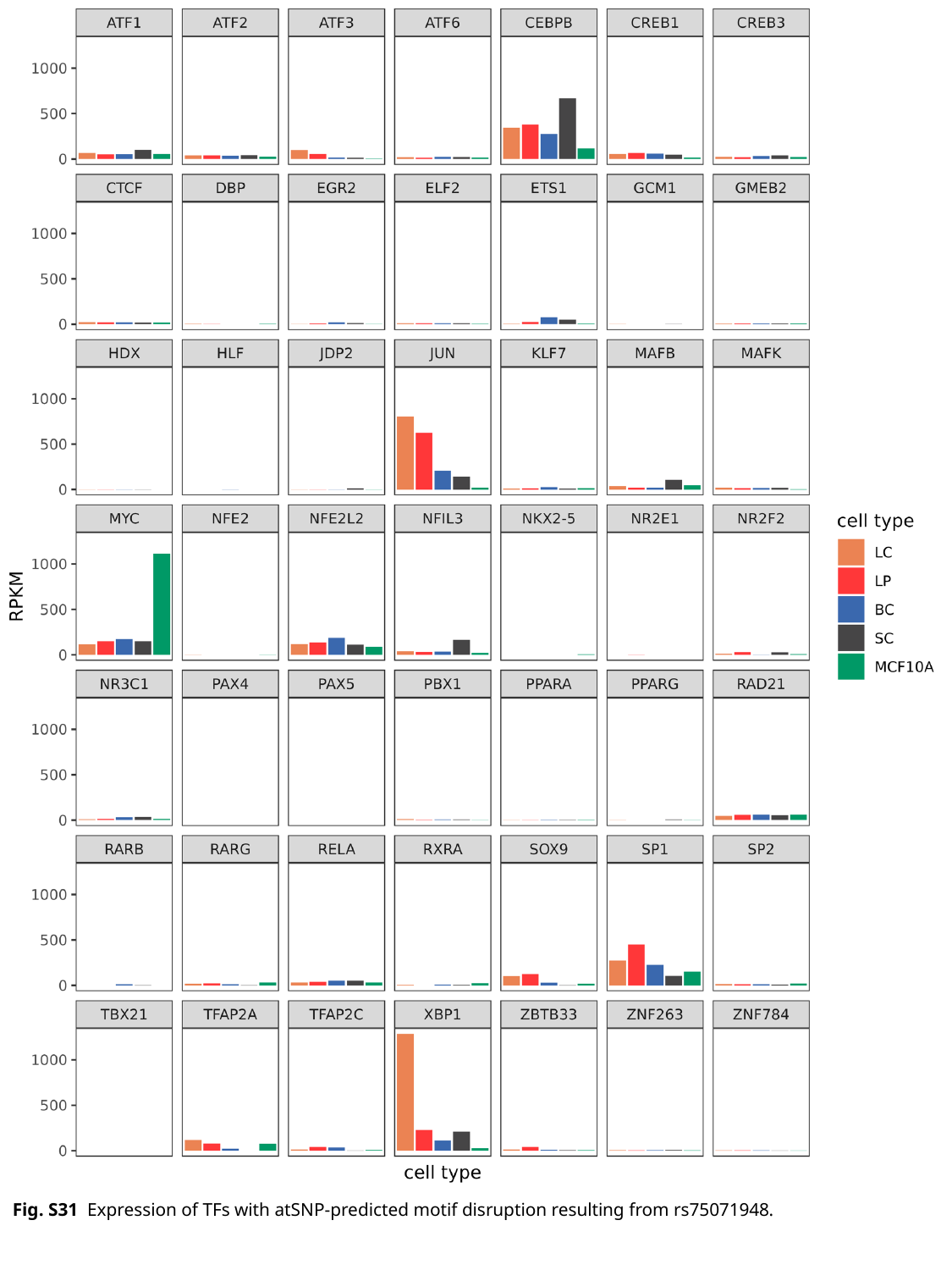

Fig. S31 Expression of TFs with atSNP-predicted motif disruption resulting from rs75071948.

### Slide 36
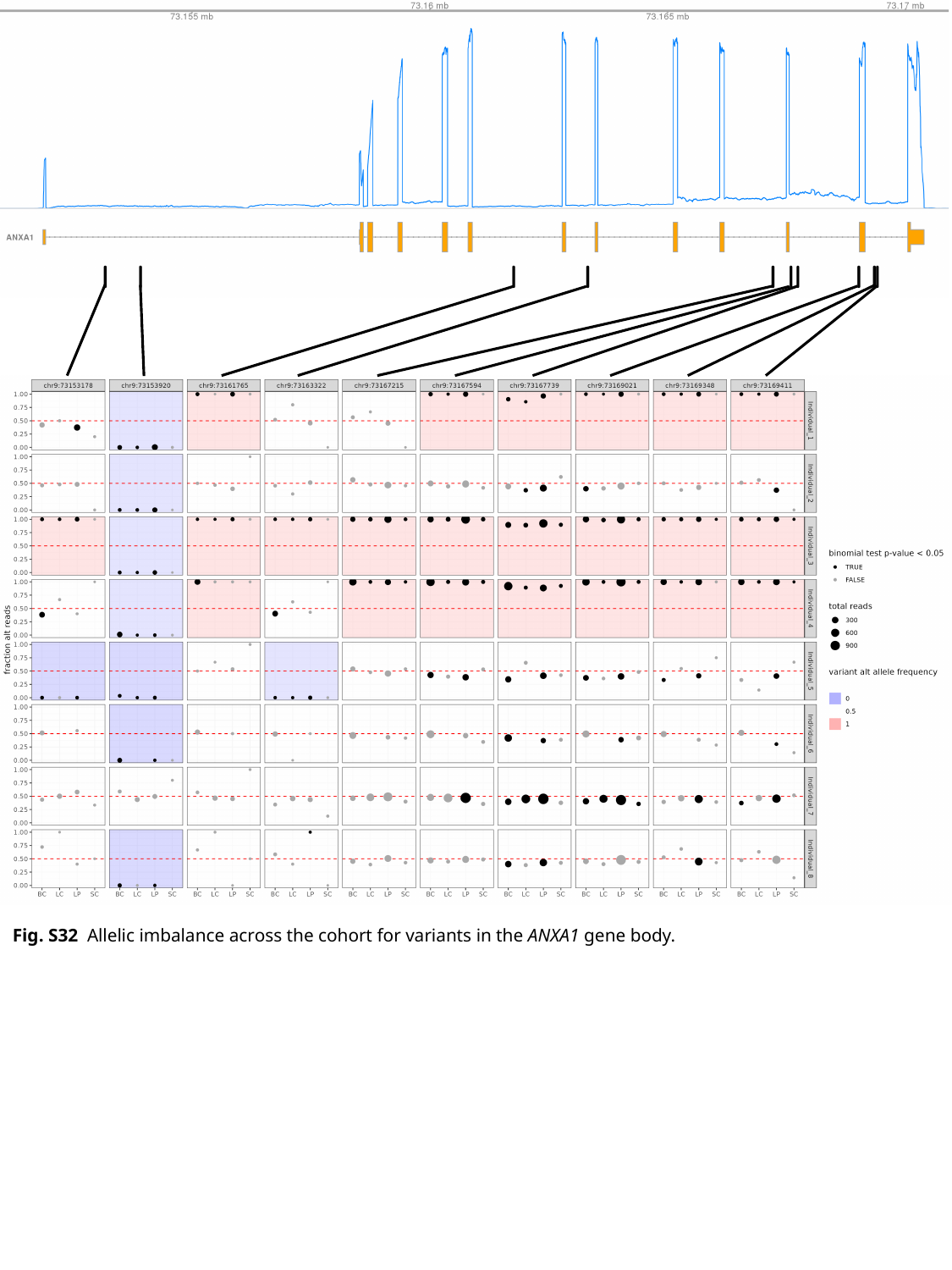

Fig. S32 Allelic imbalance across the cohort for variants in the ANXA1 gene body.

### Slide 37
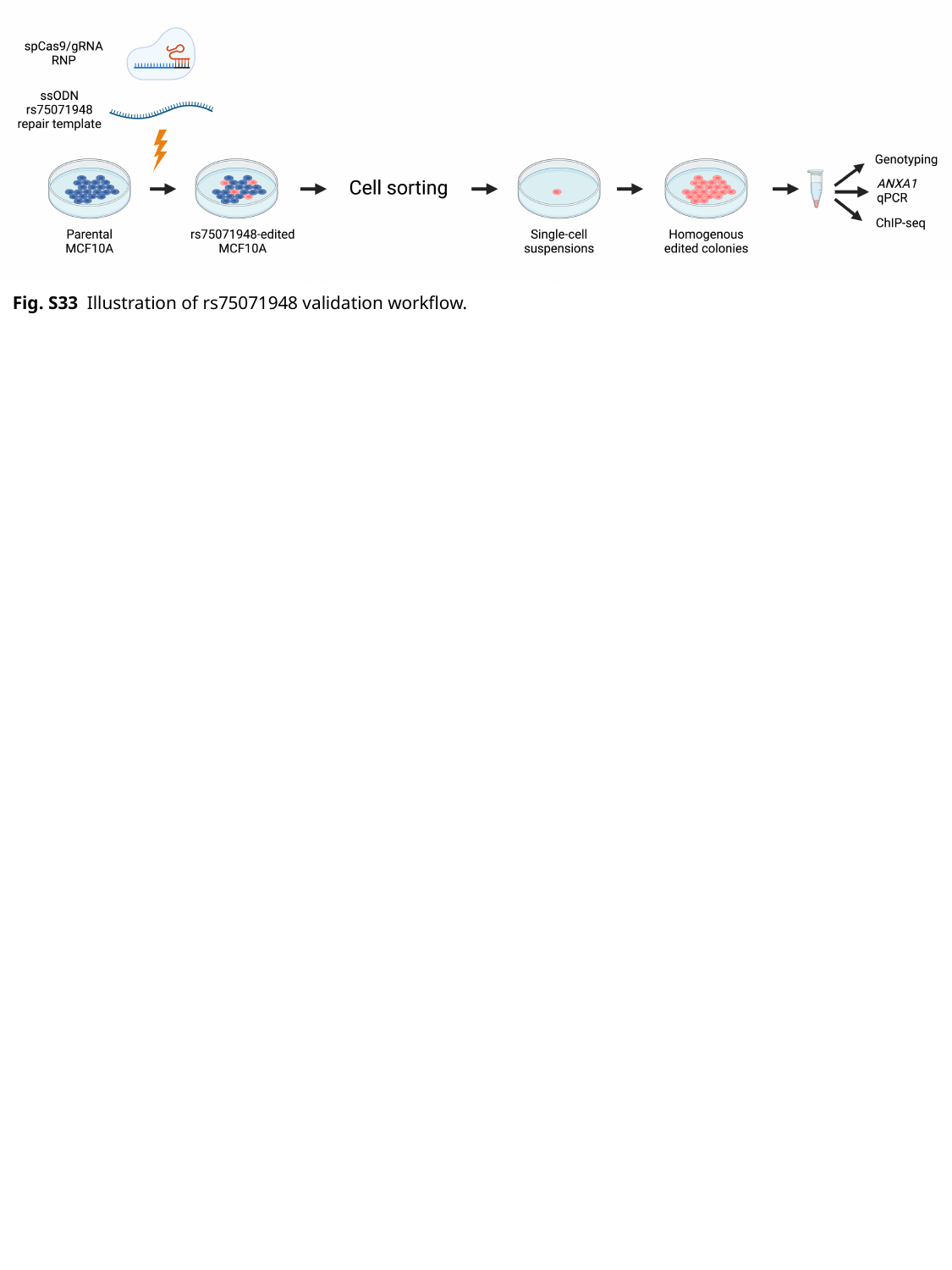

Fig. S33 Illustration of rs75071948 validation workflow.
